# Biosynthesis of plant papanridins -A group of novel oligomeric flavonoids

**DOI:** 10.1101/2023.03.01.530648

**Authors:** Yue Zhu, Seyit Yuzuak, Xiaoyan Sun, De-Yu Xie

**Affiliations:** Department of Plant and Microbial Biology, North Carolina State University, Raleigh, NC, USA; Department of Chemistry, North Carolina State University, Raleigh, NC, USA; Department of Molecular Biology & Genetic, Mehmet Akif Ersoy University, Burdur, Turkey

## Abstract

Discovery of novel flavonoids and their biosynthesis are fundamental to understand their roles in plants and benefits to human and animal health. Herein, we report a new polymerization pathway of a group of novel oligomeric flavonoids in plants. We have engineered red cells for discovering genes of interest involved in the flavonoid pathway and identified a gene that encodes a novel flavanol polymerase (FP) localized in the central vacuole. FP catalyzes the polymerization of flavanols, such as epicatechin and catechin, to produce yellowish dimers or oligomers. Structural elucidation show that these compounds are featured with a novel oligomeric flaven-flavan (FF) skeleton linked by interflavan-flaven and interflaven bonds, which are different from proanthocyanidins and dehydrodicatechins. Detailed chemical and physical characterizations further demonstrate that FFs are novel flavonoids. Mechanistic investigations show that FP polymerizes flavan-3-ols and flav-2-en-3-ol carbocation to form dimeric or oligomeric flaven-4→8-flavans, termed as papanridins. Data from transgenic, mutation, metabolic profiling, and phylogenetic analyses demonstrate that the biosynthesis of papanridins is prevalent in cacao, grape, blue berry, corn, rice, Arabidopsis and others in the plant kingdom. Given that these findings are the first report, many questions remain for answers. For instance, what are roles of papanridins in plants and what benefits do they have for human and animal health? We anticipate that these findings will promote investigations across plant, nutritional, and animal sciences to understand papanridins in plants and food products.

**Teaser:** Plant flavanol polymerase catalyzes the biosynthesis of novel oligomeric flavonoids in the plant kingdom.

## Introduction

Flavonoids are a large group of plant natural products [1–3] and play important roles in plant adaptation, pollination, and reproduction during the evolution [4–7]. Anthocyanin pigments play key roles to attract pollinators and seed dispensers for reproduction of plants [8]. Flavanols and proanthocyanidins (PAs) protect plants against diseases caused by pathogens and damages caused by UV-radiation [9]. Moreover, flavonoids are fundamental nutrients that are daily consumed by humans [10]. In general, they are strong antioxidants and have antiviral, anti-cardiovascular, anti-cancer, and anti-aging functions [11–16]. Recently, we reported that flavanol-gallates and dimeric PAs had a promising activity to inhibit SARS-Cov-2 and showed a potential to treat COVID-19 [17, 18]. These benefits indicate that the continuous discovery of new and novel flavonoids is important to innovate new medicines and nutraceuticals for human and animal health. To date, approximately 13,000 flavonoids have been reported from the plant kingdom [19]. Based on their structures and biogenesis from phenylalanine or tyrosine, flavonoids are classified to chalcones, flavanones, flavones, aurones, dihydroflavonols, flavonols, flavan-3, 4-diols, anthocyanins, flavanols, proanthocyanidins (PAs, condensed tannins), isoflavonoids, bioflavonoids, and neoflavonoids (Fig. 1). Of these groups, PAs are the only group of oligomeric or polymeric flavonoids that are biosynthesized from flavanols [3, 20]. In addition, dimeric or oligomeric catechins formed by the catalysis of peroxidases or polyphenol oxidases, e.g. yellowish dehydrodicatechin A, occur during the process of wine production and fermentation of green tea leaves [21–24]. To date, the biosynthesis of flavonoids has gained intensive studies. Pathways to each group have been biochemically, genetically, and molecularly elucidated in different plants. The main pathway includes beginning (from phenylalanine to coumaryl-CoA), early (coumaryl-CoA to dihydroflavonols), and late steps (dihydroflavonols to flavonols, anthocyanins, flavanols, and PAs) (Fig. 1) [8]. Sixteen main pathway genes identified include phenylalanine ammonia-lyase (*PAL*), coumarate 4-hydroyxlase (*C4H*), 4 coumarate-CoA ligase (*4CL*), chalcone synthase (*CHS*), chalcone isomerase (CHI), flavonone 3-hydroxylase (*F3H*), isoflavone synthase (*IFS*), flavone synthase (*FNS*), flavonoid 3’hydroxylase (*F3’H*), flavonoid 3’5’ hydroxylase (*F3’5’H*), dihydroflavonol reductase (*DFR*), flavonol synthase (*FLS*), leucoanthocyanidin reductase (*LAR*), anthocyanidin synthase (*ANS*), leucoanthocyanidin dioxygenase (*LDOX*), and anthocyanidin reductase (*ANR*) (Fig. 1) [9, 25–28]. Despite of the documented knowledge, many questions remain for answers. For example, how plants polymerize flavanols to PAs remain open for further investigation. PAs are oligomers or polymers of flavanols (flavan-3-ols) polymerized via an interflavan bond formed between the C_4_ of an upper unit and the C_8_ or C_6_ of a lower unit [20] (Fig. 1). Our previous hypotheses were that polyphenol oxidases, laccases, or peroxidases might be responsible for the catalysis of this step [20]. We proposed that these enzymes might oxidize flavanols to quinone methides to form flavanol carbocation, which was attacked by flavanols to form PAs (Fig. 1) [20]. An oxidation evidence was that a laccase-like enzyme encoded by *TT10* was reported to oxidize epicatechin and PAs to yellowish and browning compounds associated with the brownish seed coat color of Arabidopsis [29]. In addition, although evidence is limited to demonstrate these hypotheses, we recently proved the presence of flavanol carbocation involved in the formation of PAs in plant cells [30]. Moreover, recent reports revealed the complexity of polymerization of PAs (Fig. 1) [31]. A new function of LAR was characterized to catalyze the conversion of beta-cysteinyl epicatechin to epicatechin, which was a non-oxidoreduction catalysis associating with the formation of non-extractable PAs [32]. LDOX was shown to catalyze the epimerization of catechin to epicatechin via flav-2-en-3-ol [33]. Although ANR was demonstrated to catalyze the reduction of anthocyanidins to flavonols via proposed flav-2-en-3-ol, flav-3-en-3-ol, and flavan-3-one [34, 35], a recent follow-up study revealed that ANR could also converted flav-3-en-3, 4-diol to 2S, 3S, 4S-leucoanthocyanidins [36]. These new discoveries and the multiple roles of flavanols indicate that continuous efforts are necessary to understand the polymerization of PAs. Meanwhile, numerous questions remain for answers. Are there novel skeletons of flavonoids in the plant kingdom? Are flavanols, anthocyanins, flavonols, and flavones the end of each branch? In addition to PAs, can flavanols be precursors of other unknown structures? All these questions remain for investigations.

**Figure 1.**
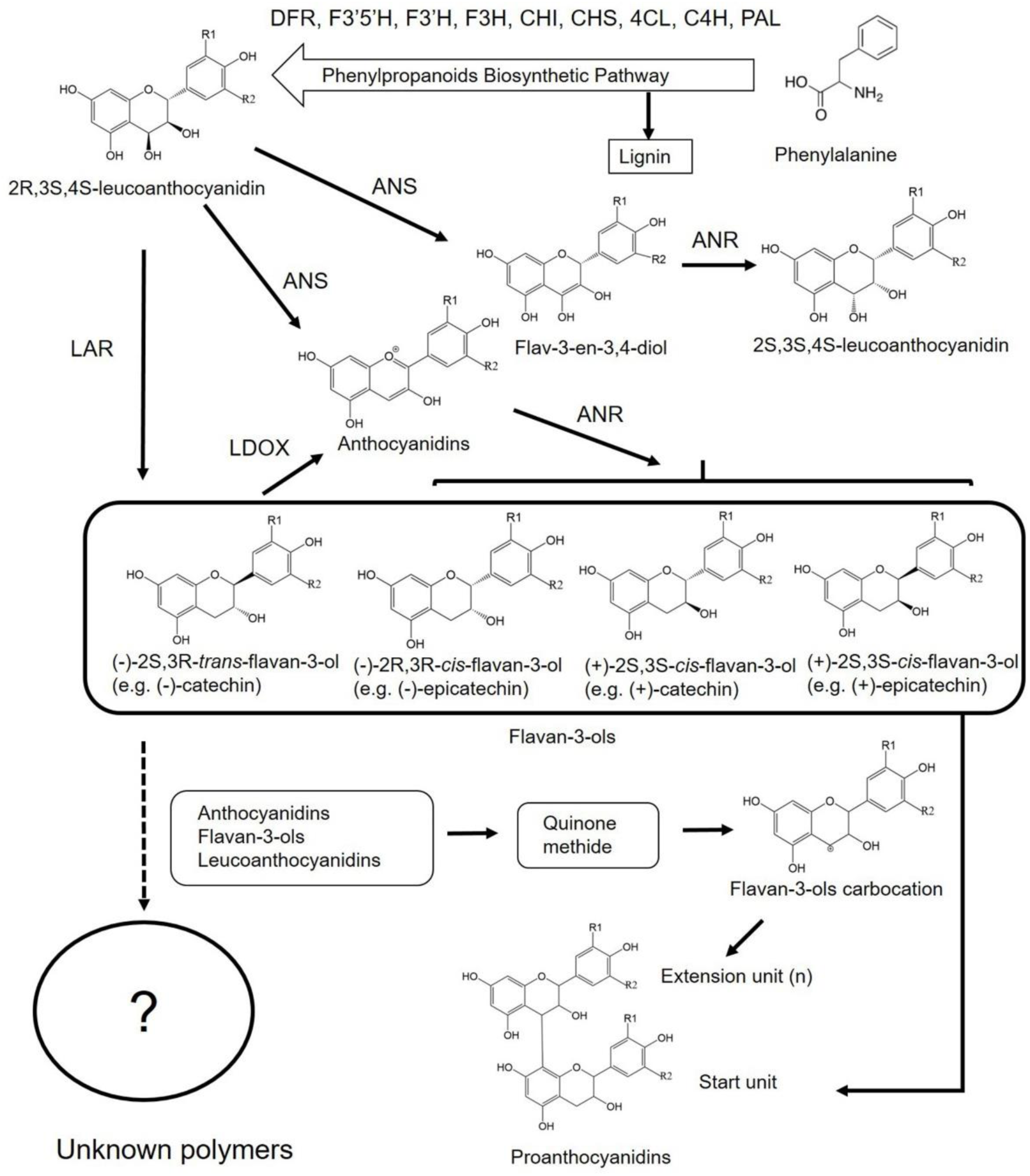
Biosynthesis of flavanols, proanthocyanidins, and other main flavonoid groups starting with phenylalanine. Proanthocyanidins are the only group of oligomeric or polymeric flavonoids from flavanols linked by an interflavan bond. The question mark means whether or not flavanols can form other oligomers or polymers. Abbreviation, PAL: phenylalanine ammonia-lyase, C4H: coumarate 4-hydroyxlase, 4CL: 4 coumarate-CoA ligase, CHS: chalcone synthase, CHI: chalcone isomerase, F3H: flavonoid 3-hydroxylase, F3’H: flavonone 3’hydroxylase, F3’5’H: flavonone 3’5’ hydroxylase, DFR: dihydroflavonol reductase, LAR: leucoanthocyanidin reductase, ANS: anthocyanidin synthase, LDOX: leucoanthocyanidin dioxygenase, ANR: anthocyanidin reductase, IFS: isoflavone synthase, FNS: flavone synthase, and FLS: flavonol synthase.

Although the direct extraction of phytochemicals from wild type plants is an effective method to discover new flavonoid compounds, metabolic engineering of plants shows a power to discover new compounds and biosynthetic pathways hidden in plants in natural conditions. *Arabidopsis thaliana* mainly produces two anthocyanins in nature [37]. However, our studies showed that the dominant *pap1-D* mutants highly expressing the *Production of Anthocyanin Pigment* 1 (*PAP1*) gene (encoding MYB75, a R2R3-MYB transcription factor) could produce 27 new anthocyanins of *A. thaliana* [38]. In addition, we have shown that the overexpression of *PAP1* led to a massive transcriptional and metabolic programing to produce new anthocyanins in tobacco plants [39–42]. In this study, we report to use our engineered PAP1 and PAP1-BAN (or ANR) plants for discovering novel flavonoids and genes of interest. PAP1 tobacco was transcriptionally programmed by the overexpression of *PAP1* [20] and named it as *N. tabacum* Xanthi PAP1. This transgenic genotype highly biosynthesizes anthocyanins in all tissues. *BAN* encodes ANR that is a key reductase catalyzing the biosynthesis of three stereo configurations of flavanols and PAs (Fig. 1) [20, 34, 43]. We have crossed PAP1 and ANR tobacco plants and created a PAP1-ANR (BAN) genotype that can produce flavanols and PAs in red cells of leaves and flowers [44]. However, although PAP1-ANR tobacco produces high contents of anthocyanin and appropriate levels of flavanols, the production of PAs is relatively low. Metabolic profiling indicated that PAP1-ANR plants produced new compounds compared to wild type ones. Based on these phytochemical phenotypes, we hypothesized that PAP1 and PAP1-ANR tobacco plants might express unknown pathways leading to the biosynthesis of new flavanol-derived metabolites. In addition, based on the data that PAs were only produced in red cells [34], we further hypothesized that red cells expressed enzymes toward the biosynthesis of PAs and unknown metabolites. To test these hypotheses, we engineered red 6R and control P3 cells from PAP1 and wild-type tobacco plants, respectively [40]. Red 6R cells produced anthocyanins, while the control P3 cells did not. These cell lines were used to identify new enzymes and genes of interest. A protein was demonstrated to be a flavanol polymerase (FP). A cDNA encoding this FP was cloned from red 6R cells. Transgenic, phytochemical, MS/MS, NMR, and chemical and physical characterization demonstrated that FP catalyzed the polymerization of flavanols to produce a group of novel yellowish dimeric and oligomeric flavonoids. Given that FP was discovered in red PAP1 cells and novel flavonoids were formed in <u>PA</u>P1-<u>ANR</u> tobacco plants, we named this group of novel flavonoids as papanridins. Phylogenetic and metabolic profiling analyses have indicated that the biosynthesis of papanridins is prevalent in the plant kingdom.

## Results

### Yellowish dimeric PA-like compounds formed by the catalysis of a flavanol polymerase identified from red 6R cells

Red 6R cells (Supplementary Fig. S1a) were engineered from our PAP1 tobacco variety that was programed to highly produce anthocyanins in all tissues by the overexpression of *PAP1* [40, 44]. In addition, White P3 cells were developed from wild type plant as a control. Based on our previous PA engineering in PAP1 tobacco plants [44], we hypothesized that red cells expressed genes encoding enzymes that catalyze polymerization of flavanols to dimeric or oligomeric PAs. Red 6R and white P3 cells were used to develop cell suspension cultures (Fig. 2a and Supplementary Fig. S1b). Both suspension cells and liquid media (supplementary Fig. S1) were collected to isolate crude proteins (Fig. 2b and supplementary Fig. S2a), which were tested for a polymerase activity with epicatechin and catechin as substrates. The crude protein extracts from both cell lines converted epicatechin and catechin to yellowish compounds (Supplementary Fig. S2b-d). In addition, proteins were boiled for 1-10 min as controls. An unexpected result was observed that after boiling for 1-5 min, boiled crude proteins also had an activity although compared with that of un-boiled crude enzymes, the intensity of yellowish color was reduced. SDS-PAGE analysis showed the presence of multiple protein bands from the boiled crude protein extracts of both 6R and P3 cells (Fig. 2b and Supplementary Fig. S2a). Further TLC analysis visualized with DMACA staining showed that un-boiled and 1-3 min boiled proteins converted epicatechin and catechin to dimeric PA-like compounds (Fig. 2c and Supplementary Fig. S2e). To understand whether these compounds resulted from a real enzymatic catalysis, crude, boiled, and proteinase K (PK)-digested proteins were separated on native gels. Then, the gels were placed in the pH7 Tris-HCl buffer supplemented with epicatechin. The four lanes with crude, 3-min and 6-min boiled, and PK-digested 6R proteins showed yellowish to deep yellowish coloration (Supplementary Fig. S2f). The three lanes with crude, 3-min boiled, and PK-digested P3 proteins showed slightly yellowish coloration, while the lane with 6-min boiled P3 proteins did not show activities. In addition, the lane with BSA control did not show yellowish coloration. These results not only indicated that both crude and 3-6 min boiled 6R proteins contained active enzymes, but also revealed that the red 6R protein extracts had different active enzymes from the P3 protein extracts (Supplementary Fig. S2f).

**Fig. 2.**
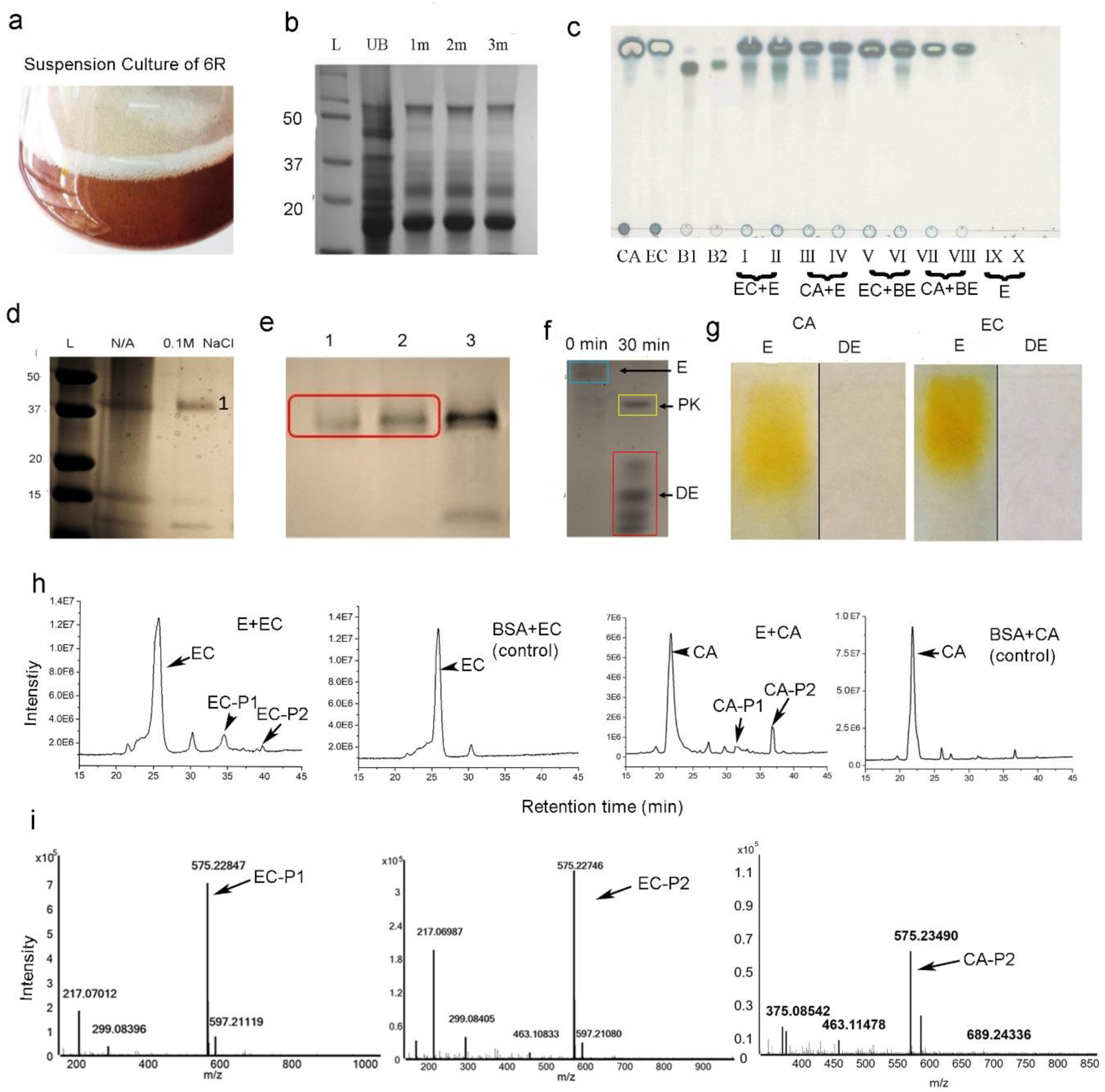
A flavonol polymerase isolated from engineered red tobacco cells. **a,** a suspension culture of red 6R cells was developed to produce proteins. **b,** a SDS-PAGE gel image shows band profiles of crude protein extracts from red 6R cells. UB: un-boiled protein and 1m, 2m, and 3m: boiled for 1, 2, and 3 min. **c,** a TLC image shows dimeric PA-like compounds produced from incubations consisting of crude proteins and catechin or epicatechin. Abbreviations, CA: catechin; EC: epicatechin; B1 and B2: procyanidin B1 and B2; EC+E (I+II): two repeated reactions (I and II) of epicatechin (EC) and un-boiled enzyme extracts (E); CA+E (III and IV): two repeated reactions of catechin (CA) and un-boiled enzyme extracts; EC+BE (V and VI): two repeated reactions of epicatechin and 3-min boiled enzyme (BE) extracts; CA+BE (VII and VIII): two repeated reactions of catechin and 3-min boiled enzyme extracts; E (IX and X): two enzyme extract controls without adding substrates. **d,** a SDS-PAGE image shows protein bands separated from a DEAE Sephadex A-20-120 column with elution buffer supplemented with 0.1 M NaCl. N/A: buffer without adding NaCl. **e,** a SDS-PAGE image shows protein bands of three fractions (1-3) isolated from a Sephadex G-75 column. **f,** a SDS-PAGE image shows the purified protein without a digestion of proteinase K (PK) (0 min) and its peptides resulted from the digestion of PK for 30 min (30 min). E: enzyme and DE: digested enzyme by PK. **g,** native PAGE images show that the enzyme purified from the Sephadex G-75 column (f) catalyzes catechin and epicatechin to yellowish compounds on the gels but PK-digested enzyme (f) cannot. CA: catechin, EC: epicatechin, E: enzyme, and DE: digested enzyme. **h,** total ion chromatograms (TICs) show that the purified enzyme converts epicatechin (E+EC) to two new products (EC-P1 and EC-P2) and catechin (CA+E) to two new products (CA-P1 and CA-P2), but BSA control does not catalyze epicatechin and catechin (BSA+EC and BSA+CA). CA: catechin, EC: epicatechin, EC-P1 and EC-P2: epicatechin products 1 and 2, CA-P1 and CA-P2: catechin products 1 and 2. **i,** the mass spectra characterize that EP-P1, EP-P2, and CA-P2 have the same value of mass to charge ratio, 575 [m/z]^-^.

Next, we used three steps, protein boiling, ion-exchange column, and Sephadex columns, to purify the active enzyme from 6R cells (Fig. 2d-e), which was examined with native gels for activity assay. *In situ* reactions on native gels showed that the purified protein converted catechin and epicatechin to yellowish compounds, while proteinase K-digested protein did not (Fig. 2f-g and Supplementary Fig. S3). LC-MS analysis detected that the enzymatic reactions converted epicatechin or catechin to two main new compounds, which were not produced from negative control incubations (Fig. 2h). These compounds from epicatechin and catechin were labeled as EC-P1, EC-P2, CA-P1 and CA-P2, which had different retention times, respectively. Further MS analysis showed that all the four compounds had the same mass-to-charge ratio value, 575.23 [m/z]^-^. This value indicates that their molecular weight is 576 Dalton, the same as that of procyanidin A1 and A2 but two protons less than dimeric procyanidins (578 Dalton), suggesting that they are not dimeric B-type PAs. Given that like flavan-3-ols and PAs, these new yellowish compounds react with DMACA to form bluish coloration, they are dimeric PA-like compounds. Based on this dimerization activity, we named the purified protein as <u>F</u>lavanol <u>P</u>olymerase (FP).

### Gene cloning and catalytic activity of recombinant FP

We used two steps to clone a cDNA encoding FP. First, the purified FP protein was sequenced to obtain seven peptides, which were used for blast search at GenBank (Supplementary Fig. S4). The blasting results showed that all peptides hit one single tobacco peroxidase (Supplementary Fig. S4). Second, we designed a pair of primers to clone approximately 1 kb cDNA from red 6R cells. Meanwhile, RT-PCR analysis revealed that it was not expressed in leaves, stems, roots of tobacco plants and control white 6W cells (Fig. 3a). A sequence alignment and modeling with deduced amino acids showed that it belonged to a member of the class III of the peroxidase family I (Supplementary Fig. S5a), which was featured with four conserved disulfide bridges and two calcium ions [45, 46]. In addition, the N-end has a secretory signal peptide. To understand its subcellular localization, we fused FP to the C-end of GFP and generated transgenic PAP1-BAN(ANR)-NtFP(FP) tobacco plants. A plasmolysis and confocal microscopic examination of transgenic tobacco roots showed that FP was localized in the central vacuole (Supplementary Fig. S6a-b).

**Fig. 3.**
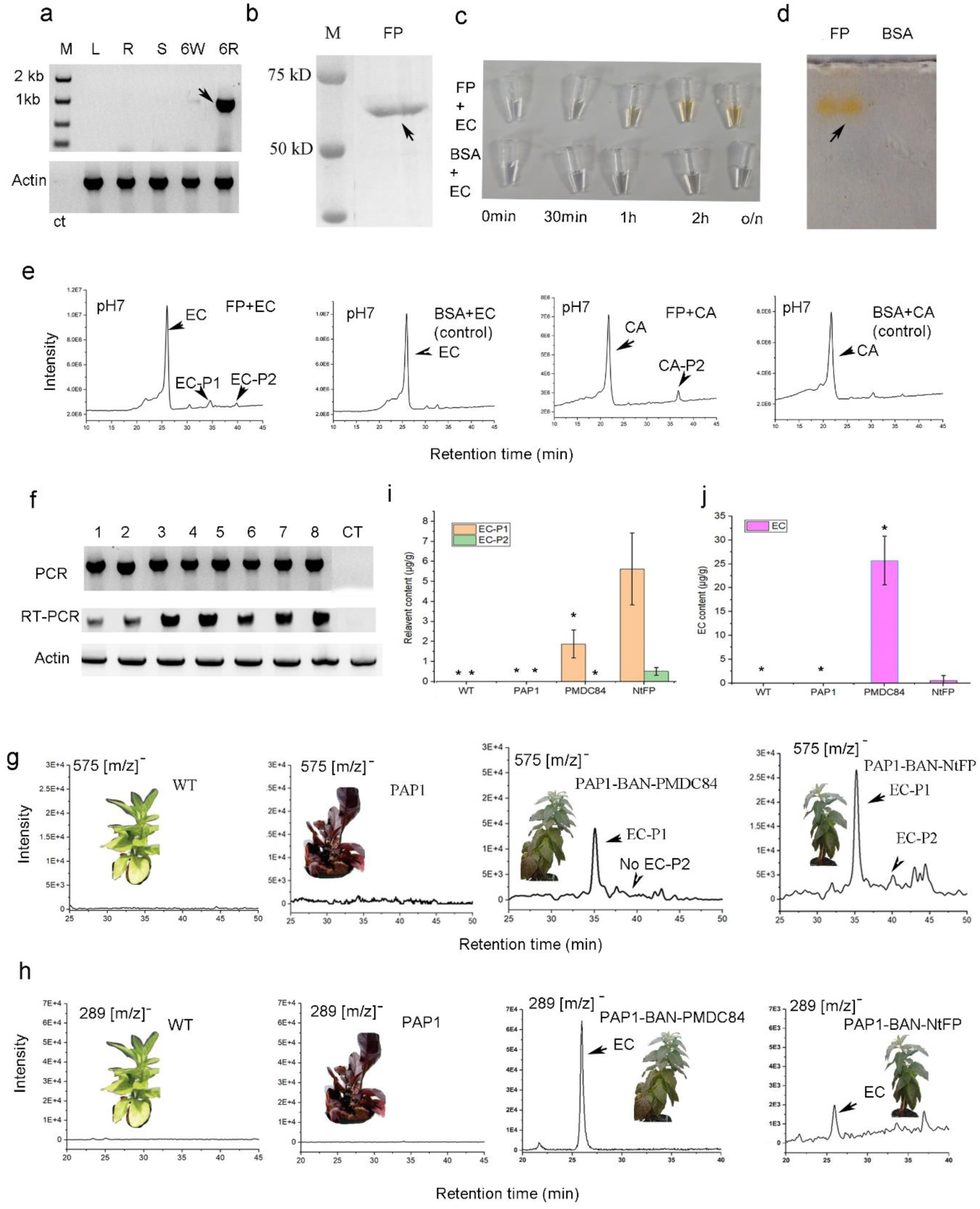
Cloning of flavanol polymerase (*FP*) cDNA and functional analysis of recombinant FP. **a,** a cDNA of *FP* was amplified from red 6R cells by RT-PCR. M, DNA ladder, L: leaves, R: roots, S: stems, and 6W: a white cell line engineered together with red 6R cells from the same explant. **b,** a SDS-PAGE gel image shows the recombinant GST-FP-His protein purified. **c,** the GST-FP-His (FP) protein catalyzed epicatechin (EC) to yellowish compounds, but BSA control did not. The incubation was composed of FP or BSA and epicatechin in 50 mM pH 7 Tris-HCl buffer that was oxygenated by a strong vortex. 0min, 30min, 1h, 2h, and o/n: reaction 0 and 30 min, 1 and 2 hrs, and overnight at room temperature. **d,** an image shows that the purified GST-FP-His protein on the native-PAGE gel converted epicatechin to yellowish compound(s), but the BSA control did not, after the gels were incubated with epicatechin for 30 min (Gels were placed in 50 mM pH 7 Tris-HCl supplemented with 1.0 µg/µl epicatechin contained in petri dishes, which were shaken on a rotary shaker). **e,** TIC profiles of 30-min reaction products revealed that GST-FP-His (E) converted epicatechin to two products (EC-P1 and EC-P2) and catechin (CA) to one main product (CA-P2), but the BSA control did not have these activities. **f,** PCR and RT-PCR demonstrated the integration and expression of the *FP* transgene in *FP* transgenic (1-8) but not in PMDC84 (vector control) transgenic PAP1-ANR plants (CT). **g,** elective ion chromatogram (EIC) analysis detected EC-P1 and EC-P2 from leaf extracts of PAP1-BAN-FP (NtFP) (*FP* transgenic PAP1-BAN plants) and EC-P2 from PAP1-BAN-MPDC84 (PMDC84 transgenic PAP1-BAN) plants, but did not detect these compounds from wild type and PAP1 tobacco plants. Plant picture inserts show their phenotypes. There were not phenotypic differences in PAP1-BAN-MPDC84 and PAP1-BAN-FP (NtFP) transgenic plants. **h,** EIC analysis detected the reduction of epicatechin (EC) in leaves of PAP1-BAN-FP (NtFP) plants compared to PAP1-BAN-MPDC84 ones. EC was not detected from wild type and PAP1 plants. Plant picture inserts show their phenotypes. **i,** the levels of both EC-P1 and EC-P2 were significantly higher in leaves of PAP1-BAN-FP (NtFP) plants than in those of PAP1-BAN-MPDC84 ones. These compounds were not detected from leaves of wild type and PAP1 plants. **j,** the content of epicatechin (EC) was significantly lower in leaves of PAP1-BAN-FP (NtFP) plants than in those of PAP1-BAN-MPDC84 ones. However, EC was not detected from leaves of wild type and PAP1 plants.

A recombinant FP was purified from its heterogeneous expression induced in *E. coli* (Fig. 3b and Supplementary Fig. S7). The incubation of the purified recombinant FP and epicatechin in the oxygenated buffer (via vortexing) produced yellowish compounds (Fig. 3c), supporting the results from the native FP (Fig. 2). Further *in situ* reaction on native gels demonstrated the catalytic activity of the recombinant FP (Fig. 3d). LC-MS analysis showed that the yellowish compounds were EC-P1 and EC-P2 from epicatechin and CA-P2 from catechin (Fig. 3e), which supported the results from the native FP analysis (Fig. 2h).

### Effects of pH, H_2_O_2_, NADPH, beta-mercaptoethanol, and vitamin C on FP activity and kinetics

The catalysis of recombinant FP was further characterized in different *in vitro* reaction conditions. First, its catalysis depended upon the presence of oxygen in the reactions. The oxygenation of the reaction buffers (pH6, pH7, and pH8) by either vortexing or adding H_2_O_2_ was essential for the FP’s catalysis to convert EC and CA to their corresponding products. When the reaction buffers were not oxygenated, the catalytic activity of FP was either undetectable or slightly detectable only (Supplementary Fig. S8a-d). Furthermore, the deoxygenation of buffers via dissolving nitrogen led to the complete loss of the catalysis of FP. Second, the effects of H_2_O_2_ on the catalytic activity of FP in pH6-7 buffers were different from that in pH8 buffer (Supplementary Fig. S8 a-d). In pH6 and pH7 buffers added with H_2_O_2_, FP quickly converted EC to brown-yellowish compounds (Supplementary Fig. S8a-d, tubes labeled as FP H_2_O_2_). TLC analysis detected PA-like compounds from the reactions (Supplementary Fig. S8b and d). LC-MS analysis showed that FP converted EC to EC-P1, EC-P2, and another 861 [m/z]^-^ compound (Supplementary Fig. S8e, labeled as FP+EC+H_2_O_2_) and converted catechin to CA-P1 and CA-P2 (Supplementary Fig. S9, labeled as FP+CA+H_2_O_2_). By contrast, FP in these two buffers without adding H_2_O_2_ did not convert EC to yellowish compounds (Supplementary Fig. S8a-d, tubes labeled as FP alone). In addition, in the BSA control reactions either in the presence or absence of H_2_O_2_, EC and catechin were not converted to their corresponding compounds (Supplementary Figs. S8a-e and S9). In pH8 buffer, enzymatic assays obtained different results. In the presence of H_2_O_2_, EC was converted to yellow-brownish compounds; however, neither could TLC nor LC-MS analysis detect PA-like compounds (Supplementary Fig. S8b and f). In the absence of H_2_O_2_, EC was converted to yellowish compounds (Supplementary Fig. S8a), from which LC-MS analysis detected EC-P1 (Supplementary Fig. S8f). In two types of negative control reactions by substituting FP with BSA (EC + H_2_O_2_ + BSA and EC+ BSA), EC was also converted to slightly yellowish compounds; however, neither could TLC nor LC-MS analysis detect PA-like compounds (Supplementary Fig. S8b and f). In addition to EC, LC-MS analysis could not detect CA-P1 and CA-P2 from catechin in all pH8 conditions (Supplementary Fig. S9). Based on these data, the yellowish coloration in the pH8 buffer likely resulted from the degradation of EC and CA, the products of which were water-soluble, thus could be hardly extracted by ethyl acetate for TLC and LC-MS analysis. Third, the velocity of the FP catalysis was linear during the incubation times from 30-120 min (Supplementary Fig. S10a). Forth, NADPH, beta-mercaptoethanol, and vitamin C inhibited the catalytic activity of FP (Supplementary Fig. S10 b). Fifth, the optimum pH and temperature for the reduction of EC was 8.0 and 50°C, respectively (Supplementary Fig. S10c-d). Sixth, the plot of velocity versus EC concentrations showed that FP was an allosteric enzyme (Supplementary Fig. S10e). Last, FP used catechol as a substrate but did not use phloroglucinol as a substrate (Supplementary Fig. S11).

### Overexpression of FP increases EC-P1, EC-P2, and PAs in PAP1-BAN tobacco

*FP* driven by two 35S promoters (Supplementary Fig. S6a) was transformed to PAP1-BAN tobacco plants to generate multiple transgenic PAP1-BAN-NtFP plants for metabolite analysis (Fig. 3f-h). Vector PMDC84 transgenic plants (PAP1-BAN-PMDC84) were also developed as controls. LC-MS analysis identified EC-P1 and EC-P2 from the leaf extracts of PAP1-BAN-NtFP plants, while only EC-P1 from the leaf extracts of PAP1-BAN-PMDC84 plants (Fig. 3g). However, neither compound was detected from PAP1 and wild type control plants. In comparison, the peak area of EC-P1 was much bigger in PAP1-BAN-NtFP extracts than in PAP1-BAN-PMDC84 extracts. Further estimation showed the significantly higher contents of these two compounds in PAP1-BAN-NtFP extracts than in control ones (Fig. 3i). By contrast, the contents of EC were significantly lower in PAP1-BAN-NtFP extracts than in PAP1-BAN-PMDC84 extracts (Fig. 3h and j). These data demonstrate that the transgenic FP converts EC into EC-P1 and EC-P2 *in planta*. LC-MS analysis also detected PB2, which was higher in PAP1-BAN-NtFP plants than in PAP1-BAN-PMDC84 ones in contents (Supplementary Fig. S12a-b). A further butanol-HCl boiling analysis showed that the content of total PAs was higher in PAP1-BAN-NtFP plants than in control ones (Supplementary Fig. S12c).

### EC-P1, EC-P2, CA-P1, and CA-P2 are novel dimeric flavonoids

We performed LC-qTOF-MS/MS analysis, phloroglucinolysis, and butanol-HCl boiling analyses to characterize EC-P1, EC-P2, CA-P1, and CA-P2. First, to annotate their potential structures, we used PB2 and PA2 as references for MS/MS analysis. Three characteristic fragmentation types of PB2 include heterocyclic ring fission (HRF), interflavan bond fission (IBF) between C4 and C8, and Retro-Diels-Alder (RDA) fission [47] (Fig. 4a-b). Skeleton based-main fragments of PB2 generated by collision-induced dissociation (CID) were 125 and 451 from HRF, 289 from IBF, and 407 from RDA (Fig. 4a-b). The main fragments derived from the CID of EC-P1 included 125 and 449 from HRF, 287 and 289 from IBF, and 407 from RDA (Fig. 4c). Another main fragment 271 was likely derived from the removal of one oxygen from the fragment of 287. The main fragments from PA2 included 125 and 449 from HRF, 285 (or 286) and 289 from IBF, and 423 from RDA (Fig. 4d). These main finger printings indicate that on the one hand, EC-P1, PA2, and PB2 have the same 125 fragment; on the other hand, EC-P1 is different from these two standards. Second, in the condition of methanol-HCl and phloroglucinol, the phloroglucinolysis of PB2 and other B-type PAs produces epicatechin-phloroglucinol (Fig. 4e) or catechin-phloroglucinol [30]. Our hypothesis was that if EC-P1 was a dimeric B-type PA molecule, it should undergo the phloroglucinolysis (Fig. 4g). We performed the phloroglucinolysis for PB2, PA2, and EC-P1. PB2 produced epicatechin-phloroglucinol with a 413 [m/z]^-^ (Fig. 4f), which was observed in PB2 control without phloroglucinolysis (Supplementary Fig. S13a). However, EC-P1 did not occur the phloroglucinolysis to produce epicatechin-phloroglucinol (Fig. 4h), which was not detected in purified EC-P1 either (Supplementary Fig. 13b). In addition, CA-P1 and PA2 did not occur the phloroglucinolysis either (Supplementary Fig. S13 c and d). These results indicate that EC-P1 and CA-P1 are not dimeric PAs. Third, we performed butanol-HCl boiling experiments. The results showed that the butanol-HCl boiling of PB2 produced red cyanidin pigment (Fig. 4i), however, that of EC-P1 could not produce any red pigment (Fig. 4j). This result indicates that the butanol-HCl boiling cannot cleave EC-P1 to form anthocyanidins. Moreover, neither could the butanol-HCl boiling cleave EC-P2, CA-P1, and CA-P2 to produce red cyanidin pigment. In conclusion, EC-P1, EC-P2, CA-P1, and CA-P2 are neither dimeric B-type nor A-type PAs; instead, they are novel dimeric flavonoids.

**Fig. 4.**
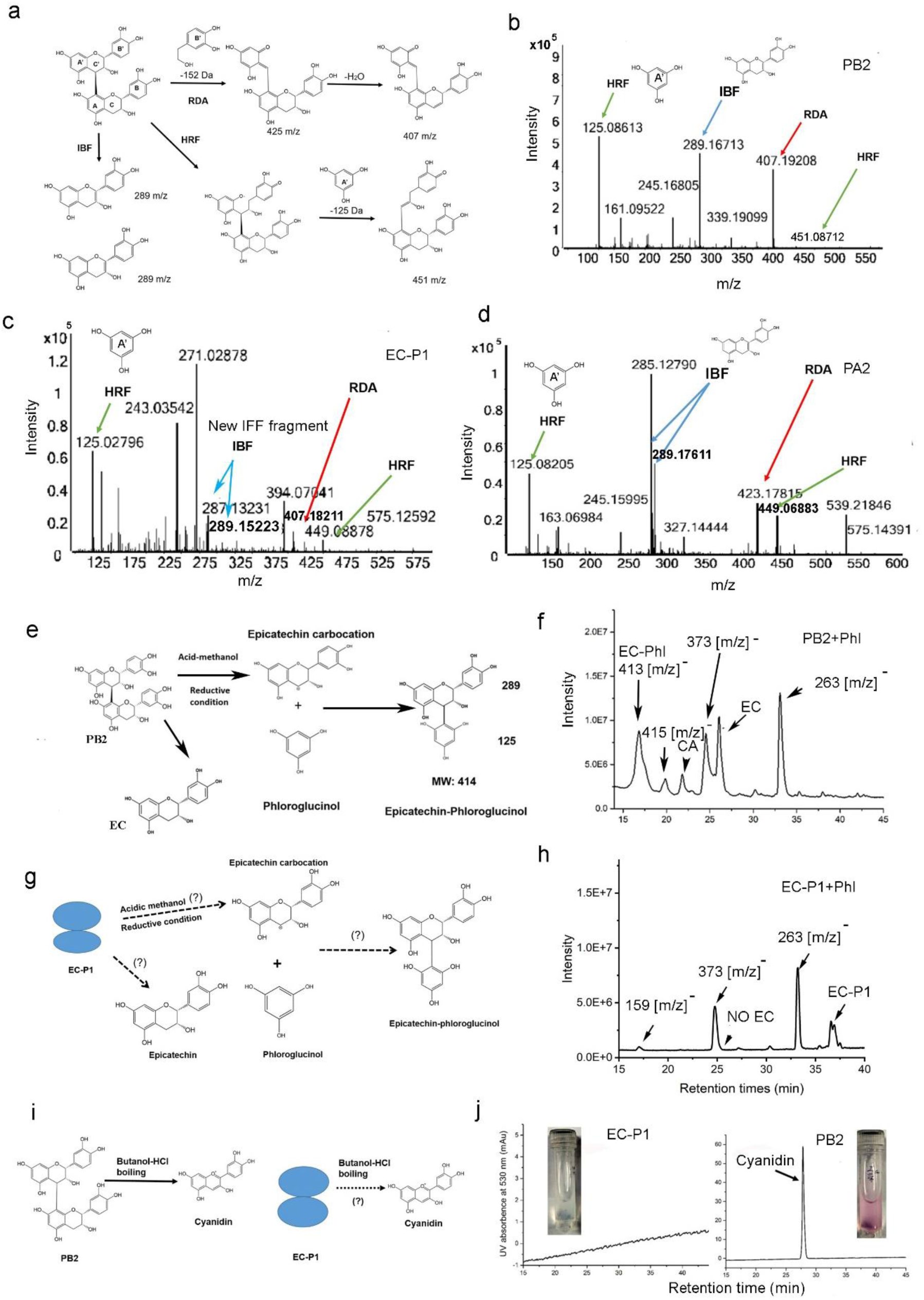
EC-P1 is a novel dimeric flavonoid. MS, phloroglucinol, and butanol-HCl boiling analyses were completed to compare the properties of EC-P1 and PB2. **a,** a scheme shows three fragmentation mechanisms of PB2 by collision induced dissociation (CID), Retro-Diels-Alder (RDA), heterocyclic ring fission (HRF), and interflavan bond fission (IBF). **b-d,** MS/MS profiles show that PB2 (b), EC-P1 (c), and PA2 (d) have the same fragmentation mechanisms, RDA, HRF, and IBF. However, except for the fragment of 125 resulted from HRF, main fragments from IBF and RDA are different among three compounds. These fragment patterns indicate that although the three compounds have the same 125 fragment resulted from A- and A’-ring, EC-P1 is a different compound from PB2 and PA. **e-f,** the phloroglucinolysis (phloroglucinol-HCl based degradation) of PB2 produces epicatechin (EC) and epicatechin-phloroglucinol (EC-Phl). (**e**), a scheme shows the theory that PB2 is degraded to produce epicatechin (EC) and epicatechin-phloroglucinol (EC-Phl). (**f**), the TIC profile detects EC-Phl with 413 [m/z]^-^ and EC derived from PB2. The 263 and 373 [m/z]^-^ values resulted from buffer used in the phloroglucinolysis. **g-h,** the phloroglucinolysis (phloroglucinol-HCl based degradation) demonstrates that EC-P1 is not a dimeric PA compound. (**g**), a scheme presents a hypothesis that the EC-P1 degradation may produce epicatechin (EC) and epicatechin-phloroglucinol (EC-Phl). (**h**) A TIC profile shows that there are no EC-Phl and EC formed from the phloroglucinol-HCl based degradation of EC-P1. These data do not support this hypothesis in (**g**). **i-j,** the butanol-HCl boiling based cleavage shows that EC-P1 is not a PA compound. (**i**), a scheme presents the theory that the butanol-HCl boiling of PB2 produces cyanidin and a hypothesis that the butanol-HCl boiling of EC-P1 may produce anthocyanidins. (**j**), HPLC profiles and coloration of the boiling show that EC-P1 cannot produce anthocyanidins (left) but PB2 can produce cyanidin (right) after the butanol-HCl boiling. These data do not support this hypothesis; instead, support that EC-P1 is not a PA compound.

### Purification of EC-P1 and unambiguous assignment of twenty-two protons by 1H NMR spectrum analysis

We purified EC-P1 via TLC and HPLC preparation isolations (Supplementary Fig. S14a-b) for structural determination by 1H NMR spectrum and other NMR spectrum analysis described below. HPLC-MS analysis showed that the isolated EC-P1 was one single peak (Supplementary Fig. S14c). EC-P1 is a yellowish compound and has maximum visible absorbance at 388 nm (Supplementary Fig. S14c-d). EC-P1 could not be crystalized. We performed 1H NMR spectrum tests to assign protons of EC-P1. Given that EC-P1 shared structural features with PB2, we used this standard as a reference to assign protons and carbons. The 1H NMR test at 240K obtained PPM values for 26 protons of PB2. 16 protons were assigned on A6, A8, B2’, B5’, B6’, C2, C3, C4, E6, F2’, F5’, F6’, G2, G3, G4, and G4 of PB2 and 10 protons were assigned to the 10 hydroxyl groups, A5OH, A7OH, B3OH, B4OH, C3OH, E5OH, E7OH, F3OH, F4OH, and G3OH (Fig. 5a, Fig. S15a, and Table 1). The same 1H NMR test of EC-P1 obtained 22 PPM values for 22 protons (Fig. S15b). The PPM values of 13 protons were similar to those of 13 corresponding protons of PB2 and they were assigned to A6, A8, B2’, B5’, B6’, E6, F2’, F5’, F6’, G2, G3, G4, and G4 of EC-P1 (Fig. 5a, Fig. S15b, and Table 1). The PPM values of 9 protons were similar to those of corresponding protons of 9 PB2 hydroxyl groups and they were assigned to be A5OH, A7OH, B3OH, B4OH, E5OH, E7OH, F3OH, F4OH, and G3OH. Given that the molecular weight EC-P1 is 576 (Fig. S14 e), 2 protons fewer than PB2, it has 24 protons. Based on PB2, two additional protons were not assigned by 1H NMR test. Instead, this 1H NMR spectrum test predicted the two other protons be C3OH and C4H (Supplementary Fig. S15b), which were determined by other NMR spectrum tests described below. This result shows that EC-P1 is structurally different from PB2

**Fig. 5.**
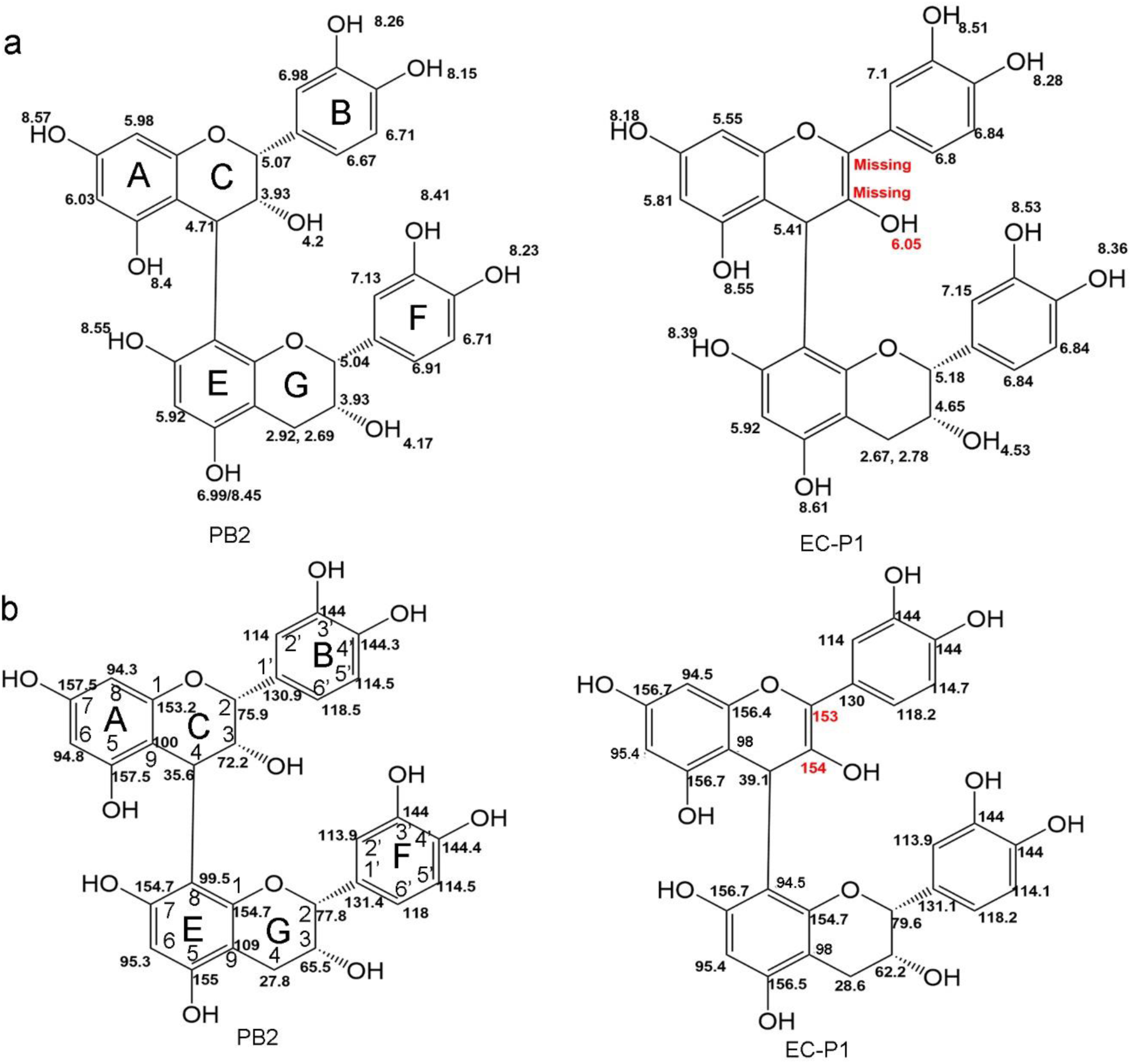
Dimeric EC-P1 consisting of a flaven-flavan structure determined by nuclear magnetic resonance (NMR) analysis. **a,** ^1^H NMR assignments of 15 protons on the six rings of PB2, 13 protons on the six rings of EC-P1, and 10 hydroxyl groups on both PB2 and EC-P1. The black numbers labeled on the two compounds show the similar PPM values of those protons on those rings and hydroxyl groups of PB2 and EC-P1, which demonstrates the same features of these protons and hydroxyl groups of the two compounds. The red numbers labeled on EC-P1 shows that the proton PPM value of the hydroxyl group at C_3_ of EC-P1 is higher than that at C_3_ of PB2. In addition to this hydroxyl group, two protons at C_2_ and C_3_ of EC-P1 are not detected, indicating a double bond formed between the two carbons. **b,** ^13^C NMR assignments of 30 carbons on PB2 and EC-P1. The black numbers labeled on the two compounds show the similar PPM values of these carbons on PB2 and EC-P1, demonstrating the same identity of these carbons. The red numbers labeled on EC-P1 show that the PPM values of C_2_ and C_3_ on EC-P1 are higher than those values of C_2_ and C_3_ on PB2, indicating a double bond formed between the carbons of EC-P1.

**Table 1.** ^13^C and ^1^H NMR assignments of PB2 and EC-P1.

| Position | PB2 |  | EC-P1 |  |
| --- | --- | --- | --- | --- |
|  | <sup>13</sup>C (PPM) | <sup>1</sup>H (PPM) | <sup>13</sup>C (PPM) | <sup>1</sup>H (PPM) |
| A5 | 157.5 | 8.4 | 156.7 | 8.55 |
| A6 | 94.8 | 6.03 | 95.4 | 5.81 |
| A7 | 157.5 | 8.57 | 156.7 | 8.18 |
| A8 | 94.3 | 5.98 | 94.5 | 5.55 |
| C9 | 100 | n/a | 98 | n/a |
| A1 | 153.2 | n/a | 156.4 | n/a |
| B1' | 130.9 | n/a | 130 | n/a |
| B2' | 114 | 6.98 | 114 | 7.1 |
| B3' | 144 | 8.26 | 144 | 8.51 |
| B4' | 144.3 | 8.15 | 144 | 8.28 |
| B5' | 114.5 | 6.71 | 114.7 | 6.84 |
| B6' | 118.8 | 6.67 | 118.2 | 6.8 |
| C2 | 75.9 | 5.07 | 153 | n/a |
| C3 | 72.2 | 3.93, 4.2(C3-OH) | 154 | n/a, 6.05 (C3-OH) |
| C4 | 35.6 | 4.71 | 39.1 | 5.41 |
| E5 | 155 | 8.45 or 6.99 | 156.5 | 8.61 |
| E6 | 95.3 | 5.92 | 95.4 | 5.92 |
| E7 | 154.7 | 8.55 | 156.7 | 8.39 |
| E8 | 99.5 | n/a | 95.4 | n/a |
| G9 | 109 | n/a | 94.5 | n/a |
| E1 | 154.7 | n/a | 154.7 | n/a |
| F1' | 131.4 | n/a | 131.1 | n/a |
| F2' | 113.9 | 6.98 | 113.9 | 7.15 |
| F3' | 144 | 8.26 | 144 | 8.53 |
| F4' | 144.4 | 8.15 | 144 | 8.36 |
| F5' | 114.5 | 6.71 | 114.1 | 6.84 |
| F6' | 118 | 6.67 | 118.2 | 6.84 |
| G2 | 77.8 | 5.04 | 79.6 | 5.18 |
| G3 | 65.5 | 3.93, 4.17(G3-OH) | 62.2 | 4.65, 4.53 (G3-OH) |
| G4 | 27.8 | 2.92, 2.69 | 28.6 | 2.67, 2.78 |

### Unambiguous assignment of the proton of the C3OH group and all 10 hydroxyl groups by seven 1H NMR thermal shifts

We performed 1H NMR spectrum tests at seven different temperatures (240K, 250K, 260K, 270K, 280K, 290K, and 300K) to examine the shifting of protons of hydroxyl groups of EC-P1 and PB2. PB2 was used as the reference. The results showed that the 10 hydroxyl groups of PB2 had ten proton chemical shifts featured by unambiguous different PPM values in seven temperatures (Fig. S16). In comparison with the 1H NMR spectrum of PB2, nine proton chemical shifts of EC-P1 had similar PPM values to those of PB2. Eight of them had 8.18 to 8.61 PPM values, which were similar to eight hydroxyl groups on A, B, E, and F rings of PB2 (Fig. S17). Based on these data, these eight protons were assigned to be A5OH, A7OH, B3OH, B4OH, E5OH, E7OH, F3OH, and F4OH, respectively (Fig. 5a, Fig. S7, and Table 1). The PPM value of the ninth chemical shift was 4.53 (Fig. S17), which was close to that of G3 hydroxyl group of PB2. This proton was assigned to be G3OH. The tenth chemical shift of proton of EC-P1 was corresponding to that of C3OH of PB2. However, the PPM value of the tenth chemical shift was 6.05, which was higher than 4.2, the proton PPM of the C3OH of PB2. Given that double bonds in the rings increase the PPM values of protons of hydroxyl groups (Fig. 5a), the 6.05 PPM is associated with a C=C bond. Based on these, the 6.05 PPM was assigned to C3OH of EC-P1. This result indicates that the increased PPM value of proton chemical shifts at C3OH of EC-P1 results from a double bond formed between C2 and C3 (Fig. 5) or C3 and C4.

### Unambiguous assignment of the protons of C4H, all 24 protons and one C2=C3 double bond by 2D ROESY NMR analyses

We performed 2D ROESY NMR tests to assign the proton of C4H and all 24 protons of EC-P1. PB2 was used as the reference. The 2D ROESY NMR analysis established the cross peaks formed by the 26 protons of PB2 in three pairs of range PPM values. The first pair was 8.0-8.8 (F2 axis from seven –OH groups on A, B, E and F rings) versus 4.6-9.0 (F1 axis, nearby protons) (Supplementary Fig. S18). The second pair was 5.5-7.5 (F2 axis, nine protons of rings on A, B, E and F rings and one proton from E5OH on E ring) versus 3.0-8.8 (F1 axis) (Supplementary Fig. S19). The third pair was 2.5-5.5 (F2 axis, seven protons from C and G rings and two protons of two –OH groups on C and G rings) versus 2.8-8 (F1 axis) (Supplementary Fig. S20). These cross peaks revealed the correlation profiles of 25 protons and their nearby protons on PB2 (Supplementary Fig. S18-S20 and Table S2). It was interesting that the proton of G3OH was identified to be one of nearby protons of F2’H; however, the nearby protons of the G3OH proton could not be identified (Supplementary Fig. S19). The same approach was applied to EC-P1. The resulting 2D ROESY NMR spectrum profiles revealed the 24 protons of EC-P1 and their nearby protons (Supplementary Fig. S21-S24 and Table S2). First, like those 10 nearby protons crossed with the 8 protons of the 8 -OH groups on A, B, E, F, and C rings of PB2, 10 nearby protons of EC-P1 were determined to correlate to the protons of eight -OH groups on A, B, E, and F rings to form 10 crossed peaks (Supplementary Fig. S21, F2 axis; Table S2). Of these ten, one had a PPM value 5.41 on C ring, three had PPM values from 5.5 to 5.92 on A and E rings, and six had PPM values from 6.8 to 7.15 on B and F rings (Supplementary Fig. S21, F1 axis). These PPM values show that these 10 nearby protons are corresponding to those protons of C4H, A6H, A8H, E6H, B2H, B5H, B6H, F2’H, F5’H, and F6’H of PB2. Based on these features, these ten nearby protons were assigned to be the protons of C4H, A6H, A8H, E6H, B2H, B5H, B6H, F2’H, F5’H, and F6’H of EC-P1 (Supplementary Table S2). The 24^th^ proton was assigned to be C4H as predicted by 1H NMR test (Supplementary Fig. 15b). Accordingly, all 24 protons were assigned to EC-P1. Second, six nearby protons with PPM values from 4.0-8.8 were identified to cross with the protons of F2’H, B2’H, B6’H, F5’H, F6’H, or B5’H to form 10 crossed peaks (Supplementary Fig. S22). These six nearby protons of EC-P1 were similar to those nearby protons of PB2, which crossed with the protons of F2’H, B2’H, B6’H, F5’H, F6’H, or B5’H (Supplementary Fig. S19). Based on these data, the six nearby protons of EC-P1 were assigned to the protons of G2H, F3’OH, B3’OH, G3H, F4’OH, and B4’OH (Supplementary Fig. S22 and Table S2). Third, seven nearby protons within PPM values from 6.0-8.8 (F1 axis) were identified to cross with the protons of E6H, A6H, A8H, C4H, or G2H to form nine crossed peaks (Supplementary Fig. S23, F2 axis). These seven nearby protons were similar to those protons of PB2, which crossed with E6H, A6H, A8H, C4H, or G2H (Supplementary Fig. S19 and S20). Based on these data, these seven nearby protons were assigned to be the protons of A7OH, E7OH, E5OH, A5OH, F4’OH, F5’H, and F2’H of EC-P1 (Supplementary Fig. S23 and Table S2). The last, four nearby protons within PPM values from 2.0-6.0 (F1 axis) were identified to cross with the protons of G3H, G4aH, or G4bH to form nine crossed peaks (Supplementary Fig. S24, F2 axis). These four nearby protons were similar to those protons of PB2, which crossed with G3H, G4aH, or G4bH (Supplementary Fig. S20). Based on these data, these four nearby protons were assigned to be the protons of G2H, G4bH, G4aH, or G3H (Supplementary Fig. S24 and Table S2). Based on the cross proton profiles of PB2, the 2D ROSEY NMR analysis allowed assigning the nearby protons of 24 protons of EC-P1. In addition, like G3OH of PB2 (Supplementary Fig. S20), no nearby protons were identified to cross with the proton of G3OH of EC-P1 (Supplementary Fig. S24). Although these data showed these similarities of proton chemical shifts between EC-P1 and PB2 (Table S2), differences between the two compounds were observed. Unlike the cross peak profiles in PB2, no nearby protons crossed with the protons of C2H, C3OH, C3H, and B6’H of EC-P1 were identified (Supplementary Fig. S21-S24 and Table S2). Furthermore, the proton chemical shifts correlating to C2 and C3 positions of EC-P1 were not observed by 2D ROSEY NMR analysis either. This result indicated that two protons were missing at C2 and C3 of EC-P1, which resulted from a double bond formed between the two carbons (Fig. 5) supported by the assignment of C3OH described above. Taken together, the 2D ROESY NMR test results show that EC-P1 has 10 protons from its hydroxyl groups and 14 protons from carbons of its six rings. Of the 24, the PPM values of 23 protons were similar to those of corresponding protons of PB2. The PPM values of C3OH between EC-P1 and PB2 are different (Fig. 5 and Table S2). Compared to PB2, the C2 and C3 positions of EC-P1 lack two protons, indicating a double bond that is formed between the two carbons. Accordingly, this new double bond leads to the result that the PPM value of the C3OH proton of EC-P1 is higher than that of the C3OH proton of PB2 (Fig. 5a). In conclusion, these data demonstrate that EC-P1 and PB2 have the same A, B, E, F, and G ring structures but have a different C-ring structure featured by a double bond between C2 and C3 (Fig. 5).

### Assignment of thirty carbons of EC-P1 by 2D HMBC NMR analysis and nomenclature of novel flaven-flavan structure-papanridins

We performed 2D HMBC NMR tests to correlate chemical shifts among protons and carbons of EC-P1 and assign all carbons. PB2 was used as the reference for this assignment. The 2D HMBC NMR tests created three pairs of range PPM values between 26 protons and 30 carbons of PB2 (Fig. 5a), 1) 8.0-8.8 (the protons of seven -OH groups, F2 axis) versus 90-160 (carbons, F1 axis) (Supplementary Fig. S25), 2) 5.8-7.5 (nine protons of A, B, E, and F rings and the proton of E5OH) versus 70-160 (carbons) (Supplementary Fig. S26), and 3) 2.8-5.4 (seven protons of C and G rings and the two protons of C3OH and G3OH) versus 20-160 (carbons) (Supplementary Fig. S27). Based on these data, the correlations of 30 carbons and 26 protons of PB2 were established (Fig. 5b and Table S3). The same method was applied to EC-P1. The 2D HMBC NMR tests also created three pairs of range PPM values between 24 protons and 30 carbons of EC-P1 (Fig. 5a), 1) 8.0-8.8 (the protons of eight –OH groups, F2 axis) versus 90-160 (carbons, F1 axis) (Supplementary Fig. S28), 2) 5.2-7.2 (nine protons on A, B, E and F rings, two protons on C and G rings, and the proton of C3OH) versus 40-160 (carbons) (Supplementary Fig. S29), and 3) 2.5-5 (three protons from the G ring and the proton of G3OH) versus 50-150 (carbons) (Supplementary Fig. S30). First, based on these paired cross peaks of chemical shifts, 15 carbons on A, B, E, and F rings of EC-P1 were identified to correlate with the protons of E5OH, A5OH, F3’OH, B3’OH, E7OH, F4’OH, B4’OH, and A7OH. The PPM values of these 15 carbons were similar to those of 15 carbons of A, B, E and F rings of PB2 (Fig. 5b). Based on these data, these 15 carbons were assigned to the A, B, E, and F rings of EC-P1. Four were A5, A6, A7, and A8 of the A ring; four were B2’, B3’, B4’, and B5’ of the B ring; three were E6, E7, and E8 of the E ring; and four were F2’, F3’, F4’ and F5’ of the F ring (Fig. 5 b and Table S3). Second, 22 carbons on the six rings of EC-P1 were identified to correlate with the protons of F2’H, B2’H, F6’H, F5’H, B’5H, B6’H, E6H, A6H, A8H, C4H, and G2H (Supplementary Fig. S29). The PPM values of the 22 carbons were similar to those of the corresponding 22 carbons on the six rings of PB2 (Fig. 5). Based on these data, four were assigned to A5, A6, A7, and A8 of the A ring. Four were assigned to B1’, B3’, B4’, and B6’ of the B ring. Four were assigned to C2, C3, C4, and C9 of the C ring. Two were assigned to E5 and E8 of the E ring. Six were assigned to F1’, F2’, F3’, F4’, F5’, and F6’ of the F ring. Two were assigned to G2 and G9 of the G ring (Table S3). Third, two carbons on the G ring of EC-P1 were identified to correlate with the protons of G3H and G4aH (Supplementary Fig. S30). The PPM values of the carbon chemical shifts of the two carbons were similar to those of G2 and G9 of PB2 (Fig. 5b). Based on these, these two carbons were assigned to G2 and G9 of EC-P1 (Fig. 5b). Although these data showed the similarities of PPM values for these assigned carbons of PB2 and EC-P1, differences were observed in the two compounds. Unlike PB2 that the proton of C4H correlates with the carbon of C2H with a 75.9 PPM and the carbon of C3H with a 72.2 PPM (Supplementary Fig. S27), the proton of C4H of EC-P1 correlates with carbons of C2H and C3H, which have 153 and 154 PPM, respectively (Fig. 5b and Supplementary Fig. S29). These features distinguish EC-P1 from PB2. The PPM values of those 24 carbons associated with a C=C bond on A, B, E, and F of PB2 and EC-P1 are from 94 to 157. These data indicate that 153 and 154 PPM values of C2 and C3 of EC-P1 result from the formation of a double bond between them (Fig. 5). Therefore, C2 and C3 of the C-ring of EC-P1 form a double bond. Taken together, the 2D HMBC NMR analysis allows the assignment of the correlation of 30 carbons and 24 protons (Fig. 5).

Based on all NMR data, EC-P1 is composed of 30 carbons, 14 protons, and 10 hydroxyl groups. Not only does it differ from dimeric B-type procyanidins, but also it distinguishes from dehydrodicatechin A and B, which only have 7 hydroxyl groups [21, 23]. These assignments showed that EC-P1 was a novel dimeric compound consisting of epicatechin (starter unit or the bottom) and flav-2-en-3-ol (the upper unit) linked by an interflaven-flavan bond between C4 of the upper unit and C8 of the bottom unit (Fig. 5). Based on the structural features of the building units, EC-P1 is a dimeric flaven-4→8-flavan (FF) flavonoid. Given that this compound is formed from the catalysis of FP from PAP1 cells and biosynthesized in <u>PAP</u>1-<u>ANR</u> plants and share certain features of PAs, we named this type of novel compounds as “papanridin”. Subsequently, we named EC-P1 and EC-P2 from epicatechin as papanridin ECII-A and -B, and CA-P1 and CA-P2 as papanridin CAII-A and -B, in which “II” means dimers.

### Evidence of C4→C8 linkage based polymerization of EC-P1 and formation of higher oligomeric papanridins

We hypothesized that FP polymerized papanridin via carbocation formed in the reactions and the formation of C4→C8 linkage. FP catalyzed epicatechin to flav-2-en-3-ol carbocation, and then epicatechin nucleophilically attacked the carbocation to form papanridin ECII A and B (Fig. 6a). We also hypothesized that the flav-2-en-3-ol carbocation could be nucleophilically trapped by phloroglucinol to form flav-2-en-3-ol-phloroglucinol (MW, 412 Dalton) (Fig. 6a). To test these hypotheses, phloroglucinol was added to the incubation consisting of pH 6.0 buffer, EC, FP, and H_2_O_2_. LC/MS analysis showed that an apparent peak with a 411 [m/z]^-^ overlaid with EC-P1 formed from the FP reaction, but it was not observed from two controls (Fig. 6b). Further EIC analysis showed this 411 [m/z]^-^ compound from the FP and phloroglucinol incubations but not from controls (Fig. 6 c). To characterize this 411 [m/z]^-^ compound, MS/MS analysis revealed two characteristic fragments, 125 from phloroglucinol and 285 likely from flav-2-en-3-ol (Fig. 6d). This feature was substantiated by MS/MS fragmentation of epicatechin-phloroglucinol, a positive control, which produced two main fragments, 125 from phloroglucinol and 287 from epicatechin (Fig. 6 d). To further demonstrate the presence of flav-2-en-3-ol carbocation, we used quercetin and taxifolin as negative controls, because these two molecules with a carbonyl group at C4 cannot form carbocation at C4. We incubated FP, quercetin or taxifolin, H_2_O_2_, and phloroglucinol. LC/MS analysis could not detect the formation of quercetin-phloroglucinol and taxifolin-phloroglucinol from the reactions (Supplementary Fig. S31). These data supported the presence of flav-2-en-3-ol carbocation formed in the enzymatic reaction of FP and EC. Moreover, we hypothesized that trimers with a MW 698 could be formed from the incubations of phloroglucinol, FP, and EC. EIC analysis revealed five 697 [m/z]^-^ peaks (Supplementary Fig. S32a). MS/MS analysis showed that the CID of 697 generated 571, 409, and 287 fragments, which corresponded to flav-2-en-3-ol→flav-2-en-3-ol carbocation molecules (571=572 - 1), flav-2-en-3-ol→phloroglucinol (409= 411-2), and flav-2-en-3-ol, respectively (Supplementary Fig. S32b). Therefore, these compounds are trimeric flav-2-en-3-ol C4→C8 flav-2-en-3-ol C4→phloroglucinol (Supplementary Fig. S32c). Based on these data, LC/MS was performed to examine whether higher papanridin oligomers were formed in the enzymatic reactions that consisted of FP, H_2_O_2_, and EC with (Fig. 6c) or without phloroglucinol (Supplementary Figs. S8b and S14a). The resulting data showed that papanridin ECII-A and -B were produced in the two types of FP reactions (Fig. 6c). Furthermore, EIC analysis showed that two trimers, two tetramers, and one pentamer were produced and their [m/z]^-^ values were 861, 1147, and 1433; however, these compounds were not observed from all BSA control incubations (Supplementary Fig. S33a). MS/MS analysis further showed that these new compounds were trimeric flav-2-en-3-ol C4→C8 flav-2-en-3-ol C4→C8 epicatechin (MW, 862 Dalton), tetrameric flav-2-en-3-ol C4→C8 flav-2-en-3-ol C4→C8 flav-2-en-3-ol C4→C8 epicatechin (MW, 1148 Dalton), and pentameric flav-2-en-3-ol C4→C8 flav-2-en-3-ol C4→C8 flav-2-en-3-ol C4→C8 flav-2-en-3-ol C4→C8 epicatechin (MW, 1434 Dalton) (Supplementary Fig. S33a-b).

**Fig. 6.**
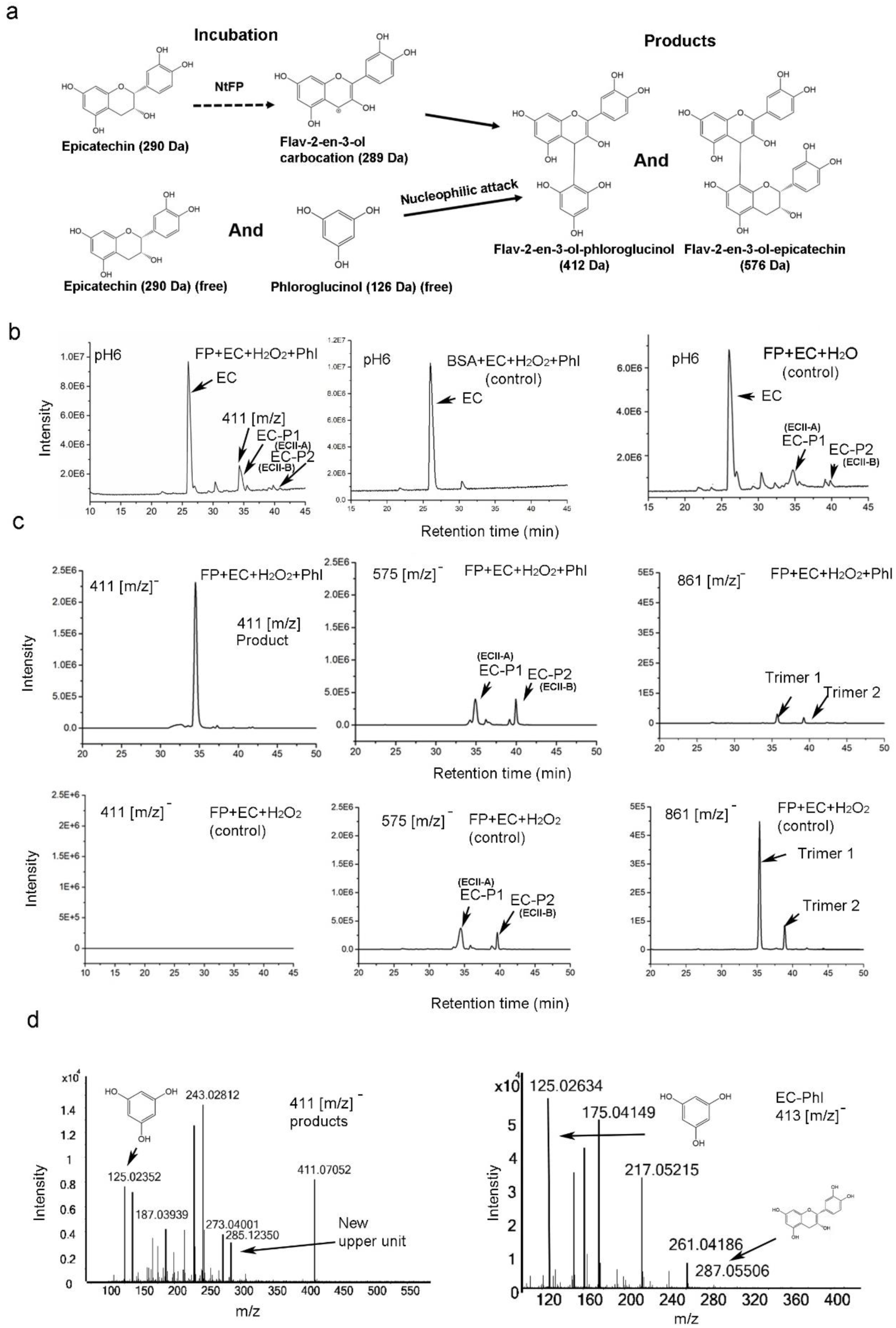
Evidence for flav-2-en-3-ol carbocation produced in enzymatic reactions and in papanridins. Enzymatic reactions consisted of flavanol polymerase (FP) or BSA (control), epicatechin (EC), hydrogen peroxide, and phloroglucinol added or not added in pH6 buffer. **a,** a scheme presents a hypothesis that flav-2-en-3-ol carbocation is produced in the enzymatic reactions; free nucleophilic epicatechin and phloroglucinol in the reactions can competitively attack the carbocation at C_4_ to form dimeric or oligomeric compounds. **b,** TIC profiles showed that the incubation consisting of FP, epicatechin (EC), hydrogen peroxide, and phloroglucinol produced EC-P1 (ECII-A), EC-P2 (ECII-2), and a new compound with a 411 [m/z]^-^ (left) overlaid with EC-P1, while the control incubation consisting of BSA, EC, hydrogen peroxide, and phloroglucinol did not produce these compounds (middle). Meanwhile, the control incubation consisting of FP, EC, and hydrogen peroxide produced EC-P1 and EC-P2 but did not produce the new compound with a 411 [m/z]^-^ (right). **c,** The upper EIC profiles demonstrated that the incubation consisting of FP, epicatechin (EC), hydrogen peroxide, and phloroglucinol produced the new 411[m/z]^-^ compound (left), EC-P1 (ECII-A) and EC-P2 (ECII-B) with a 575 [m/z]^-^ (middle), and two trimers with an 861 [m/z]^-^ value (right). The bottom EIC profiles demonstrated that the incubation consisting of FP, EC, and hydrogen peroxide did not produce the new 411[m/z]^-^ compound (left), but produced EC-P1 (ECII-A) and EC-P2 (ECII-B) (middle) and two trimers (right). **d,** the new 411 [m/z]^-^ compound is not epicatechin-phloroglucinol. MS/MS profiles showed the different main fragments between the 411 [m/z]^-^ compound and epicatechin-phloroglucinol. The 411 [m/z]^-^ compound has a main fragment of 285, while epicatechin-phloroglucinol has a main fragment of 287. Both have the 125.02 fragment, which corresponds to phloroglucinol. Therefore, 411 was fragmented to 125 and 285 and epicatechin-phloroglucinol (413) was fragmented to 125 and 287, indicating the presence of a new flava-2-en-3-ol carbocation rather than epicatechin carbocation formed in the enzymatic reactions.

### Decrease of dimeric papanridins in fp mutant of Arabidopsis thaliana

To genetically support the biosynthesis of papanridins in plants, we blasted *FP* homologs in the Arabidopsis genome and identified one homolog, namely *AtFP* here. We further obtained one *atfp* mutant. LC-MS/MS based profiling detected papanridin ECII-A and -B in the seed extracts of wild-type *A. thaliana* but not in those of *atfp* mutants (Fig. 7a and Supplementary Fig. S34). In addition, LC-MS/MS analysis showed that the level of EC was higher in the mutant than in wild type ones (Supplementary Fig. S35). These results indicate the biosynthesis of papanridins from EC in *A. thaliana*.

**Fig. 7.**
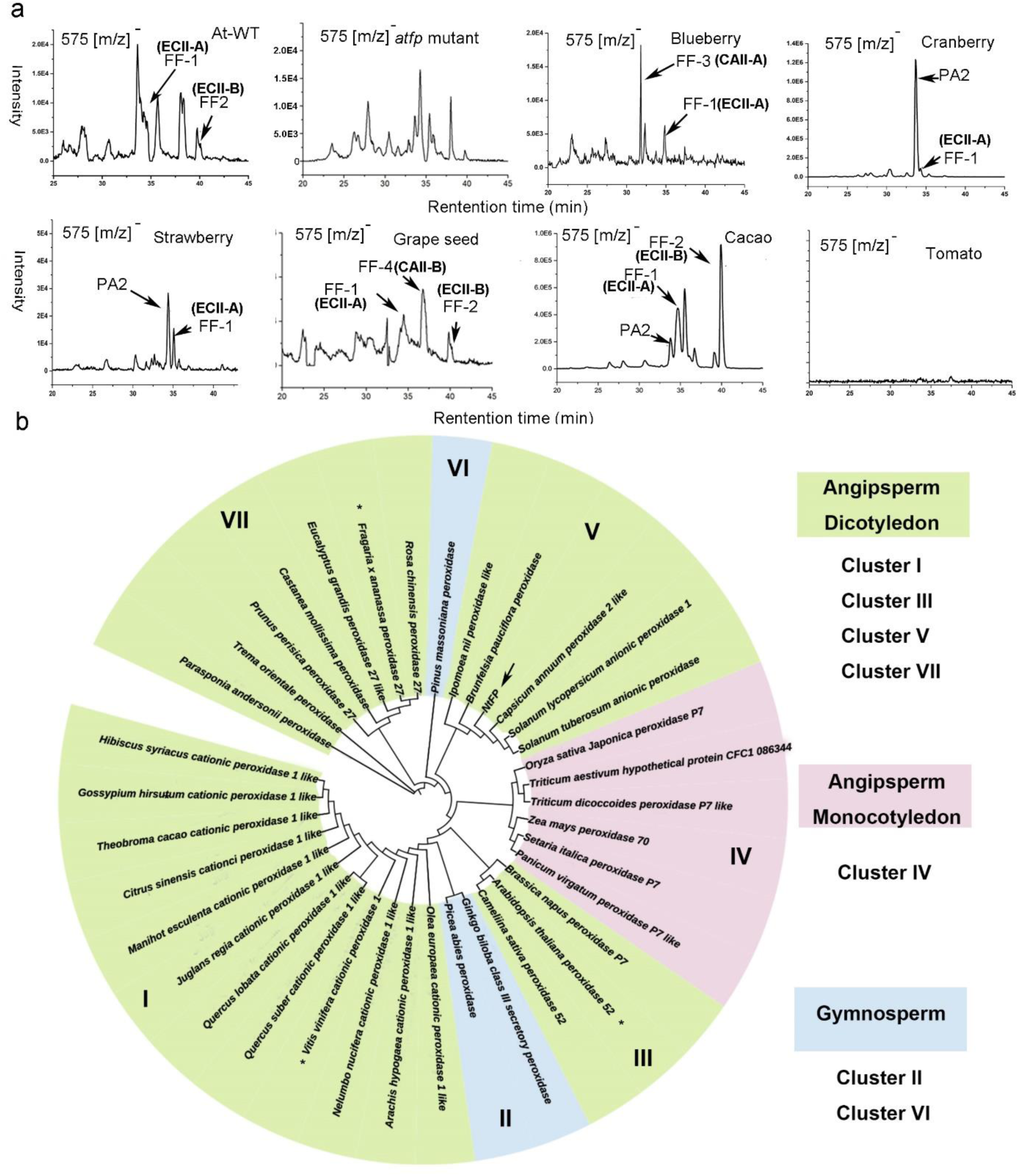
Prevalence of dimeric papanridin (flaven-flavan, FF) compounds in flavan-3-ol- and PA-rich tissues of plants. Mature seeds of *Arabidopsis thaliana* and the mutant *atfp*, berries of blueberry, cranberry, strawberry, and tomato, seeds of grape and cacao were analyzed with LC-MS/MS. **a,** EIC profiling detected papanridin ECII-A (FF1) and ECII-B (FF2) in seeds of Arabidopsis, ECII-A (FF1) and CAII-A (FF3) in blue berry, ECII-A (FF1) in cranberry and strawberry, ECII-A (FF1), ECII-B (FF2), and CAII-B (FF4) in grape seeds, ECII-A and -B in cacao seeds. However, these dimmers could not be detected in seeds of *atfp* mutant and berry of tomato. PA2 was detected in cranberry, strawberry, and cacao seeds. **b,** an unrooted phylogenetic tree was developed with amino acid sequences of FP (NtFP) and 36 homologs across gymnosperm and angiosperm. The numbers on branches are approximate likelihood ratio test values that are used to provide statistical support for the branches of the tree.

### Papanridins in flavan-3-ol-rich plants and phylogenetic analysis

LC-MS/MS profiling was performed to analyze dimeric or oligomeric papanridins (flaven-flavan, FF) in flavan-3-ol and PA-rich plants to understand whether these new flavonoids are prevalent in plants. Plant tissues analyzed included berries of blueberry, cranberry, and strawberry, muscadine grape seeds, and cacao seeds. In addition, tomato berry lacking flavan-3-ols was used as a negative control. The resulting data showed that blueberry produced papanridin ECII-A and CAII-A, cranberry and strawberry had papanridin ECII-A, muscadine seeds produced papanridin ECII-A and -B, and CAII-D, and cacao seeds produced papanridin ECII-A and -B. However, tomato did not produce these compounds (Fig. 7a). Meanwhile, A-type procyanidin A2 was identified in cranberry, strawberry, and cacao seeds (Fig. 7a). EC was produced in blueberry, cranberry, strawberry, muscadine seeds, and cacao seeds. CA was produced in blueberry, cranberry, and muscadine seeds. Nevertheless, neither of them was produced in tomato berry (Supplementary Fig. S35). These data indicate the biosynthesis of papanridins in these flavan-3-ol and PA-rich plants.

We further blasted FP homologs in plant genomes curated in the GenBank at NCBI and obtained 37 homologs across gymnosperms and angiosperms. Included were homologs from *A. thaliana*, cacao, bunch grape (*V. vinifera*), and strawberry, which were shown to produce dimeric papanridins. Due to the unavailability of genomic sequences for blueberry and cranberry, homologs were not obtained for these two plants. The 37 sequences were used to establish an unrooted phylogenetic tree (Fig. 7b), which was featured by seven clusters, I-VII. It was interesting that three gymnosperms were grouped into two clusters. Six monocotyledonous plants were grouped into one cluster. Dicotyledonous plants were grouped into four clusters. Cacao and grape FPs were grouped to cluster I. Arabidopsis, tobacco, and strawberry FPs were grouped into cluster III, V, and VII. Although other homologs remain for studies, this phylogenetic analysis indicates that FP is prevalent in the plant kingdom.

## Discussion

Our findings add a group of novel oligomeric flavonoids prevalent in the plant kingdom, namely papanridins herein. Plant flavonoids are the largest group of plant polyphenols [19, 48–50]. By 2000, more than 5000 flavonoids were reported from the plant kingdom [11, 13, 14, 51]. In the past nearly two decades, as more phytochemical isolations were investigated, approximately 8000 new flavonoids were reported from plants [19], which were classified into chalcone, flavanones, flavones, aurones, dihydroflavonols, flavonols, flavan-3,4-diols, anthocyanins, flavanols, proanthocyanidins, isoflavonoids, bioflavonoids, or neoflavonoids [3, 19, 52–55]. PAs are the only oligomeric or polymeric flavonoids that are synthesized from flavanols via an interflavan bond between starter and extension flavan-3-ol units [20]. Herein, our findings via engineered red cells show a group of novel oligomeric or polymeric flavonoids featured by a skeleton flaven-flavan (FF). We termed them papanridins. Our experimental data characterize that papanridins are derived from flavanols but have different structural, chemical, and physical features from PAs. In addition, when we observed the yellowish color of EC-P1 (ECII-A), we proposed that it might be dehydrodicatechin featured by 7-OH groups and C-O interflavan linkage resulted from the catalysis of PPO [21–23]. However, NMR data showed EC-P1was featured by 10-hydroxyl groups (Fig. S17). Moreover, higher oligomers formed from EC-P1. These data demonstrate that papanridins are different from dehyrodicatechins. In summary, papanridins have the following chemical and physic features. 1) Flavanols are initial substrates of papanridins. 2) Papanridins were polymerized from flavanols as the starter unit and flav-2-en-3-ols as extension unit (Figs. 5 and 8). 3) The bottom and the first upper unit are linked by an interflaven-flavan bond formed at C_8_ and C_4_. 4) The extension units are linked by interflaven bounds. 5) Dimeric papanridin ECII-A and -B, papanridin CAII-A and -B, trimers, and tetramers are yellowish and have an absorption maximum at 388 nm (Supplementary Fig. S14a and d). 6) Neither can phloroglucinolysis occur in papanridins in the acidic methanol condition, nor can the butanol-HCl boiling cleave them to produce anthocyanidins (Fig. 4 g-j). 7) Papanridins are weakly soluble but unstable in water and can be oxidized to quinones. 8) Papanridins are prevalent in flavan-3-ol rich plants (Fig. 7). What functions papanridins have in the plants is unclear. Based on metabolic profiling of *FP* transgenic PAP1-BAN plants and *atfp* mutant plants, one hypothesis is that papanridin is likely associated with the biosynthesis of PAs (Fig. 8a). As *FP* was overexpressed in PAP1-BAN plants, the contents of PB2 and total PAs were significantly increased (Supplementary Fig. S12), while the content of EC was significantly decreased (Fig. 3h and j). Since this is the first report, it can be anticipated that a great number of studies are necessary to understand the functions of papanridin in plants.

**Fig. 8.**
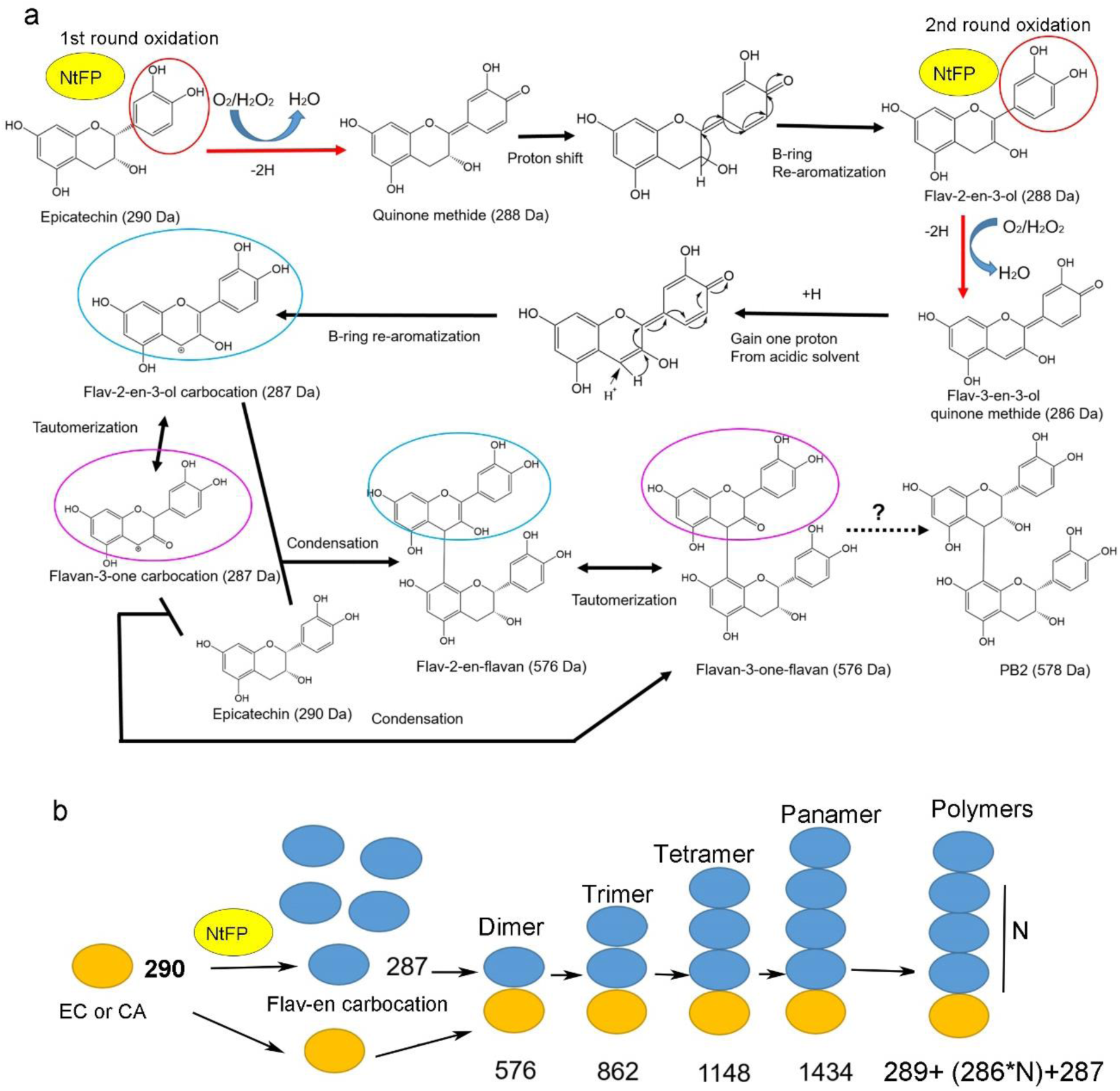
Hypothetic mechanisms proposed for the formation of dimeric and oligomeric papanridins from flavanols via flav-2-en-3-ol carbocation polymerized by FP. **a,** a pathway is proposed to start with the FP-catalyzed oxidation of epicatechin to form flav-2-en-3-ol carbocation and then dimeric papanridins. First, FP oxidizes epicatechin to quinone methide, which undergoes a spontaneous proton shift and B-ring re-aromatization to form flav-2-en-3-ol. Second, FP oxidizes flav-2-en-3-ol to its quinone methide, which undergoes addition of one H^+^ from a weak acidic condition and then B-ring re-aromatization to form flav-2-en-ol carbocation. Then, available epicatechin molecules attack the C_4_ of the carbocation to form dimeric flav-2-en-3-ol→epicatechin, which further may tautomerize to flavan-3-one→epicatechin and flav-3-en-3-ol→epicatechin. Although what functions dimeric papanridins have is unknown, it can be speculated that they might be reduced to produce PAs. **b,** a carton is created to show the polymerization mechanism by which in the oxygen presence, plant cells use FP to biosynthesize, dimers, trimers, and higher degree oligomers. Once one dimeric papanridin (flaven→flavan) forms, it can nucleophilically attack flav-2-en-3-ol carbocation to form a trimer. Further sequential addition of flav-2-en-3-ol carbocation produces higher degree oligomers.

Our findings demonstrate that FP catalyzes the polymerization of papanridin via flavan-3-ols and flav-2-en-3-ol carbocations. Flavanols are the substrates of FP and the start unit of papanridins, while flav-2-en-3-ols are the extension units. Our trapping experiments with nucleophilic phloroglucinol provided evidence for the presence of flav-2-en-3-ol carbocation in the catalytic reactions of FP and EC (Fig. 6). Although the mechanism by which this carbocation is formed from EC in the reactions remains for investigations, we hypothesize two steps that FP oxidizes EC to form flav-2-en-3-ol carbocation (Fig. 8a). The first oxidation step is that in the presence of oxygen, FP oxidizes EC to quinone methide. The proton at C_3_ shifts to the B-ring via C_2_ leading to the B-ring re-aromatization that forms flav-2-en-3-ol. The second oxidation step is that FP oxidizes flav-2-en-3-ol to form flav-3-en-3-ol quinone methide, to which one proton from the acidic condition was added. This proton addition leads to the second time of B-ring re-aromatization to form flav-2-en-3-ol carbocation with a positive charge at C_4_. EC then nucleophilically attacks the flav-2-en-3-ol carbocation leading to the condensation via the formation of an interflaven-flavan bond. Accordingly, a new flav-2-en-3-ol→EC with a MW 576 Dalton is formed. Given that two dimeric compounds are obtained with the same MW 576 Dalton, we further hypothesize that like B-type PAs, there are two types of stereo configurations, flav-2-en-3-ol-(4α→8)-EC and flav-2-en-3-ol-(4β→8)-EC. To provide evidence that flav-2-ev-3-ol was produced in the incubations, we used EIC to analyze metabolites from the FP catalysis of EC and CA. The resulting data showed that four from EC and three from CA had a 287 [m/z]^-^, while these compounds were not detected from negative controls (Supplementary Fig. S36a). Given that only EC, FP, and H_2_O_2_ were used in the reactions, these compounds were formed from EC via the catalysis of FP. Besides, given that we used negative mode for ionization, the 287 [m/z]^-^ value resulted from non-charged compounds with a MW 288 Dalton, which is the MW of flav-2-en-3-ol. Why were more than one 287 [m/z]^-^ peaks detected from the catalysis? Based on our previous report that flavan-3-one and flav-3-en-3-ol are two tautomers from the spontaneous tautomerization of flav-2-en-3-ol in the formation of EC [30, 35], we hypothesize that additional peaks result from the tautomerization. A comparison among the 287 [m/z]^-^ peaks (Supplementary Fig. S36a), one peak (P2) was obtained from both EC and CA. This result showed that the same compound was produced from EC and CA, indicating that the configuration of P2 is not affected by these two isomers. Thus, P2 does not have a stereo configuration between C_2_ and C_3_; instead, it has a double bond C_2_ and C_3_. In another words, P2 is flav-2-en-3-ol (Supplementary Fig. S36c). In addition, we propose that three other peaks from EC are 2R-flav-3-en-3-ol, 2S-flav-3-en-3-ol, and 2R-flavan-3-one and two other peaks from CA are 2S-flavan-3-one and 2S-flav-3-en-3-ol (Supplementary Fig. S36c). We further propose that flav-2-en-3-ol can spontaneously tautomerize to flavan-3-one and flav-3-en-3-ol. The past studies have reported that these three tautomers occur in plants. Our and other recent reports demonstrated that flav-2-en-3-ol was involved in the formation of PAs [30, 33, 36]. A flav-2-en-3-O-glycoside was reported to associate with the formation of cyanins in black soybean [56, 57]. Flav-3-en-3-ol glycosides were also isolated from a medicinal plant [58]. Based on our metabolic profiling of dimeric papanridins in Arabidopsis seeds and five flavanol-rich plants and FP homolog analysis (Fig. 7), we propose a model to interpret the polymerization mechanism of papanridins (Fig. 8b). In this model, additional flav-2-en-3-ol carbocations are added to the upper unit of dimeric papanridins to form oligomers and then polymers. The evidence was trimers, tetramers, and pentamers that were observed in the enzymatic reactions of FP (Supplementary Fig. S33a).

## Materials and methods

### Plant materials and chemical reagents

Wild type tobacco (*Nicotiana tabacum* var. Xanthi) [40], red/purple PAP1 transgenic 6R and wild-type P3 tobacco cell lines [40], lightly red PAP1-ANR (PAP1-BAN) tobacco [44], *Arabidopsis thaliana*, blue berries, cranberry, muscadine grape berries, raspberry, cacao seeds, and tomato fruit were used in this study. Seeds and fruits were purchased from grocery stores. Red/purple PAP1 transgenic 6R and wild type P3 cell lines were established from red PAP1 tobacco (*N. tabacum* var. Xanthi-PAP1) and wild-type tobacco plants, respectively [40]. Red 6R cells constitutively overexpress Arabidopsis *PAP1* gene and produces high production of anthocyanins, while wild type P3 cells do not. PAP1-ANR (BAN) tobacco is a homozygous progeny of *PAP1* and *ANR* transgenic plant, which produces anthocyanins, flavanols, and proanthocyanidins in leaves and flowers [44]. Plants were grown in the greenhouse supplemented with natural light.

Authentic (+)-catechin, (±)-catechin, (-)-epicatechin, (-)-epigallocatechin, procyanidin A2, procyanidin B2, phloroglucinol, and catechol were purchased from Sigma (St. Luis, USA). These compounds were dissolved in methanol to make 10 mg/ml stocking solutions, which were stored in a -20°C freezer. Organic solvents used are specified in each different experiment.

### Callus and cell suspension culture

The medium, subculture of 6R and P3 cells, and tissue culture conditions were as reported previously [40]. Since 2008, these two cell lines have been subcultured for studying anthocyanin biosynthesis and plant flavonoids. Briefly, the medium was composed of MS medium (pH 5.7) containing 0.25 mg l^-1^ 2,4-dichlorophenoxyacetic acid, 0.10 mg L^-1^ kinetin, 30 g l^-1^ sucrose and 0.8% (w/v) phytoagar. Calli (Supplementary Fig. S1) were aseptically subcultured with a 15-day interval in a growth chamber facilitated with 22 °C and a 16 h/8 h light/dark photoperiod. Cell suspension cultures were established from 6R and P3 cells. The liquid medium used was prepared from the subculture medium by the removal of agar. A 50 ml aliquot was contained in a 250 ml Erlenmeyer flask (Supplementary Fig. S1). Each flask was inoculated with 5 g calli and then placed on a rotary shaker with a speed of 100 rpm in a growth chamber with the same growth condition as described above. After 15 days, suspension cells were filtered with autoclaved 10 and 50 µm metal sieves. Cells harvested between the two types of sieves were subcultured in 50 ml liquid medium. After 3 times of subculture, appropriate cell suspension cultures were established. Suspension cultures were developed for 6R and P3 cells as comparison. To obtain sufficient amount of cells, 10 g (fresh weight, FW) suspension cells after filtering were cultured in 1-liter liquid medium contained in a 6-liter E-flask and cultured for 15 days (Supplementary Fig. S1b). Cells were filtered (Supplementary Fig. S1 d-e) and collected to liquid nitrogen, and then stored in an 80°C freezer until the use for the following experiments. The remaining liquid medium was transferred to 500 ml tubes for 10 min of centrifugation at 8,000 ×g and 4°C. The supernatant (Supplementary Fig. S1g) was concentrated and then stored in 4°C for the following experiments.

### Protein Extraction

Frozen 6R and P3 cells were ground into fine powder in liquid nitrogen. The protein extraction buffer prepared was composed of 0.1 M Tris-Cl (pH 7.0) and 10 mM EDTA. Thirty grams of powder were suspended in 150 mL of the protein extraction buffer contained in a 500 ml tube and vortexed for one min. The resulting mixture was sonicated for 30 min and then shaken on a rotary shaker at 100 rpm for 20 min at 4°C. The suspension mixture was centrifuged for 20 min at 10,000 rpm under 4°C. The supernatant was transferred to a clean beaker and then filtered through a Whatman filter paper to obtain clear protein extracts (Supplementary Fig. S1f). In addition, calli were also extracted to obtain crude proteins (Supplementary Fig. S1c). The clear protein extract was pipetted into a Millipore 15 mL 10K filter, which was centrifuged at 4000 rpm and 4°C. Finally, 150 ml protein extract was concentrated to obtain 15 ml for SDS-PAGE (10%) analysis and concentration measurement using Coomassie Brilliant Blue, and then stored in a -20°C freezer for catalytic analysis and protein purification described below.

### In vitro enzymatic assays of native proteins and thin layer chromatography analysis

*In vitro* assays were performed to test the catalysis of native protein extracts from cells and liquid medium. Substrates tested included (-)-epicatechin, (+)-catechin, phloroglucinol, and catechol. Crude protein extracts tested included those from 6R and P3 proteins described above. Buffers were oxygenated by strongly vortexing tubes for 5 min at room temperature immediately prior to enzymatic incubations. Enzymatic reactions were carried out in 500 µl volume contained in a 1.5-ml Eppendorf polyethylene tube, which was composed of 475 µl of buffer oxygenated (0.02 M pH 7.0 or pH 8.0 Tris-HCl buffer or 0.05 M pH 6.0 phosphate-citric buffer), 15 µl protein extracts (10 µg), and 5 µl substrate (0.5 mg). In addition, crude protein extracts were boiled for 1-10 min for enzymatic assay. Accordingly, four types of control reactions were performed, including substrate absence, enzyme absence, BSA as the substitute of enzyme, and 1-10 min boiled crude protein extracts. All reactions were incubated for 30 min at room temperature and stopped by adding 1.0 ml of ethyl acetate (EA). The tubes were then vortexed and centrifuged at 5,000 rpm for 5 min. The EA phase was transferred to a new 1.5 ml tube and completely evaporated in a speed vacuum. The remained residues were dissolved in 200 µl of HPLC-grade methanol for thin layer chromatography (TLC), HPLC, and HPLC-MS analyses described below.

TLC was used to examine enzymatic products. Ten µl of methanol extract was loaded onto an aluminum-backed silica Kieselgel 60 F254 TLC sheet (0.2 mm layer thickness, EM Sciences, USA). Authentic standards of catechin, epicatechin, procyanidin B1, and procyanidin B2 (0.1-1 µg) were loaded onto the same TLC sheet as positive controls. The developing agent freshly prepared was composed of formic acid: water: ethyl acetate (1:1:18, v/v). Samples on TLC plates were separated with the developing agent in a glass chamber. After separation, plates were taken out from the chamber and then dried in the air in a chemical fume hood. Meanwhile, 0.1 % dimethylaminocinnamaldehyde (DMACA) was prepared by dissolving it in ethanol: HCl (50%: 3M). Finally, compounds were visualized by spraying 0.1% DMACA onto TLC plates.

### Native enzyme purification

Based on the catalytic screening results, the crude protein extracts from 6R cells were subjected to three steps of purification to isolate the active enzyme. First, given that the catalytic protein was thermally stable, the crude protein extracts were boiled for one min and then centrifuged at 12,500 rpm for 10 min to remove all thermally instable proteins in the pellets. The resulting supernatant included active enzymes. Second, the supernatants were subjected to a purification via an anion exchanger chromatography. Columns for ion-exchange chromatography were prepared by loading 15 ml DEAE Sephadex A-20-120 into a 25 ml syringe, washed 5 times with 15 ml ddH2O, and then equilibrated with 0.02 M pH 7.0 Tris-HCl buffer. The supernatant was loaded onto a column, which was washed with 0.02 M pH 7.0 Tris-HCl. After the column was washed additional three times (each with 15 ml) with this buffer, proteins were eluted with three types of elution buffers, 0.02 M pH 7.0 Tris-HCl buffer added with 0 M (buffer I), 0.1 M (buffer II) and 0.2 M NaCl (buffer III). The elution started with buffer I, followed by II, and III. Each elution buffer was used to elute the column three times and the buffer volume used at each time was equal the column volume. As a result, nine elution fractions were collected, each of which was tested for catalytic activity as described above. The most active fractions were combined for further purification. Third, this step was gel chromatography (size-exclusion) purification. Columns were prepared by loading 15 ml Sephadex G-75 into a 25-ml syringe and then washed with 45 ml 0.02 M Tris-Cl pH 7.0 buffer. The active fractions from step 2 were loaded onto a G-75 column and then eluted with 0.02 M Tris-Cl pH 7.0 buffer. The elution was collected every 30 seconds to obtain different fractions. All fractions were analyzed by a 10% SDS-PAGE. All single band fractions with the same molecular weight were combined and concentrated to 5 ml using a Millipore 15 ml 10K filter as described above. Each purified enzyme fraction was quantified with the Bio-Rad Bradford protein assay kit and examined using a 10% SDS-PAGE to check the relative purity. The bands from SDS-PAGE were cut and then collected to a clean 1.5 ml tube for protein sequencing at Duke Center for Genomics and Computational Biology (https://genome.duke.edu/cores-and-services/proteomics-and-metabolomics). Sequencing steps followed the center’s protocol. Finally, all purified proteins were stored at -80°C for enzymatic assays.

### In situ catalytic test on native-polyacrylamide gel and nomenclature of flavanol polymerase

Native-polyacrylamide gel electrophoresis (Native-PAGE) was completed to test enzymatic reactions on gel. Native-polyacrylamide gel (10.0 %) was freshly prepared prior to electrophoresis. Separation gel was prepared using 2.33 ml of 30% acrylamide/bisacrylamide mixture, 1.75 ml of 4× Tris (pH 8.8), 2.85 ml of ddH_2_O, 70 µl of 10% (w/v) ammonium persulphate (APS), and 5 µl of tetramethylethylenediamine (TEMED). Stacking gel was prepared using 500 µl of 30% acrylamide/bisacrylamide mixture, 375 µl of 4 × Tris-HCl (pH 6.8), 2.1 ml of ddH_2_O, 30 µl of 10% APS, and 4.0 µl of TEMED. A comb was used to create 40 µl volume wells for loading samples. Native gel was prepared in a caster (10 x 8 cm), which was then installed in a Bio-Rad Mini Protean® Tetra Cell Electrophoresis apparatus (Bio-Rad, USA). Ten µg of protein sample was mixed with 10 µl of protein loading buffer, which was composed of 50% glycerol, 0.5 M Tris-HCl (pH:7.0), 0.15 µM bromophenol blue (Bio-Rad), and ddH_2_O. In addition, the purified protein was digested with proteinase K for one hour. Bovine serum albumin (BSA) and proteinase K digested protein were used as negative controls. Tris-glycine running buffer (1 x) was prepared with 3.03 g Tris and 14 g glycine for electrophoresis. The polyacrylamide gel and 1-liter running buffer were placed in an electrophoresis chamber. The samples were gently loaded into the wells of gels. In addition, 10 μg of Bio-Rad Precision Plus Protein™ ladder was loaded as a molecular weight standard. The electrodes of the electrophoresis apparatus were wired to a power source. The electrophoresis apparatus was placed in a container with ice to avoid the increase of temperature. The voltage of electrophoresis was set at 70 volts for 30 min and then increased to 120 volts for 90 min. After the completion of electrophoresis, the gel was rinsed using ddH_2_O and 0.02 M pH 7.0 Tris-HCl buffer and then placed in a glass tank for incubation with a substrate. The gel was suspended in 0.02 M pH 7.0 Tris-HCl buffer containing 1 mg/ml (+)-catechin or (-)-epicatechin. The incubation was carried out on a rotary shaker with a speed of 15 rpm. Color changes on the gel were recorded at 0, 30, and 60 minutes after incubation started. The appearance of a yellowish color indicated that proteins had a catalytic activity. Given that the purified protein catalyzed the formation of oligomeric compounds from epicatechin and catechin, herein, we named this enzyme as flavanol polymerase (FP). In addition, given that it was obtained from red tobacco cells, we named it *N. tabacum* FP (NtFP) to indicating its specie origination.

### Cloning of FP cDNA from 6R cells

We cloned *FP* (or *NtFP*) cDNA from red 6R cells. First, the oligomeric peptides obtained from protein sequencing were used to blast against tobacco sequences in the GenBank curated by NCBI. All oligomeric peptides hit one tobacco peroxidase gene (Supplementary Fig. S4). Based on the sequence, a pair of primers was designed to clone its cDNA from our red 6R cells. The forward primer was 5’-GGGGACAAGTTTGTACAAAAAAGCAGGCTTAATGGCTTTTCGTTTGAGTCA-3’, which contained an attB1 recombinant sequence and an ATG start codon. The reverse primer was 5’-GGGGACCACTTTGTACAAGAAAGCTGGGTATCAGTGGTGGTGGTGGTGGTGCATAGAAGCCACAGAGCT-3’, which contained an attB2 recombinant sequence, a 6-His tag encoding sequence, and a TCA stop codon.

Total RNA samples were isolated from 6R calli and other samples using a TRI Reagent™ Solution (ThermoFisher, Catalog number: AM9738) and treated with DNase to remove genomic DNA as we reported recently [42]. We used a high capacity cDNA Reverse Transcription Kit (Applied Biosystems, USA) to reversely transcribe one µg of DNA-free RNA into the 1^st^ strand cDNA in a 20 µl reaction volume by following the manufacturer’s protocol. Two µl of the 1^st^ strand cDNA product of 6R cells was used as the template for PCR to amplify a *NtFP* cDNA. The thermal cycle consisted of 94°C for 5 min, followed by 30 cycles of 94°C for 30 s, 60 °C for 1 min, and 72°C for 30 s. The final extension step was 10 min at 72 °C. The amplified cDNA was then cloned into the gateway pDONR221 vector by using the Gateway BP Clonase II Enzyme Mix (Invitrogen, USA) following the manufacturer’s protocol. The ligated products were introduced into competent *E. coli* ccdB Survival 2 T1^R^ cells via a common heating shock transformation protocol, which were streaked on agar-solidified LB medium supplemented with 50 mg/l antibiotics. Positive colonies were selected and then inoculated to 5 ml liquid LB medium contained in plastic tubes, which were placed on an incubator set up with a speed of 250 rpm and 37°C for overnight. *E. coli* was harvested to isolate the pDONR221-FP plasmid, the steps of which were as we reported previously [59].

### Expression of recombinant cDNA in E. coli and purification

A recombinant FP protein was induced for enzymatic assays. A pair of primers was designed to include *GST* and *His*-tag sequences in the 5’- and 3’-end of *FP*, respectively (Supplementary Table 1). In addition, nucleotides were added at both ends for Gateway cloning. The recombinant pDONR221-FP plasmid was used as the template for PCR to amplify a recombinant cDNA including *GST-FP-His-tag* sequences. After the accuracy of *GST-FP-His-tag* sequence was conformed, it was cloned to the pDEST15 protein expression vector by using Gateway LR Clonase II Enzyme Mix (Invitrogen, USA) by following the manufacturer’s protocol. The ligation product was introduced into competent BL21 (DE3) plysS *E. coli* strain, which was inoculated on agar-solidified LB medium supplemented with 200 mg/l ampicillin and 34 mg/l chloramphenicol. Antibiotics-resistant colonies were screened to select positive colonies as we reported previously [59]. One positive colony was used to isolate the positive recombinant pDEST15-FP-His vector as described above and then used to induce recombinant GST-FP-His protein.

A positive colony was inoculated into 10 ml of liquid LB medium supplemented with 200 mg/L ampicillin and 34 mg/L chloramphenicol in a 50 ml tube. The tube was placed in a rotatory incubator and shaken at 200 rpm for 16 hrs under 37°C. When its OD value at 600 nm reached 1.0, 1.0 ml of suspension culture was inoculated into 100 ml of liquid LB medium containing 200 mg/l ampicillin and 34 mg/l chloramphenicol in a 250 ml Erlenmeyer (E) flask. The E-flask was placed on an incubator, which was shaken at a speed of 120 rpm and at 37 °C until its absorption measured at 600 nm reached 0.7. Then, the culture temperature was reduced to 30°C and isopropyl-1-thiol-beta-D-galactopyranoside (IPTG) was added to the cultures to a final concentration of 1.0 mM. After 4 hrs of induction, the suspension cultures were transferred to 200 ml tubes, which were centrifuged at 6,000 rpm for 5 min. The remained pellets were frozen in liquid nitrogen and then stored at -80°C until the isolation of the recombinant protein.

The *E. coli* pellets were completely suspended in 10 ml of denaturing extraction buffer (8 M urea, 20 mM Tris-Cl pH 8.0, 10 mM imidazole) and then lysed for 5 min with supersonic sound. The resulting lysate mixture was centrifuged at 12,000 rpm and 4°C for 20 min. The supernatant containing crude proteins was transferred to a new tube for recombinant protein purification.

The purification of recombinant protein was carried out on a Ni-NTA column (G-Biosciences, USA). One ml of Ni-NTA Sefinose resin was loaded onto a plastic column. After the resin was stacked tightly, the column was washed three times with ten volumes of denaturing extraction buffer. Then, 10 ml of crude protein extracts were loaded onto the top of the resin column and eluted by gravity at a flow rate of 0.5 ml/min. The column was washed three times using four times the volume of the denature extraction buffer to wash off non-specific column-binding proteins. Then, the column was placed in a cold room and washed three times using ten times the volumes of refolding buffer (20 mM Tris-Cl pH 8.0, 10 mM imidazole), which refolded the recombinant protein on the column. The refolded recombinant protein was then eluted with 3 ml of elution buffer (20 mM Tris-Cl pH8.0, 250 mM imidazole) to obtain recombinant GST-FP-His protein. The quality of the purified recombinant protein was examined by electrophoresis on a 12% SDS-PAGE as described above.

### Enzymatic assays of recombinant FP

The enzymatic assay of the recombinant protein followed the same protocol for its native protein described above. In brief, the enzymatic reaction was composed of 475 µl 0.02 M Tris-HCl (pH 7.0 or pH8.0) buffer or 0.05 M pH 6.0 phosphate-citric buffer, 0.5 mg substrates, and 10 µg recombinant protein in a total volume of 500 µl in 1.5 ml tubes. To test the requirement of oxygen, 0.5 µl 30% hydrogen peroxide was added to reactions. In addition, reactions without substrates, without protein, or with BSA as a substitute of the enzyme were performed as controls. All reactions were incubated at 37 °C for 30 min and then stopped by adding one ml of ethyl acetate (EA). The tubes were vortexed and centrifuged with a speed of 5,000 rpm at 4°C for 5 min. The EA phase was transferred to new 1.5 ml tubes and completely evaporated in a speed vacuum. The residues were dissolved in 200 µl of methanol for HPLC-qTOF-MS/MS analysis.

Enzymatic assays were performed to optimize reaction time, pH, and temperature, and then to characterize enzymatic kinetics of FP in the optimized buffer that was oxygenated with 0.5 µl 30% H_2_O_2_. First, to optimize reaction time, each enzymatic reaction was composed of 475 µl of 0.02 M Tris-Cl (pH 7.0) buffer, 0.5 mg (-)-epicatechin, and 10 µg of recombinant protein. The reaction times were 30 min, 45 min, 60 min, 90 min, 120 min, and 180 min. The velocity of reactions was calculated based on the decrease of (-)-epicatechin. Second, to optimize pH value, both 0.05 M phosphate sodium buffer and 0.1 M Tris-HCl buffer were tested. Each reaction was composed of 475 µl phosphate sodium buffer (pH 6.2 to 8.0) or Tris-HCl buffer (pH 7.0 to 9.5), 0.5 mg (-)-epicatechin, and 10 µg of recombinant protein. All reactions were performed at 37 °C for 30 min. Third, to optimize temperature value, each enzymatic reaction consisted of 475 µl of 0.02 M Tris-Cl (pH 7.0) buffer, 0.5 mg (-)-epicatechin, and 10 µg of recombinant protein and incubated for 30 min. The reaction temperatures tested included 35, 40, 45, 50, 60, 70, and 80 °C. Forth, to calculate kinetic parameters, reactions were composed of 475 µl of 0.02 M Tris-Cl (pH 7.0) buffer, 10 µg of recombinant protein, and 0.312, 0.5, 0.625, 0.825, 1, 1.2, 1.4, or 1.7 µM of (-)-epicatechin. The reactions were performed for 30 min at 37 °C. All reactions were stopped by adding EA to extract metabolites as described above. The enzymatic products were analyzed by LC-qTOF-MS/MS.

### Purification of enzymatic products

EP-P1 was one of main products formed from the enzymatic reactions with epicatechin as substrate. We purified it via three steps. First, a 1000 ml-scaled reaction with 100 mg epicatechin substrate was developed to produce products. The enzymatic reactions were composed of 5 mg crude protein extract of 6R cells, 100 mg (-)-epicatechin, 500 ml 0.05 M phosphate-citric buffer (pH 6.0), and 0.5 ml 30% hydrogen peroxide in a 1000 ml Erlenmeyer flask. The reactions were performed at room temperature for 30 min and stopped by adding 1 liter of EA to extract products as described above. The EA phase was transferred to a clean 500 ml round flask for evaporation with a rotatory evaporator at room temperature. Second, the remained residue was dissolved in 2 ml EA and loaded onto a few TLC silica gel 60 plates (2 mm thickness, 20 cm × 20 cm, EMD Millipore, USA). The TLC plates were developed with a solvent mixture consisting of water: formic acid: ethyl acetate (1:1:18), v/v) in a glass chamber. After the full development of plates, yellowish silica bands were collected and those bands with the same Rf value were pooled together for further purification. The resulting silica gel mixtures containing yellowish EP-P1 were grounded into a fine powder, which was completely suspended in 10 ml ddH_2_O in a 50 ml tube. Then, 25 ml EA was added to the mixture and the tube was vortexed for 1 min. The mixture was centrifuged at 3,000 rpm for 5 min and the EA phase was transferred to a new 50 ml polypropylene tube. Last, the EA extracts was dried with a nitrogen gas stream at the room temperature. The remaining residue was dissolved in 1 ml methanol for further purification with HPLC on Shimadzu LCMS-2010 EV instrument. In brief, compounds were separated on an Advantage 300 C18 column (250 × 10 mm, 5µm, Thomason Liquid Chromatography, USA). The mobile phase solvents used for elution included 1% acetic acid in water (solvent A, HPLC grade acetic acid, HPLC grade water) and 100% acetonitrile (solvent B, HPLC grade). A gradient solvent system, which was developed to separate compounds, was composed of ratios of solvent A to B: 80:20 (0–5 min), 80:20 to 70:30 (5–10 min), 70:30 to 55:45 (10–20 min), 55:45 to 50:50 (20–30 min), 50:50 to 45:55 (30–35 min), 45:55 to 80:20 (35-36 min). After the last gradient step, the column was equilibrated and washed for 10 min with a solvent ratio of A: B 80:20. The flow rate was 0.6 ml/min and the injection volume was 50 µl. Chromatograms were recorded at 280 nm and 380 nm. All elutes were collected to different tubes labelled with numbers. The same elutes collected from different separations were combined and extracted with 50 ml EA. The EA phase was transferred to new tubes and dried with a nitrogen gas stream to obtain purified compounds. A small amount of the compound was dissolved in 1 ml methanol and then the purity was examined by LC-qTOF-MS/MS. The remained major part was used for NMR analysis and physical and chemical characterization described below.

### Butanol: HCl cleavage of enzymatic products

Butanol: HCl (50:50) boiling is a classic method to cleave and measure PAs. As described previously [44], we used this method to cleave enzymatic products. In addition, PB2 was used as a positive control. In brief, 5 µl PB2 (50 µg) or a purified product (50 µg EP-P1 or EP-P2) was mixed with 950 µl butanol: HCl in a 1.5 ml tube. The mixture was boiled for 1 hr and then cooled down to the room temperature. The boiled mixture was used to measure absorbance at 550 nm. After measurement, the samples were evaporated to remove butanol and water in a speed vacuum. The remaining residue was suspended in 200 µl 0.1% HCl in methanol for LC-qTOF-MS/MS assay.

### Acid-catalytic degradation and phloroglucinol competition assays

The acid-catalytic and phloroglucinol-based degradation (phloroglucinolysis) is a classic method to determine extension units of PAs. The reaction was composed of 450 µl of 0.1 N HCl in methanol, 50 mg/ml phloroglucinol, 10 mg/ml ascorbic acid, and 50 µl of 1 mg/ml purified products (EP-P1 or CA-P1) in a 1.5 ml tube. Then, the reaction tubes were incubated at 50 °C for 20 min. After cooled to the room temperature, the reaction products were examined with LC-qTOF-MS/MS. In addition, 50 µl of 1 mg/ml procyanidin B2 and procyanidin A2 were used as controls.

### Competition assays between phloroglucinol and epicatechin

The phloroglucinol and epicatechin competition experiment was carried out in a 500 µl volume contained in a 1.5 ml tube. The reaction was composed of 479.5 µl 0.05 phosphate-citric buffer (pH6.0), 5 µl (-)-epicatechin (100 µg/µl), 5 µl phloroglucinol (100 µg/µl), 10 µl native enzyme or BSA (10 µg/µl), and 0.5 µl 30% hydrogen peroxide. In addition, taxifolin and quercetin (100 µg/µl) were used to substitute epicatechin as controls. The tube was incubated at the room temperature for 30 min and then added 1 ml of EA to stop the reaction. The tubes were vortexed three times each for 2 min, followed by centrifugation at 5,000 rpm for 5 min. The EA phase was transferred to a new 1.5 ml tube, which was placed in a speed vacuum for evaporation. The remaining residues were dissolved in 200 µl methanol for HPLC-MS analysis.

### Overexpression of FP in PAP1-ANR tobacco and PA assay

The stop codon of the *FP* ORF was removed by PCR (Because of *N. tabacum* origin, herein, we used *NtFP* to represent *FP* to define its species origin). The resulting *NtFP* fragment was cloned to the Pdonr223 vector using a Gateway BP Clonase II Enzyme Mix (Invitrogen, USA) by following the manufacturer’s protocol. This cloning obtained a recombinant Pdonr223-NtFP vector. The purified Pdonr223-NtFP plasmid was used to clone *NtFP* into the binary vector PMDC84 with Gateway LR Clonase II Enzyme Mix (Invitrogen, USA) by following the manufacturer’s protocol. This cloning generated a recombinant binary vector, PMBC84-NtFP, in which *NtFP* was fused to the 5’-end of *GFP* to obtain *NtFP-GFP*, which was driven by a 35S promoter (Supplementary Fig. S6a). Both PMDC84-NtFP and PMDC84 vectors were introduced into *Agrobacterium tumefaciens* strain GV3101 for genetic transformation. The PAP1-ANR(BAN) tobacco [44] was used for genetic transformation. MS medium, plant hormones used, and all steps of transformation and screening of transgenic plants followed our protocols reported previously [44]. The PAP1-BAN-NtFP and vector control PAP1-BAN-PMDC84 transgenic plants were obtained on the selection medium supplemented with 50 mg/L hygromycin and further confirmed with PCR and RT-PCR. Fifteen PAP1-BAN-NtFP and 15 PAP1-BAN-PMDC84 transgenic plants, five PAP1 plants, and five PAP1-ANR control plants were grown in pot soil and placed side by side in the greenhouse to develop flowers. Seeds were collected from T0 plants and germinated on MS medium containing 50 mg/L hygromycin. Resistant T1 seedlings were grown in the greenhouse to develop flowers for analysis of papanridins, flavan-3-ols, and PA as described above.

### Plasmolysis and subcellular localization of NtFP

The seeds of T1 PAP1-BAN-NtFP and PAP1-BAN-PMDC84 transgenic plants were germinated on MS medium containing 50 mg/L hygromycin for five days to obtain young seedlings. Half of the transgenic seedlings were pretreated with 30% sucrose for 10 min to create plasmolysis of seedlings. In addition, wild type seedlings pretreated with BCECF fluorescent dye were used as positive control of plasmolysis, which followed a classic protocol reported in the literature. In brief, five-day old wild type tobacco seedlings were placed in a perfusion solution (0.2 mM CaCl_2_, 50 mM glucose, 3µM BCECF ∼AM, and 10 mM MES buffer, pH 6.0) for 30 min at the room temperature, and then washed three times with 0.1 mM CaCl_2._ The seedlings were held in the perfusion solution without BCECF∼AM for 1 hour. All seedlings were examined under a confocal laser-scanning microscope (Leica TCS SP2, Leica, Germany). The light wavelength was set at 488 nm to examine the fluorescent signal of GFP.

### Extraction of papanridins, flavan-3-ols, and proanthocyanidins

Frozen samples were ground into fine powder in liquid nitrogen. Two hundred milligrams of powder were weighed into a 1.5 ml tube and then suspended in 1.0 ml 70% acetone. The tube was vortexed 30 seconds, followed by centrifugation for 5 min at 5,000 rpm. The supernatant was pipetted into a new 1.5 ml tube. Acetone was evaporated in a speed vacuum until the water phase left in the bottom of tubes. The remained water phase was added with 500 µl water followed by adding 200 µl chloroform. The mixture was then vortexed and centrifuged at 10,000 rpm to gain the upper water and bottom chloroform phases. The chloroform phase was pipetted to a dispensable container to remove chlorophyll, lipids, and fatty acids. This step was repeated once. The water phase in the tube was added with 1 ml EA and vortexed for 1 min. The tube was centrifuged at 10,000 rpm for 5 min. The EA phase was then transferred into a new 1.5 ml tube and evaporated completely in a Savant SpeedVac vacuum. The remaining residues were dissolved in 200 µl of methanol for HPLC-qTOF-MS/MS assay. Three biological samples for each plant species or cultivar were used to extract these metabolites. PB2 and EC were used to develop standard curves. The contents of EC-P1 and EC-P2 were estimated as equivalent values with the PB2 as standard. The content of EC was estimated using its standard curve. Butanol-HCl boiling described above was performed to estimate the total amount of PAs. Standard deviation was calculated. Student T-test was performed to evaluate statistical differences (P-value less than 0.05, significantly).

### LC-qTOF-MS/MS analysis

HPLC-qTOF-MS/MS analysis was performed to profile the enzymatic products according to our protocol reported previously [47]. HPLC-qTOF-MS/MS analysis was carried out on Agilent 6520 time-of-flight-MS/MS instrument (Agilent Technologies, Santa Clara, CA, USA). The optimized mobile phase solvents included 1% acetic acid in water (solvent A: 1% HPLC grade acetic acid in LC-MS grade water) and 100% acetonitrile (solvent B) (LC-MS grade). A gradient elution system was optimized with the two mobile solvents to separate metabolites on an Eclipse XDB-C18 analytical column (250 × 4.6 mm, 5 µM, 25 ◦C, Agilent). The gradient solvent system was composed of ratios of solvent A to B: 95:5 (0–5 min), 95:5 to 90:10 (5–10 min), 90:10 to 85:15 (10–15 min), 85:15 to 45:55 (15–45 min), 45:55 to 25:75 (45–50 min), 25:75 to 95:5 (50–60 min). After the last gradient step, the column was equilibrated and washed for 10 min with solvents A: B (95:5). The flow rate was 0.4 ml/min. The injection volume of the samples was set up at 5.0 µl. The drying gas flow and the nebulizer pressure were set at 12 l/min and 50 psi, respectively. Metabolites were ionized with the negative mode. The mass spectra were scanned from 100 to 3000 m/z. The acquisition rate was three spectra per second. Other optimized MS conditions included fragmentor: 150 V, skimmer: 65 V, OCT 1 RF Vpp: 750 V, and collision energy: 30 psi. The authentic (-)-epicatechin, (+)-catechin, procyanidin B2, procyanidin A2 were used as the references.

### Nuclear magnetic Resonance (NMR) Analysis

The NMR analysis of the sample EC-P1 (∼2 mg) in 600 µl of acetone-d6 was conducted on a Bruker NEO 600 MHz spectrometer equipped with a Broadband Inverse Probe in METRIC (https://research.ncsu.edu/metric/). The authentic PB2 standard was used as a reference for structural determination.1D ^1^H NMR spectra were recorded at various temperatures ranging from 240K to 300K with 10K per increment. The ^1^H chemical shift and structure conformation were assigned by using rotating frame nuclear overhauser enhancement spectroscopy (ROESY) at 240K. The ^13^C resonance was assigned by heteronuclear multiple bond correlation (HMBC) experiments performed at 240K.

### Phylogeny analysis

A BLAST search obtained 37 homologues of NtFP amino acid sequences from the GenBank curated by NCBI (https://blast.ncbi.nlm.nih.gov/Blast.cgi). An alignment was completed using the online tool Clustal Omega (https://www.ebi.ac.uk/Tools/msa/clustalo/). The alignment further generated a phylogenic tree by using NGphylogeny.fr tool (https://ngphylogeny.fr/).

## Acknowledgements

We are thankful to Dr. Hanna Gracz for her advice and discussion in NMR data analysis and assignment of protons and carbons. We are appreciative of Dr. Peter Thompson for his discussion in assignment of protons and carbons. We are grateful to Dr. Richard Dixon from University of North Texas for his critical reading and suggestions.

## Author Contributions

Conceptualization: DYX

Methodology: YZ, SY, XS, DYX

Investigation: YZ, SY, XS, DYX

Funding acquisition: DYX

Project administration: DYX

Supervision: DYX

Writing – original draft: YZ, SY, DYX

Writing – review & editing: YZ, SY, XS, DYX

## Completing interests

Authors declare that they have no competing interests.

## Data and materials availability

All data are available in the main text or the supplementary materials.

---

**Figures S1-S4 Native protein purification and activity (Figure 2)**

**Fig. S1.**
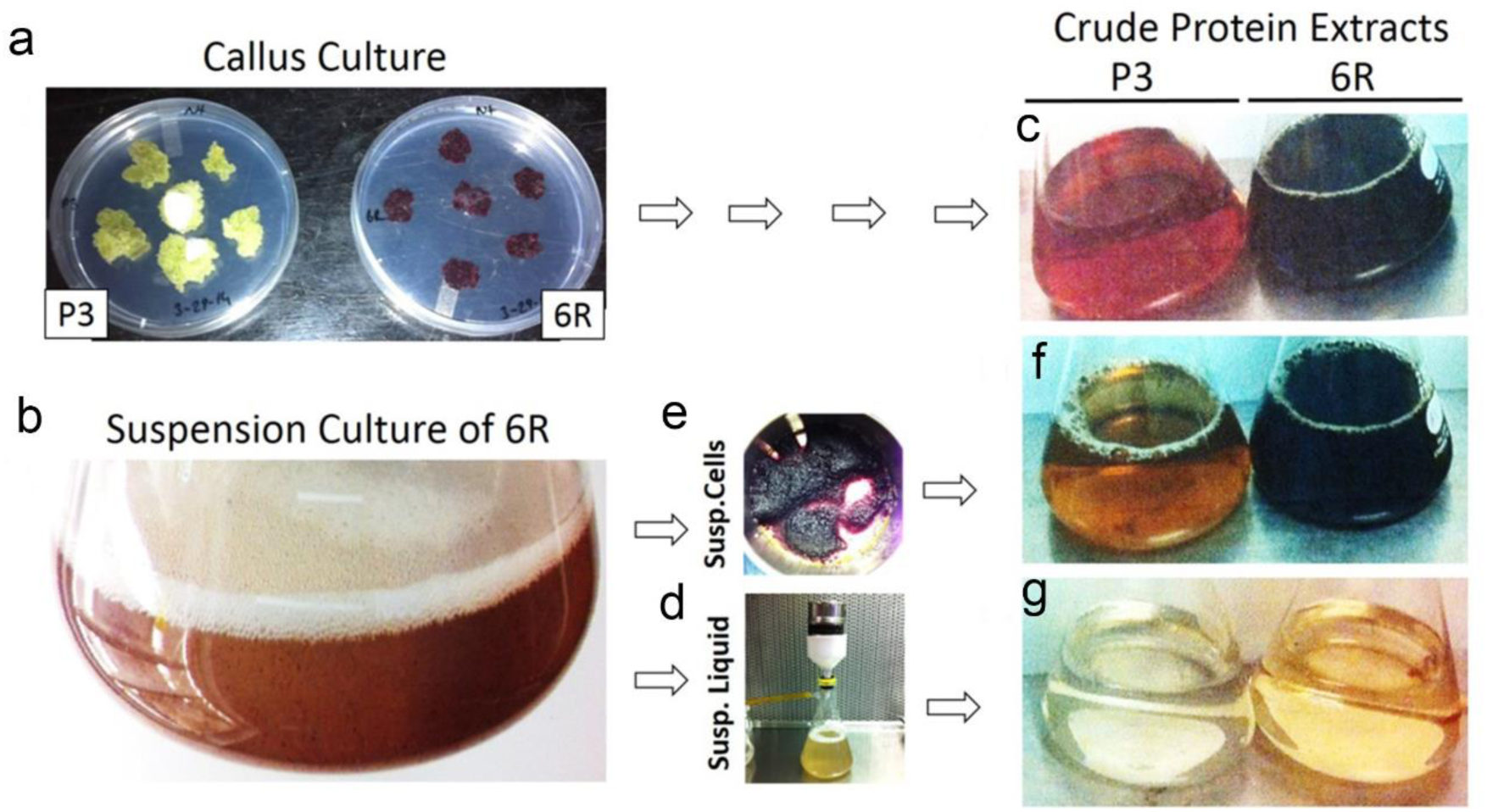
Callus and suspension culture of red and wild type cells for protein extraction. **a,** Callus culture of wild type tobacco (P3) and PAP1 transgenic red tobacco (6R) cells were grown on the subculture media. **b,** Suspension culture of red 6R tobacco cells were grown in liquid medium. **c,** Crude protein solutions were extracted from P3 and 6R calli, **d,** Liquid medium and suspension cells were harvested from suspension culture. **e,** Suspension 6R cells were harvested. **f,** Crude proteins were extracted from P3 and red 6R suspension cells. **g,** Crude proteins were extracted from P3 and red 6R liquid media.

**Fig. S2.**
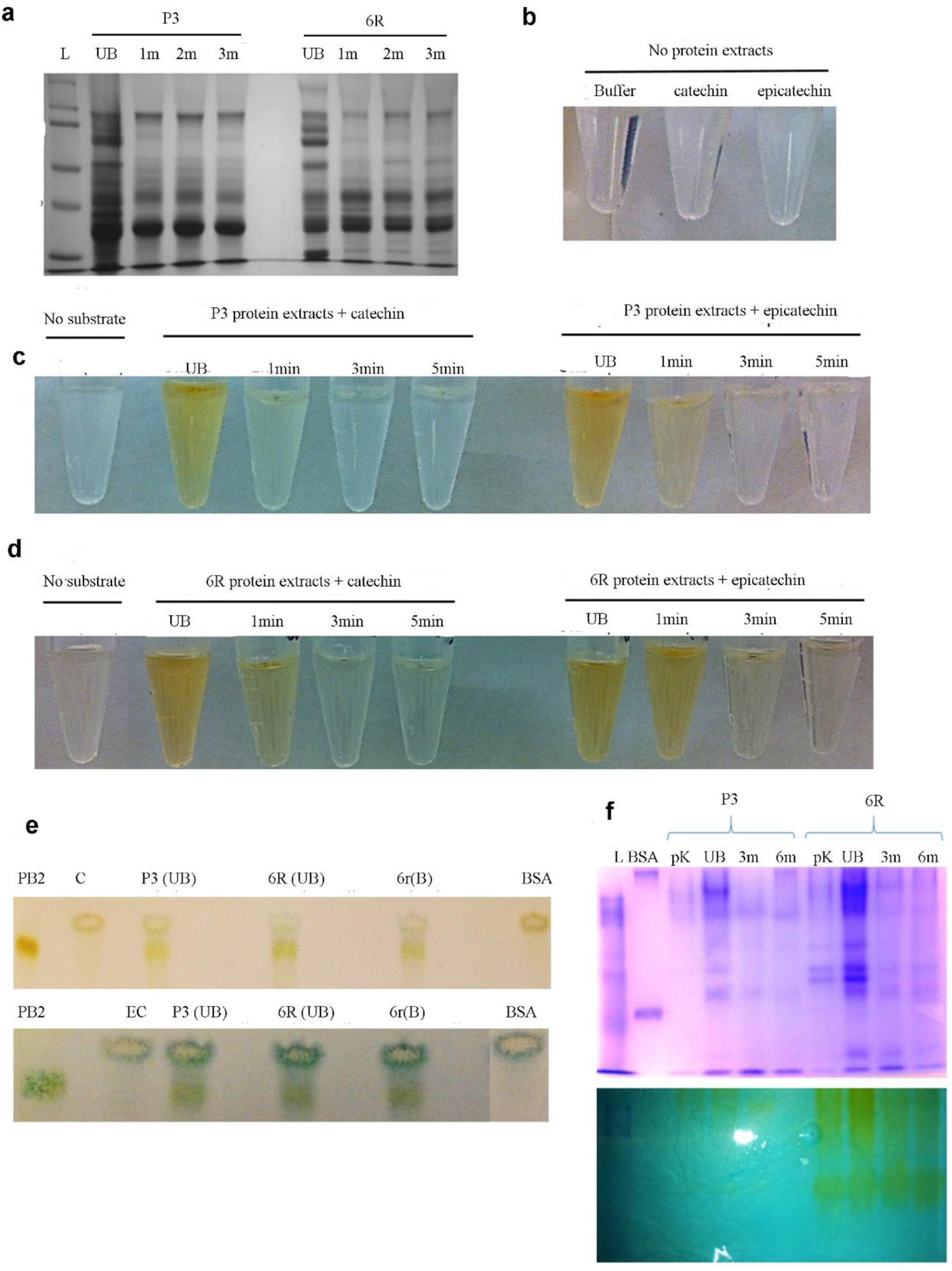
Crude proteins extracted from 6R and P3 cell suspension cultures and incubation with substrates. **a,** A SDS-PAGE image shows different profiles of crude proteins extracted from 6R (shown in Figure 2 too) and P3 cell suspension cultures and their corresponding boiled proteins. L, protein ladder (kDa), UnB: Un-boiled crude protein extracts, and 1-3m: proteins after crude protein extracts were boiled for 1, 2, and 3 minutes at 100°C. **b-d,** Incubations of crude protein extracts with catechin or epicatechin as substrate produce new yellowish or brownish-yellowish compounds. Crude proteins used here include un-boiled and boiled (1, 3, and 5 min, labeled as 1m, 3m, and 5m). **b,** buffer and two substrates in the buffer are colorless. **c;** un-boiled and 1 min-boiled P3 crude proteins converted catechin and epicatechin to yellowish or light brownish compounds. **d,** un-boiled and 1 and 3 min-boiled 6R crude proteins converted catechin and epicatechin to yellowish or light brownish compounds. **e,** TLC and DMACA staining showed PA-like oligomers formed from *in vitro* incubations of crude protein extracts of 6R and P3 suspension cells with (+)-catechin or (-)-epicatechin. Both crude 6R protein extracts and 1 min-boiled proteins were tested for catalytic activity. Bovine Serum Albumin (BSA) was used as protein control. In addition, the negative (-) labelling denoted controls that only proteins were used in the incubations without substrates added. Standards include procyanidin B2 (PB2), (+)-catechin (C) and (-)-epicatechin (EC). 6R(UB): un-boiled 6R proteins, 6r(B): 1 min boiled 6R protein, and P3(UB): un-boiled P3 proteins. **f,** Native-PAGE images show the protein band profiles from un-boiled, 3 and 6 min-boiled, and protease K digested crude proteins of P3 and 6R. A few gels were stained with Coomassie blue to show protein band profiles, while the others were incubated with 100 µM epicatechin to show *in situ* reaction. The incubations with epicatechin not only demonstrated that crude proteins converted epicatechin to yellowish compounds, but also revealed the catalytic differences between P3 and 6R proteins. L: protein ladder (kDa) and pK: proteins digested by proteinase K.

**Fig. S3.**
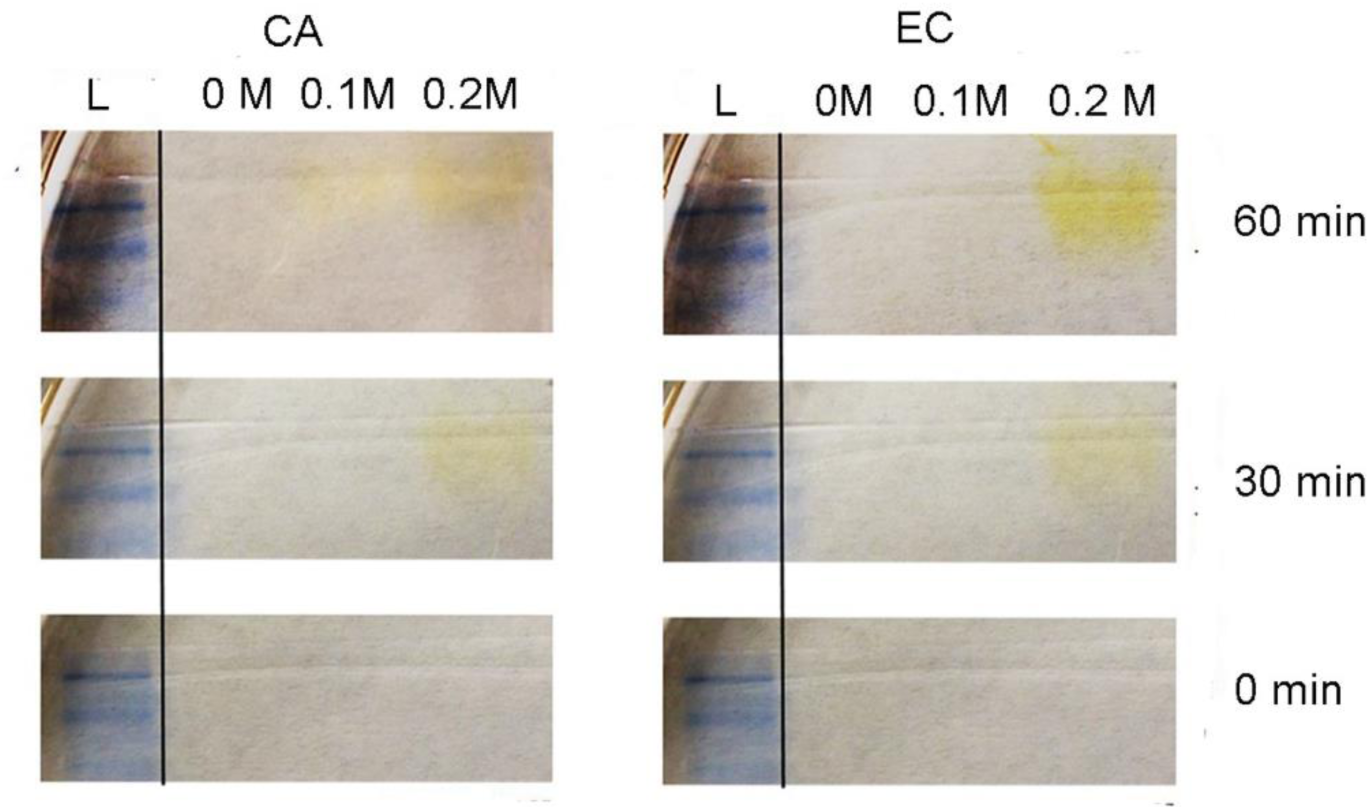
Protein fractions isolated from ion-exchange columns (IECs) and examination of their activity via an incubation of native-PAGE gels and substrates. Proteins were eluted through IECs and loaded onto a Native-PAGE with buffer supplemented with 0, 0.1 M, and 0.2 M NaCl. L; protein ladder (kDa). Native PAGE gel images show the dynamics of yellowing products formed on the gels after incubated with catechin and epicatechin for 0, 30 and 60 min. These data demonstrate the presence of an active enzyme.

**Fig. S4.**
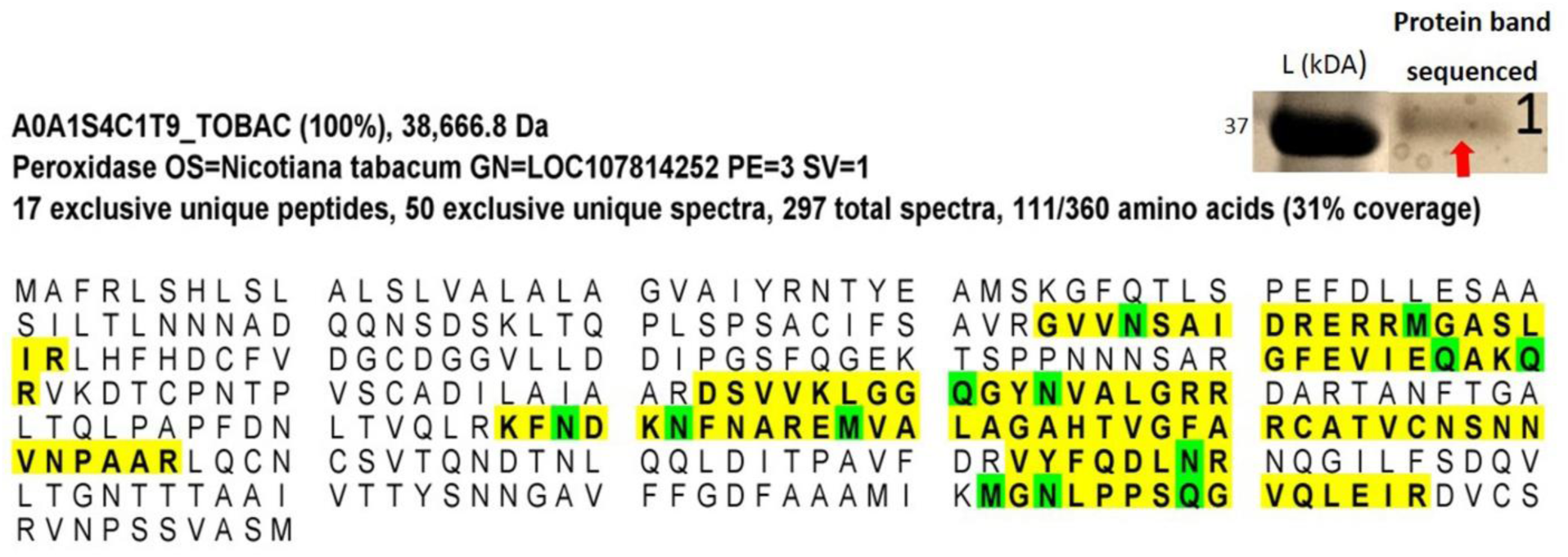
Peptide sequences of an active protein fraction and blasting results. The amino acid sequences of six peptides highlighted with a yellow background were obtained from proteomics. The blast search with the six peptides hit one protein in the GenBank, which was annotated to be a peroxidase.

**Figures S5-S7 support cloning and functional characterization of FP (Figure 3)**

**Fig. S5.**
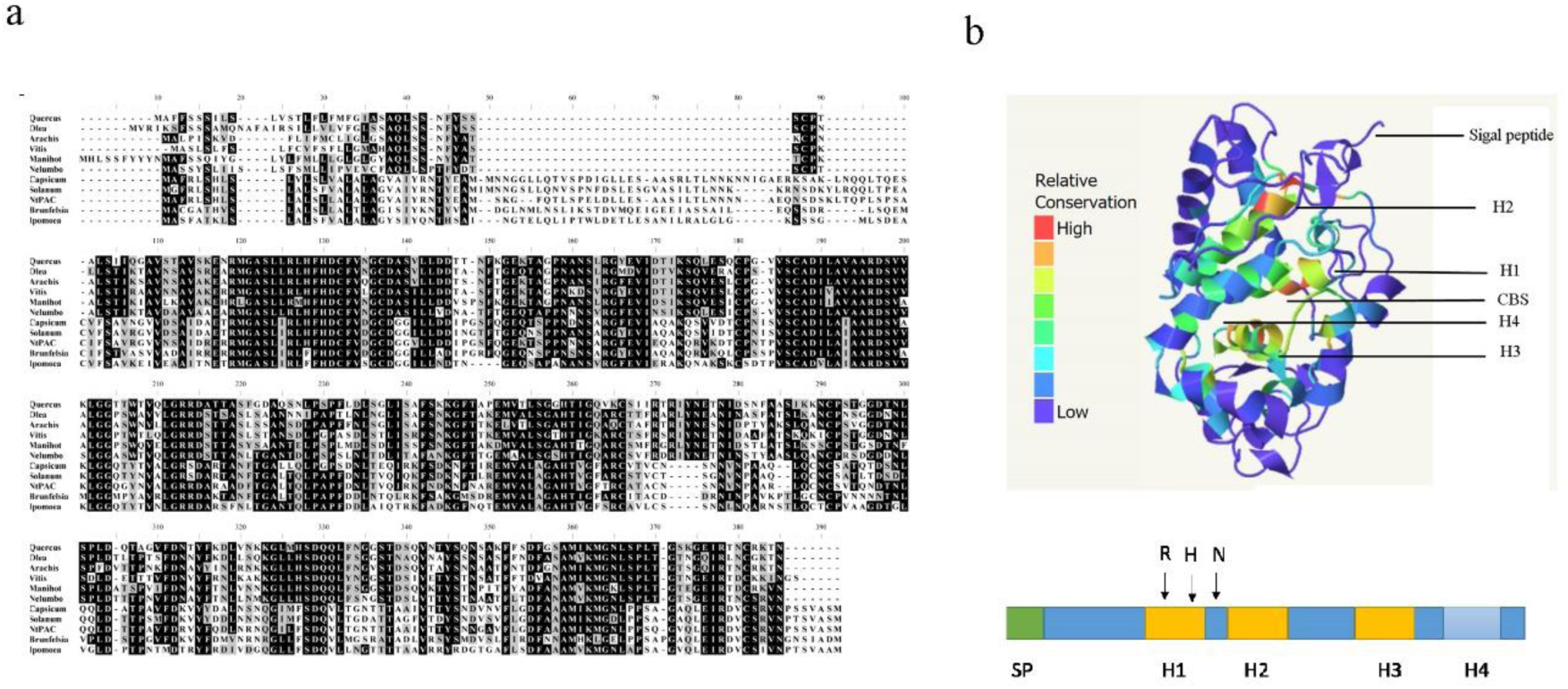
An alignment of amino acid sequences from 11 FP homologs and 3D modeling of FP. **a,** a sequence alignment was developed using amino acid sequences of FP and 10 homologs. **b,** a 3D model of FP was predicted based on a peanut peroxidase as the template. SP: signal peptide, H1, H2, H3, and H4: helix. CBS: calcium binding sites including R, H, and N residues.

**Fig. S6.**
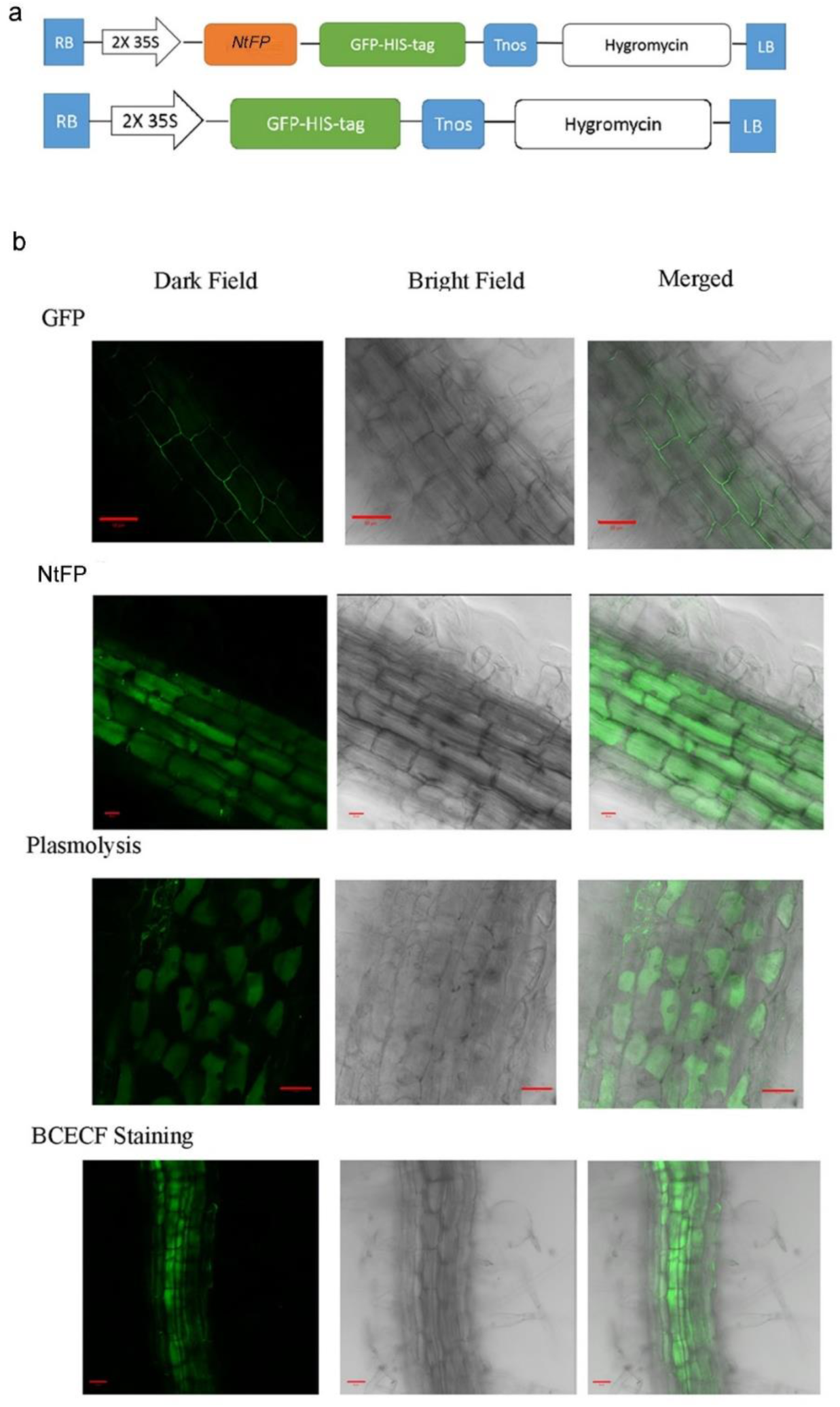
Constructs and subcellular localization of FP (NtFP) in stably transgenic PAP1-ANR (BAN)-FP plants created from the transformation of PAP1-ANR (BAN) tobacco. **a,** Two T-DNA cassettes were developed for genetic transformation, the T-DNA cassette of the PMDC84 binary vector and the T-DNA cassette of a recombinant binary vector developed from PMDC84 by fusing *NtFP* to the 5’-end of *GFP* gene for subcellular localization. These two T-DNA cassettes were transformed into PAP1-ANR plants. PMDC84 was used as a vector control. **b,** Confocal microscopy images showed the vacuolar localization of the GFP fluorescence signal of NtFP-GFP in root cells of transgenic PAP1-ANR-FP tobacco seedlings. GFP: GFP control; NtFP: fused NtFP-GFP in vacuoles; plasmolysis: fused NtFP-GFP in vacuoles; BCECF staining: positive control of vacuolar localization.

**Fig. S7.**
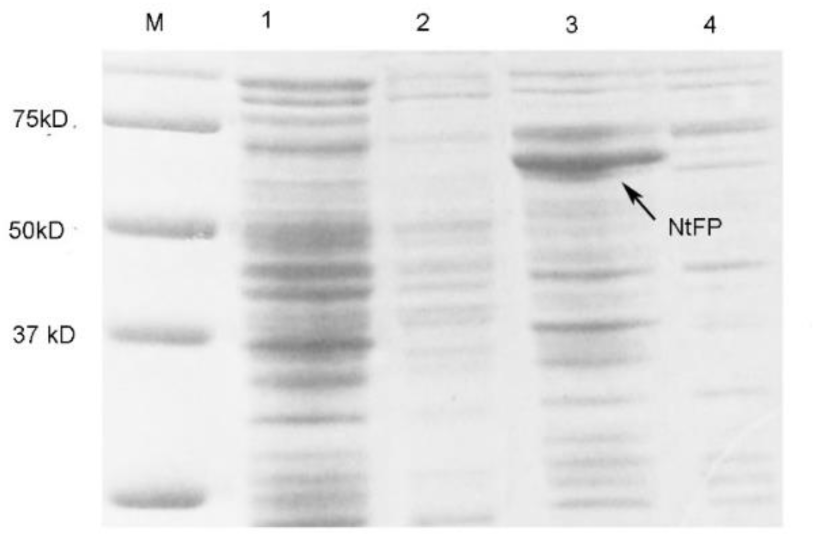
A SDS image showing recombinant flavanol polymerase (FP or NtFP) in the inclusion body heterogeneously induced in *E. coli*. M: protein ladder marker, 1: proteins induced from *E. coli* with a control vector by IPTG, 2: proteins expressed in *E. coli* with an NtFP vector without IPTG induction, 3: proteins in the inclusion body induced from *E. coli* with an NtFP vector by IPTG, and 4: soluble proteins induced from *E. coli* with an NtFP vector by IPTG.

**Figures S8-S11 support enzymatic characterization**

**Fig. S8.**
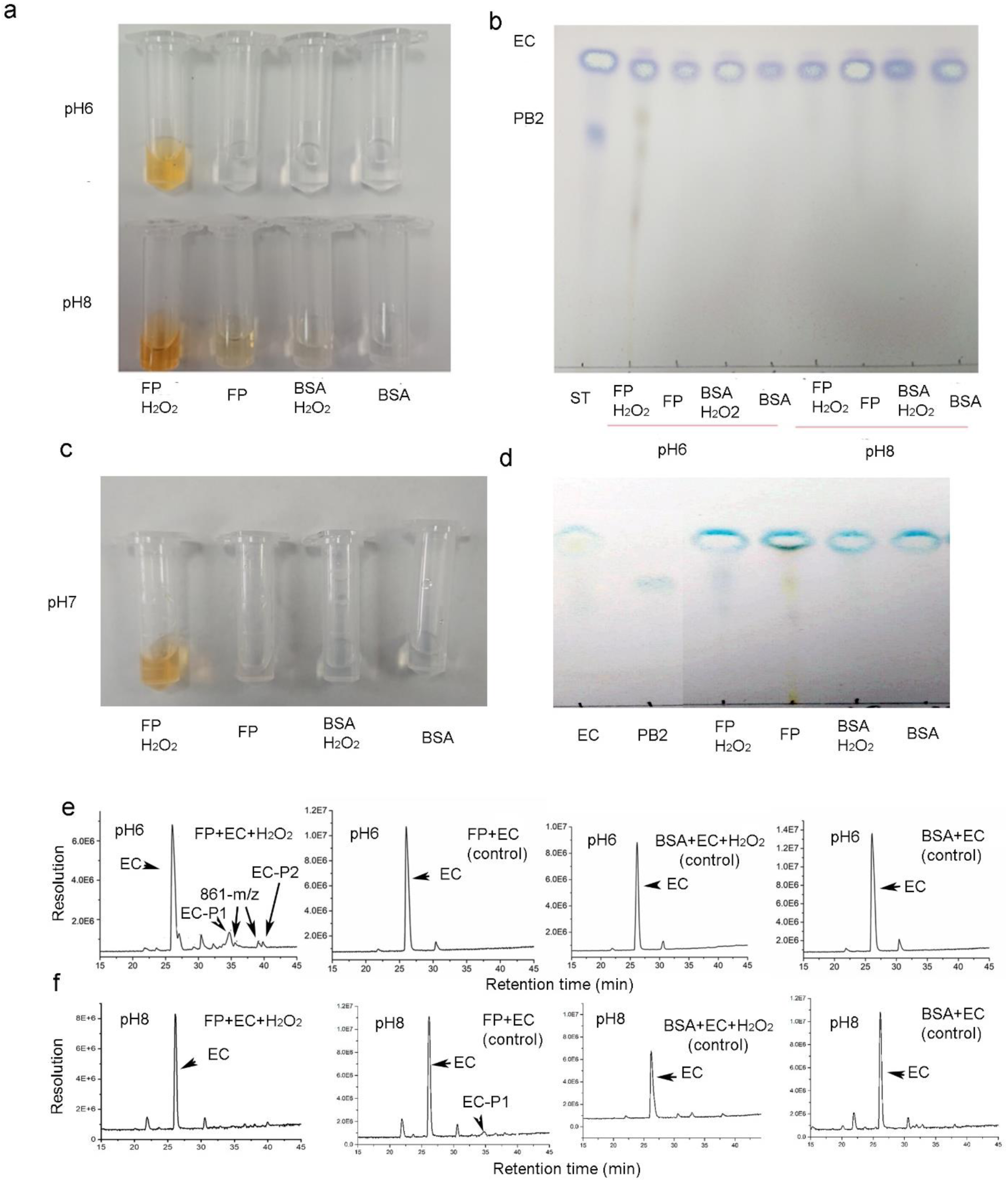
Effects of hydrogen peroxide and three pH buffers on the formation of yellowish compounds in enzymatic reactions consisting of FP and (-)-epicatechin. **a,** Effects of pH6 and pH8 as well as hydrogen peroxide on yellow color formation from the *in vitro* incubation consisting of native FP and epicatechin. Buffers were not vortexed for oxygenation prior to adding enzyme and epicatechin. In the pH6 buffer, only the incubation with FP and hydrogen peroxide (FP+H_2_O_2_) converted epicatechin to yellowish compound(s). In the pH8 buffer, except for the control with BSA protein only (BSA), other incubations (FP+H_2_O_2_, FP, and BSA+H_2_O_2_) converted epicatechin to light yellowish to slightly brownish compounds. These results indicate that in the pH6 buffer, hydrogen peroxide is required for the catalysis to produce yellowish compounds, while, in the pH8 buffer, hydrogen peroxide enhances producing yellowish and brownish compounds. **b,** TLC analysis showed that in the pH6 buffer, PA-like products were formed from the incubation of FP, epicatechin, and hydrogen peroxide (FP+H_2_O_2_) but were hardly detected from three control conditions (FP, BSA+H_2_O_2,_ and BSA). In the pH8 buffer, PA-like compounds were hardly detected from the incubation of FP, epicatechin, and hydrogen peroxide (FP+H_2_O_2_) and three control incubations. The incubations were extracted with ethyl acetate, which was evaporated. The residues were dissolved in methanol for TLC. TLC plates were sprayed with DMACA reagents. **c-d,** hydrogen peroxide promoted the formation of yellowing compound in the incubations in the pH7 buffer (**c**). TLC analysis showed that PA-like compounds were formed in the incubation of FP, epicatechin, and hydrogen peroxide ((FP+H_2_O_2_), but hardly detected in the incubation of FP and epicatechin (FP) and were not detected in two BSA controls (BSA+H_2_O_2_ and BSA) (**d**). **e,** TIC profiles showed that in the pH6 buffer, EC-P1, EC-P2 and other peaks were formed from the incubation consisting of FP, epicatechin, and hydrogen peroxide (FP+EC+H_2_O_2_) but not produced from three control incubations, FP and epicatechin (FP+EC), BSA, epicatechin and hydrogen peroxide (BSA+EC+H_2_O_2_), and BSA and epicatechin (BSA+EC). **f,** TIC profiles showed that in the pH8 buffer, EP-P1 was only detected from the incubation consisting of FP and epicatechin (FP+EC) but not formed in three other incubations, FP, epicatechin, and hydrogen peroxide (FP+EC+H_2_O_2_), BSA, epicatechin, and hydrogen peroxide (BSA+EC+H_2_O_2_), and BSA and epicatechin (BSA+EC).

**Fig. S9.**
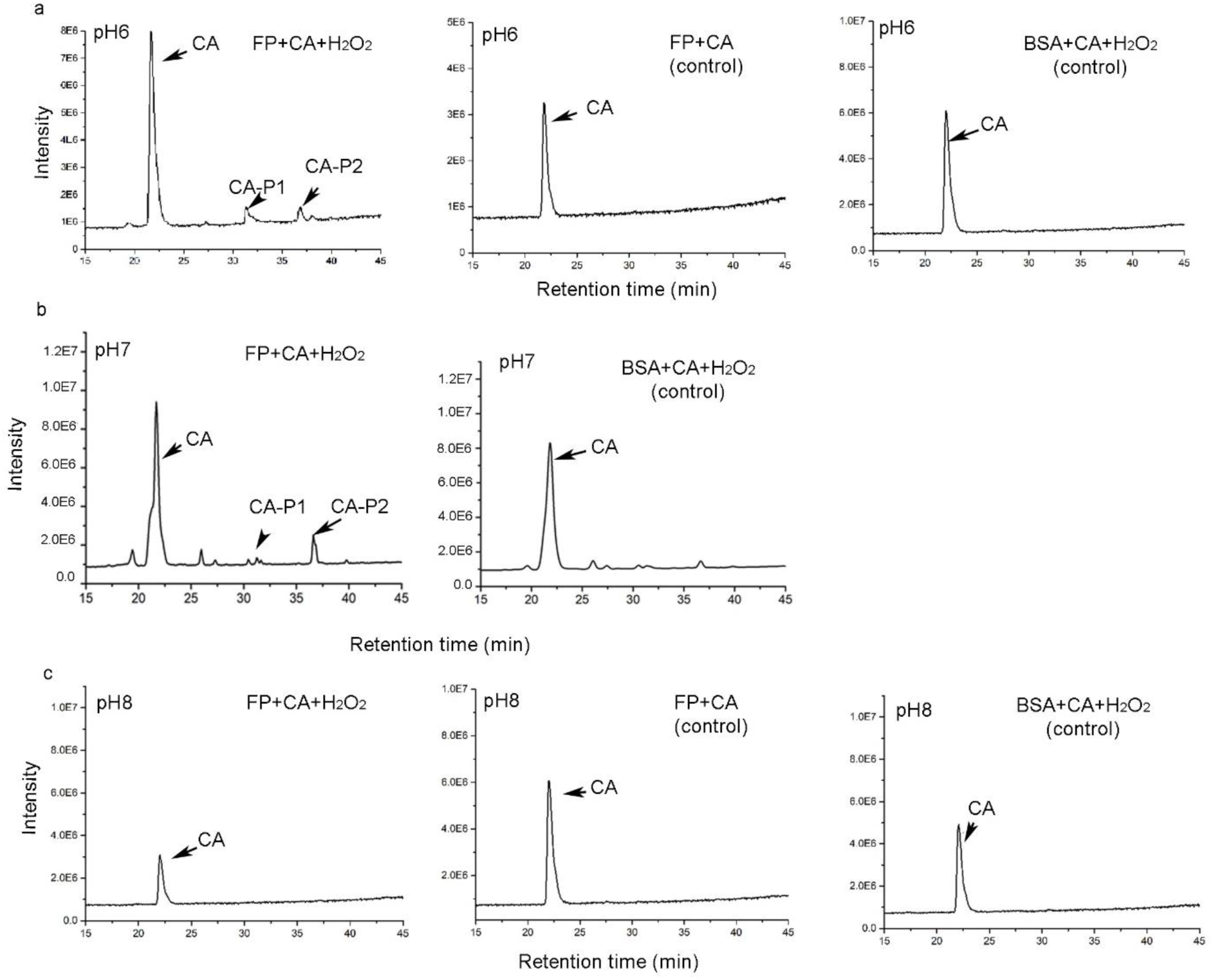
Effects of hydrogen peroxide and three pH buffers on the formation of yellowish products in the enzymatic reactions consisting of FP and catechin. **a,** TIC profiles showed that in the pH6 buffer, CA-P1 and CA-P2 were formed in the incubation consisting of FP, catechin, and hydrogen peroxide (left: FP+CA+H2O2) but not formed in the two control incubations, FP and catechin (FP+CA) and BSA, catechin, and hydrogen peroxide (BSA+CA+H_2_O_2_). **b,** TIC profiles showed that in the pH7 buffer, CA-P1 and CA-P2 were formed in the incubation consisting of FP, catechin, and hydrogen peroxide (FP+ CA+H_2_O_2_) but not formed in the control incubation consisting of BSA, catechin, and hydrogen peroxide (BSA+CA+H_2_O_2_). **c,** TIC profiles showed that in the pH8 buffer, no products were formed in three incubations FP, catechin, and hydrogen peroxide (FP+CA+H_2_O_2_), FP and catechin (FP+CA), and BSA, catechin, and hydrogen peroxide (BSA+CA+H_2_O_2_).

**Fig. S10.**
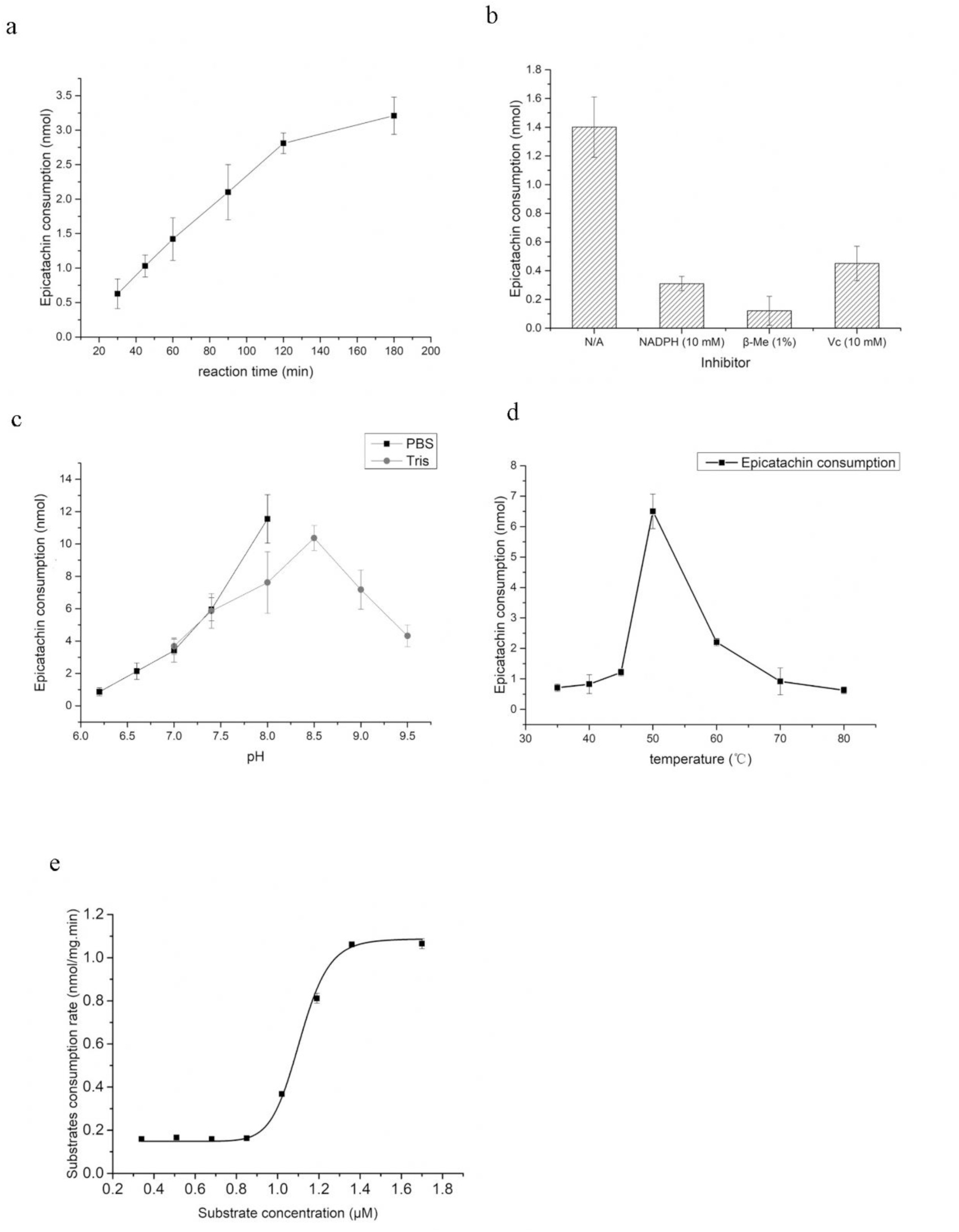
Effects of reaction time, inhibitors, pH values, temperatures, and epicatechin concentrations on enzyme activity. The recombinant FP was used to test these conditions. Due to at least two products formed in the reactions, the reduction of epicatechin concentrations was used to optimize reaction time, to identify inhibitors, to estimate pH and temperature optima, and to calculate Km, Kcat, and Vmax values. **a-d,** plots show effects of reaction time (a), inhibitors (b), pH values (c), and temperature (d) on the activity of recombinant enzyme. **e,** a plot of velocity versus epicatechin contraction shows a sigmoid kinetics of the recombinant FP.

**Fig. S11.**
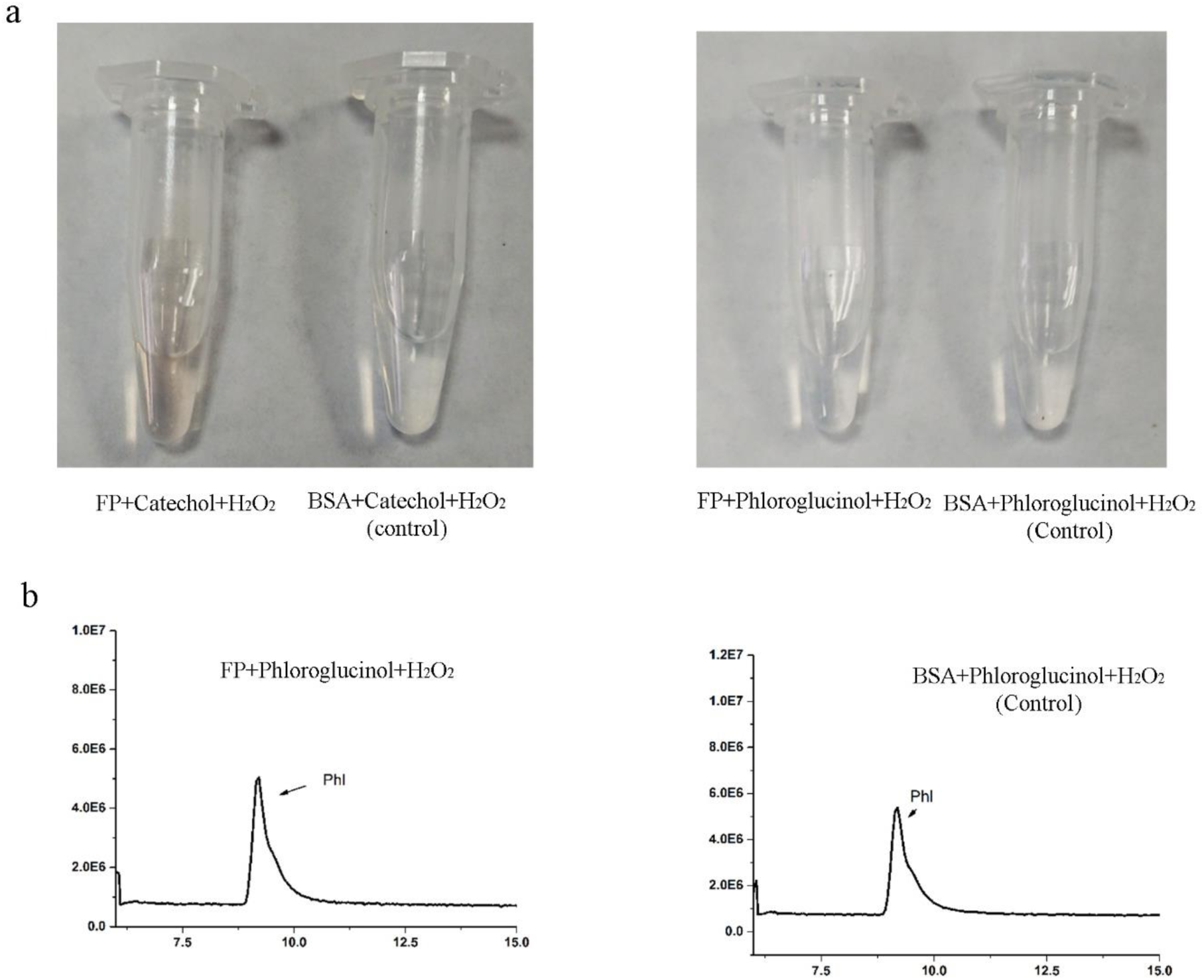
FP catalyzes catechol but does not catalyze phloroglucinol. Incubations were composed of catechol or phloroglucinol, FP or BSA control, and hydrogen peroxide in the pH6 buffer. **a,** images showed that the recombinant FP converted catechol to yellowish compounds but BSA control did not; the recombinant FP and BSA control did not have catalytic activity with phloroglucinol as the substrate. **b,** TIC profiles showed that the level of phloroglucinol was not changed between the FP and BSA control incubations, indicating that FP could not use this compound as a substrate. HPLC/MS could not detect substrate and products from the incubations of FP and catechol, because catechol was completely catalyzed into water-soluble quinone, which could not be extracted with ethyl acetate.

**Figure S12.**
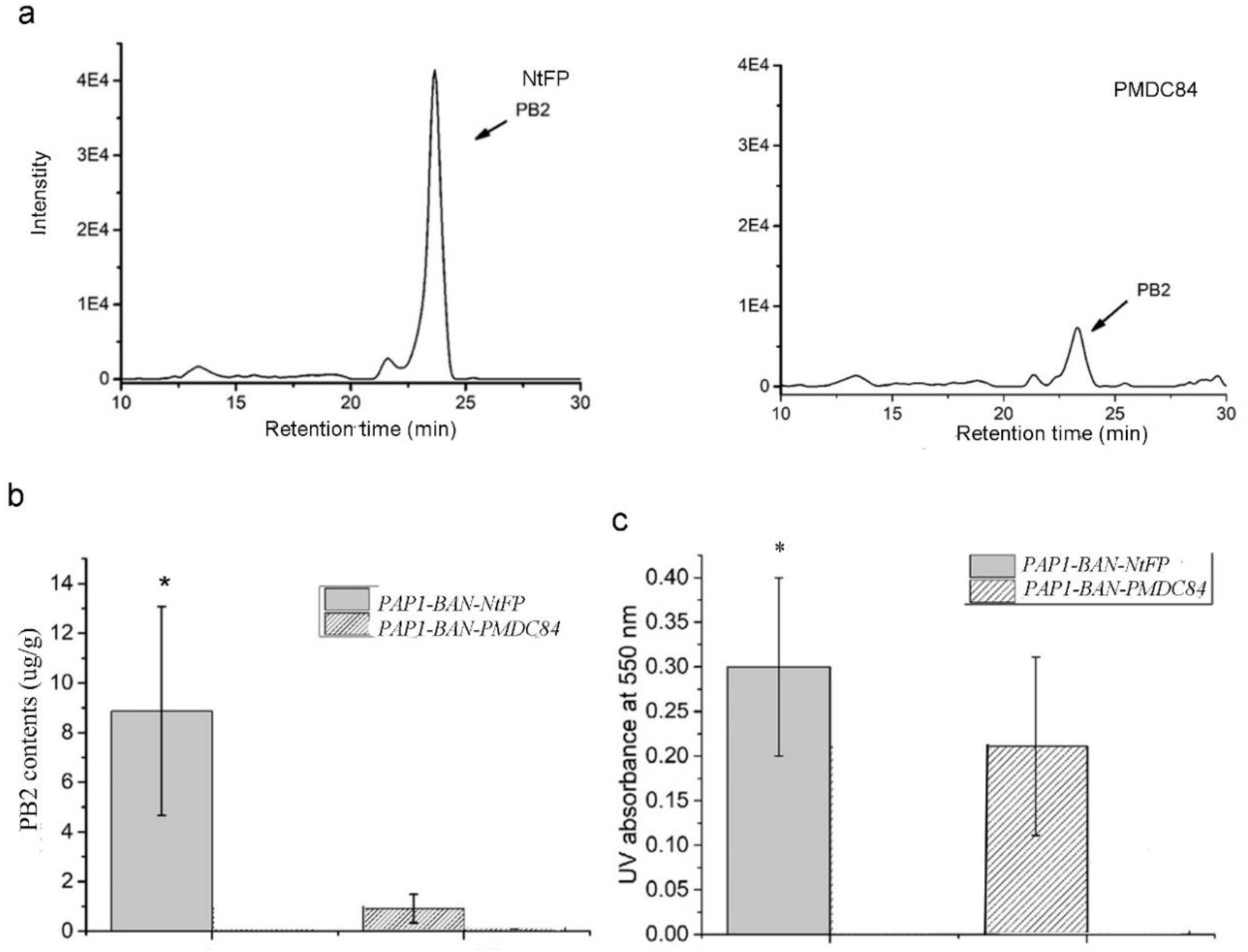
Analysis of proanthocyanidins in *FP (NtFP)* transgenic PAP1-BAN plants. **a,** EIC profiles show PB2 detected in leaves of both *PAP1-BAN-NtFP* and *PAP1-BAN-PMDC84* (vector control) transgenic tobacco plants. **b,** the PB2 content was significantly higher in leaves of *PAP1-BAN-NtFP* than in those of *PAP1-BAN-PMDC84* transgenic tobacco plants. **c,** the UV absorbent values of extracts boiled in butanol-HCl were measured at 550 nm to estimate the levels of PAs. The absorbent values were higher in leaf extracts of *PAP1-BAN-NtFP* plants than in those of *PAP1-BAN-PMDC84* transgenic tobacco plants, indicating the increase of PAs in *PAP1-BAN-NtFP* plants.

**Figs. S13-S30 support novel compounds biosynthesized by FP (Figures 4 and 5)**

**Fig. S13.**
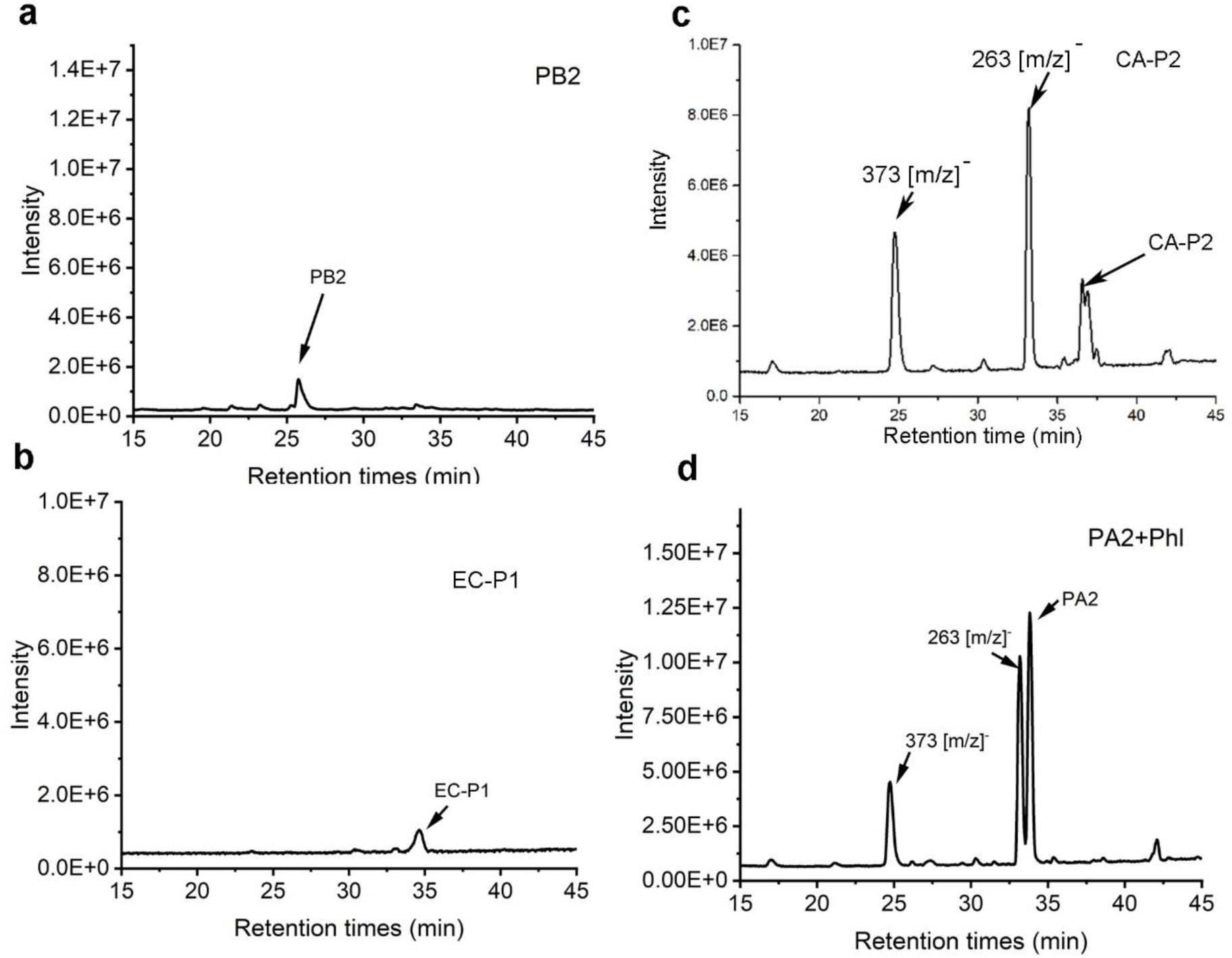
No catechin-phloroglucinol and epicatechin-phloroglucinol produced from hydrochloric acid (HCl)-catalyzed cleavage (phloroglucinolysis) of CA-P2 and procyanidin A2 (PA2) in presence of phloroglucinol (support Figure 4 e-g) a-b, TIC profiling showed procyanidin B2 standard and purified EC-P1 (both dissolved in methanol) prior to phloroglucinolysis as controls. c-d, TIC profiling did not detect catechin-phloroglucinol and epicatechin-phloroglucinol formed from the methanol HCl-catalyzed cleavage of CA-P2 (c) and PA2 (d). These data indicate that neither can HCl-methanol cleave CA-P1 to catechin and catechin carbocation, nor it can cleave PA2 to epicatechin and epicatechin carbocation. Therefore, phloroglucinol cannot nucleophilically attack epicatechin carbocation and catechin carbocation to form epicatechin-phloroglucinol and catechin-phloroglucinol, respectively. Peaks of 263 [m/z]^-^ and 373 [m/z]^-^ observed from the phloroglucinolysis were not observed in procyanidin B2 standard and EC-P1 because these two resulted from buffers used for phloroglucinolysis of PB2, EC-P1, CA-P2, and PA2.

**Fig. S14.**
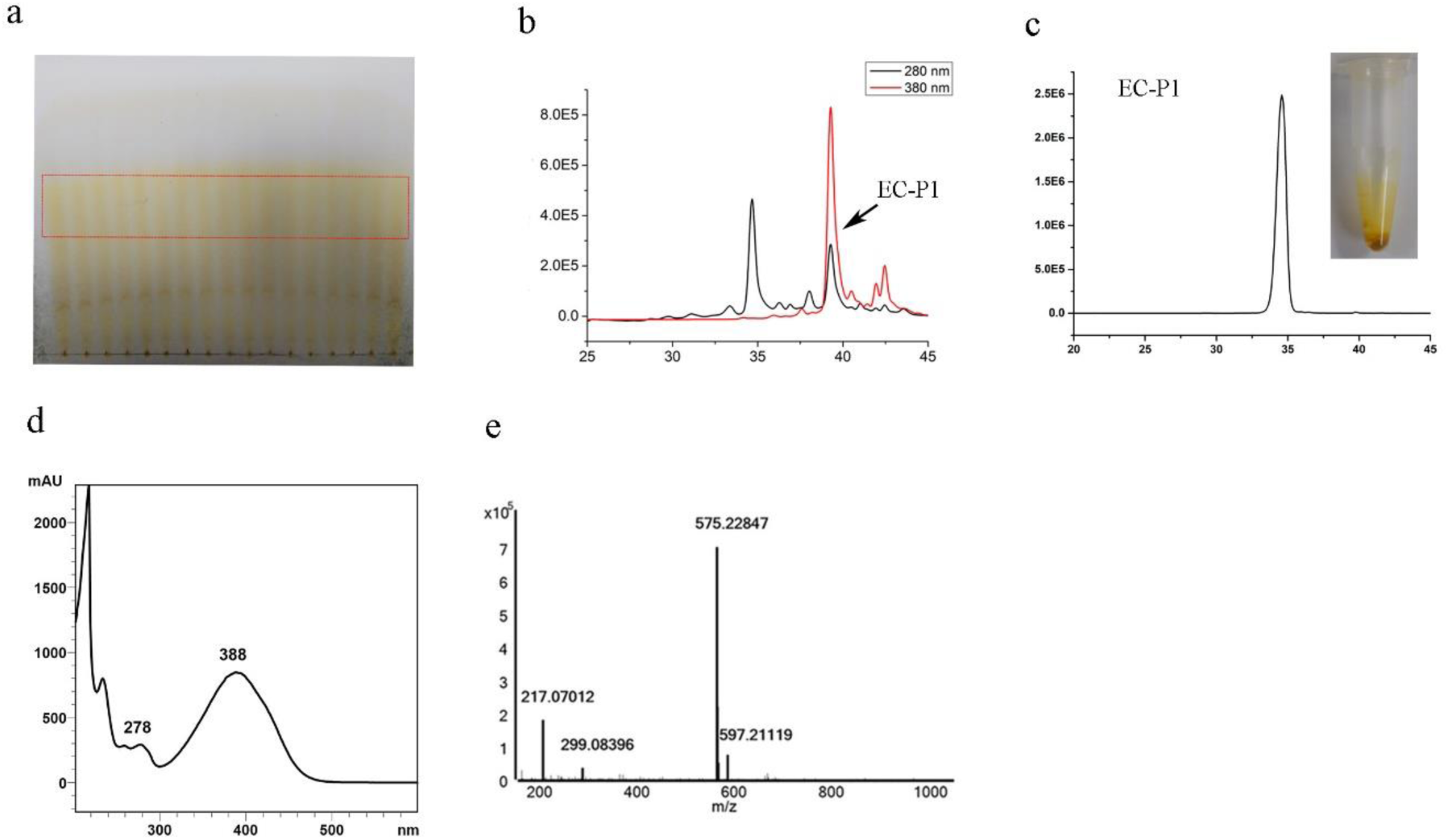
Purification of EC-P1 produced from enzymatic reactions and its UV spectrum and mass spectrum features. **a-c,** TLC preparation (**a**) and HPLC separation (**b**) were used to isolate EP-P1. Total ion chromatography (TIC) examination shows one single peak of EC-P1 purified (**c**). **d,** The UV-spectrum profile shows a featured absorption of EP-P1at 388 nm. **e,** The mass spectrum shows an accurate mass-to-charge ratio of EP-P1, 575.22847 [m/z]^-^.

**Figures S15-30, Data of NMR support Figure 5.**

**Fig. S15.**
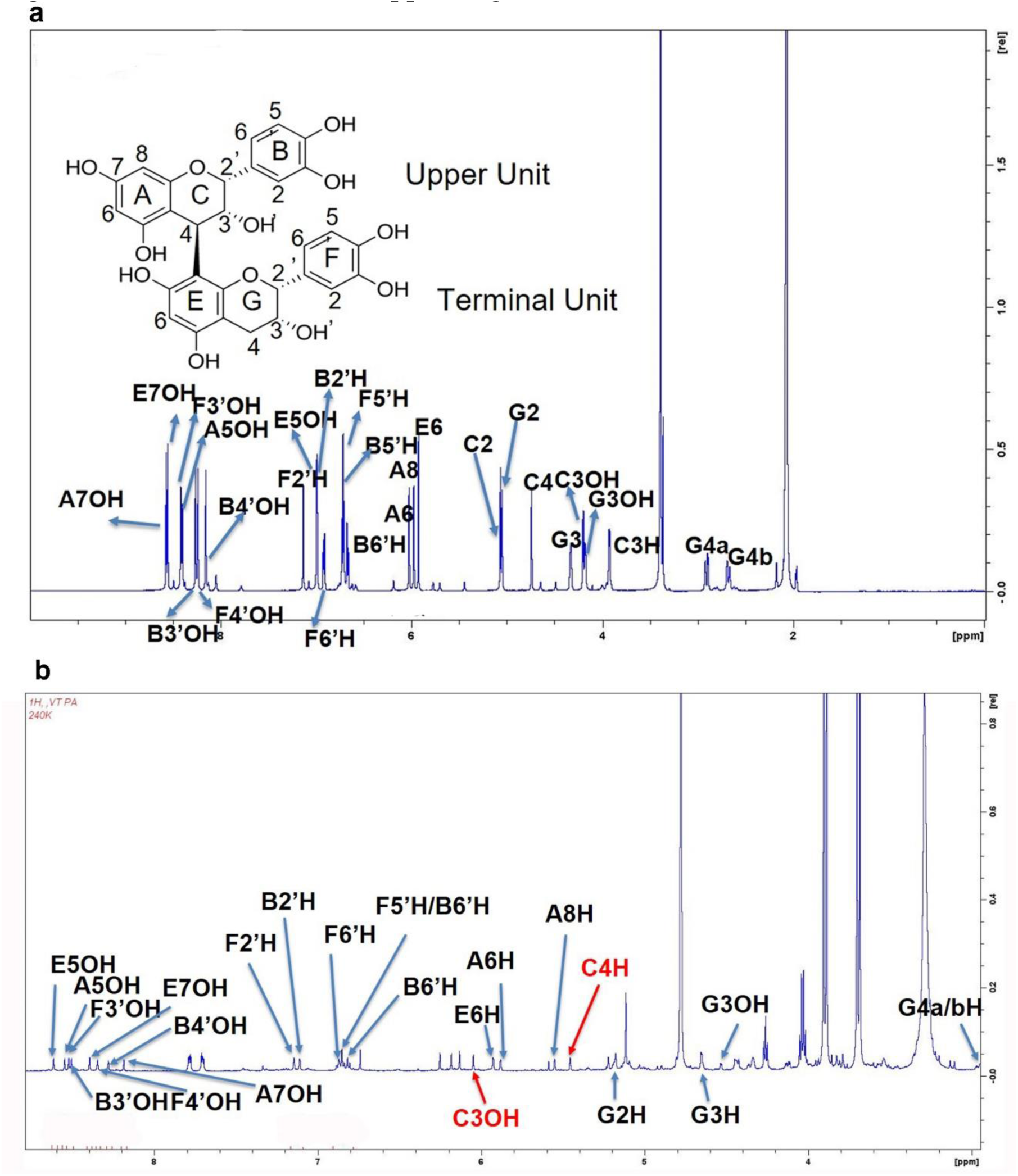
^1^H NMR profiles showing different PPM values of proton chemical shifts of PB2 and EC-P1. a, based on their PPM values, 16 protons were assigned to the A, B, C, E, F, and G rings and 10 protons were assigned to the10 hydroxyl groups. b, 13 protons (black bold letters) were assigned to A, B, C, E, F, and G rings and 9 protons were assigned to the 9 hydroxyl groups A, B, C, E, F, and G rings. Red colored “C3OH” and “C4H” mean that these two protons predicted by ^1^H NMR test were assigned by ^1^H NMR chemical shift and ROSEY NMR analysis.

**Fig. S16.**
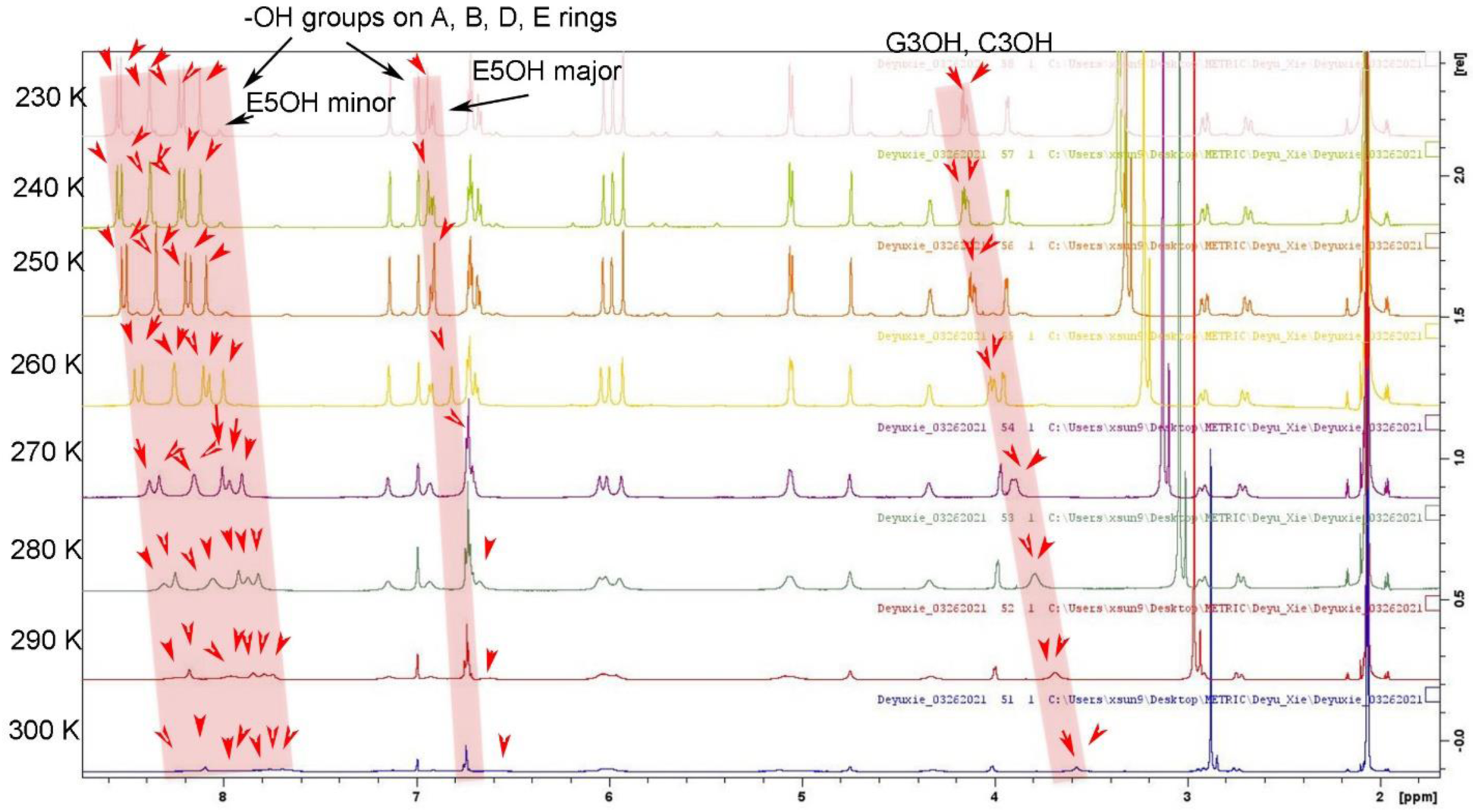
Effects of seven different temperatures on ^1^H NMR profiles of 10 hydroxyl groups of PB2 and assignment of their protons. ^1^H NMR profiles show that seven temperatures affect 10 proton chemical shifts in 10 hydroxyl groups. The PPM value of each proton chemical shift increases as the tested temperature decreases from 300K to 240K. The red arrowheads indicate each proton in the hydroxyl groups, and tilted pink rectangles indicate the PPM shifting from low to high values as the tested temperature decreases.

**Fig. S17.**
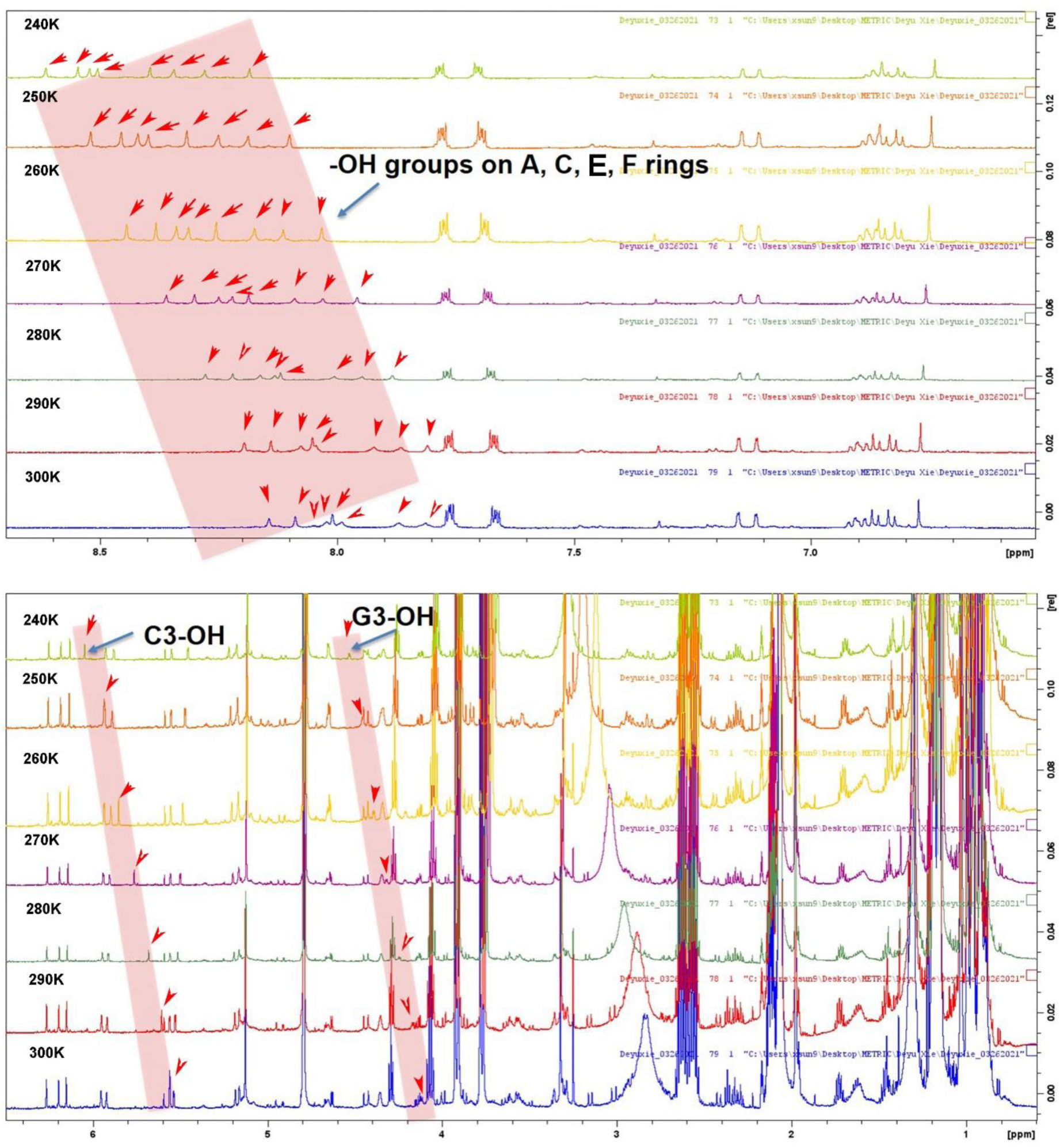
Effects of seven different temperatures on ^1^H NMR profiles of 10 hydroxyl groups of EC-P1 and assignment of their protons. ^1^H NMR profiles show effects of temperatures on 8 proton chemical shifts in eight hydroxyl groups on A, C, E, and E rings (the upper panel) and 2 proton chemical shifts of C3-OH on C ring and G3-OH on G-ring (the bottom panel). The PPM value of each proton chemical shift increases as the tested temperature decreases from 300K to 240K. The red arrowheads indicate each proton in the hydroxyl groups, and tilted pink rectangles indicate the PPM shifting from low to high values as the tested temperature decreases.

**Fig. S18.**
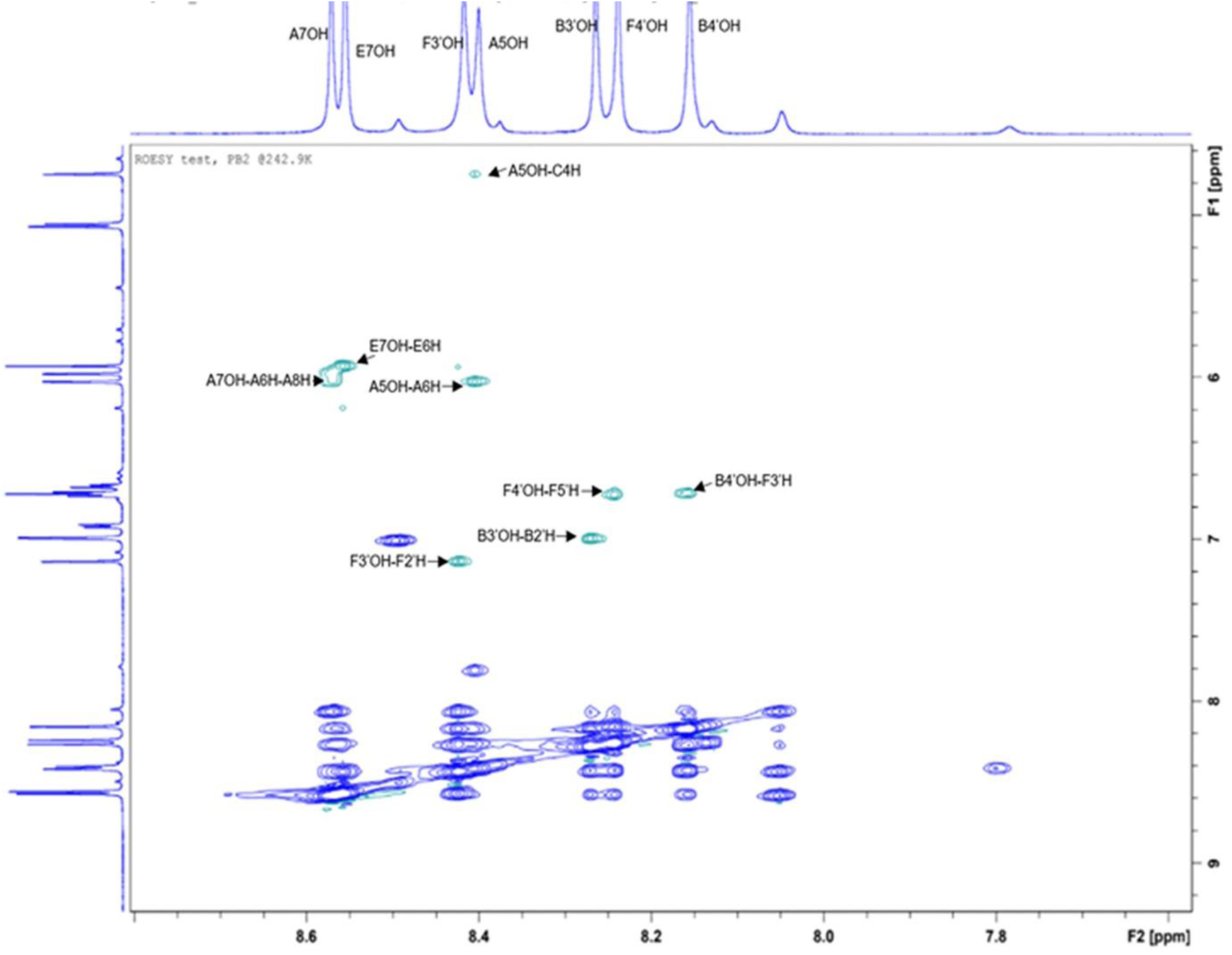
2D ROESY NMR analysis of procyanidin B2. This 2D NMR plot shows 8 cross peaks formed by the protons of A7OH, E7OH, F3’OH, A5OH, B3’OH, F4’OH, and B4’OH with PPM values from 8.0 to 8.8 (F2 axis) and their nearby protons with PPM values from 4.8 to 7.4 (F1 axis). These cross peaks establish the correlations of A7OH and A6H, A7OH and A8H, E7OH and E6H, F3’OH and F2’H, A5OH and A6H, B3’OH and B2’H, F4’OH and F5’H, and B4’OH and F3’H.

**Fig. S19.**
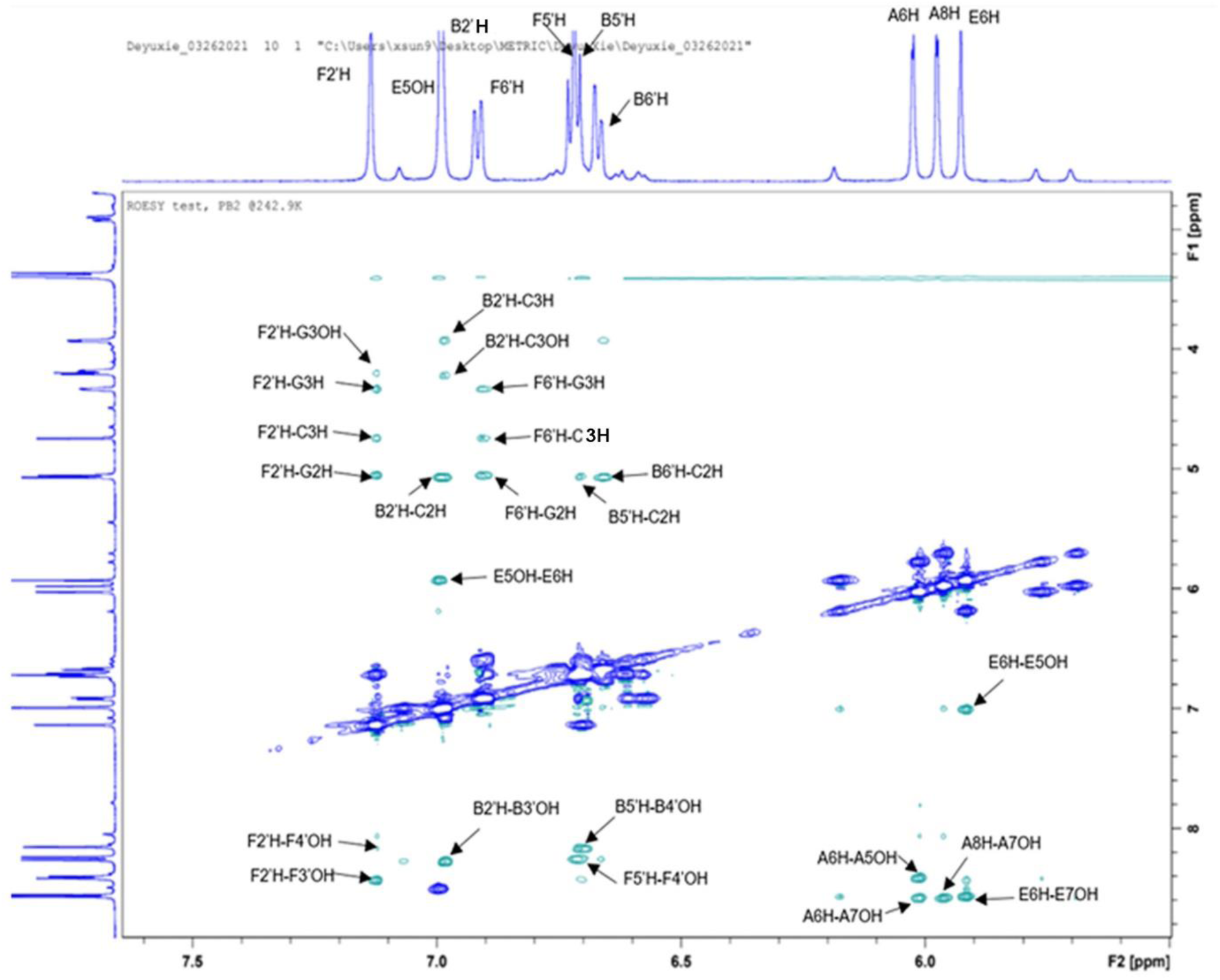
2D ROESY NMR analysis of procyanidin B2. This 2D NMR plot shows 23 cross peaks formed by the protons of F2’H, E5OH, B2’H, F6’H, F5’H, B5’H, B6’H, A6H, A8H, and E6H with PPM values from 5.5 to 7.5 (F2 axis) and their nearby protons with PPM values from 3.4 to 8.8 (F1 axis). These cross peaks establish correlations between these protons.

**Fig. S20.**
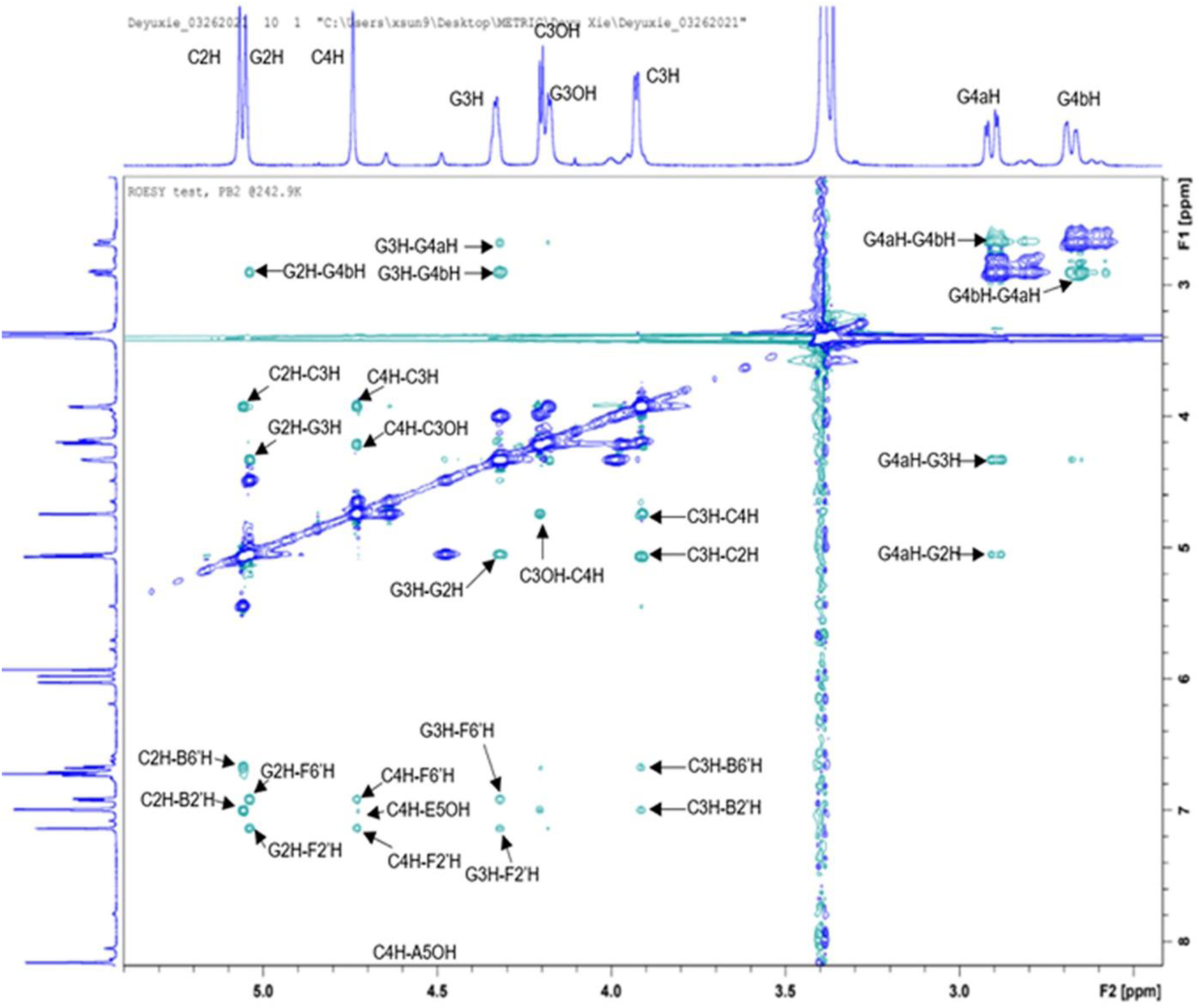
2D ROESY NMR analysis of procyanidin B2. This 2D NMR plot shows 27 cross peaks formed by the protons of C2H, G2H, C4H, G3H, C3OH, G3OH, C3H, G4aH, and G4bH with PPM values from 2.5 to 5.4 (F2 axis) and their nearby protons with PPM values from 2.5 to 8.2 (F1 axis). These cross peaks establish the correlations between these protons.

**Fig. S21.**
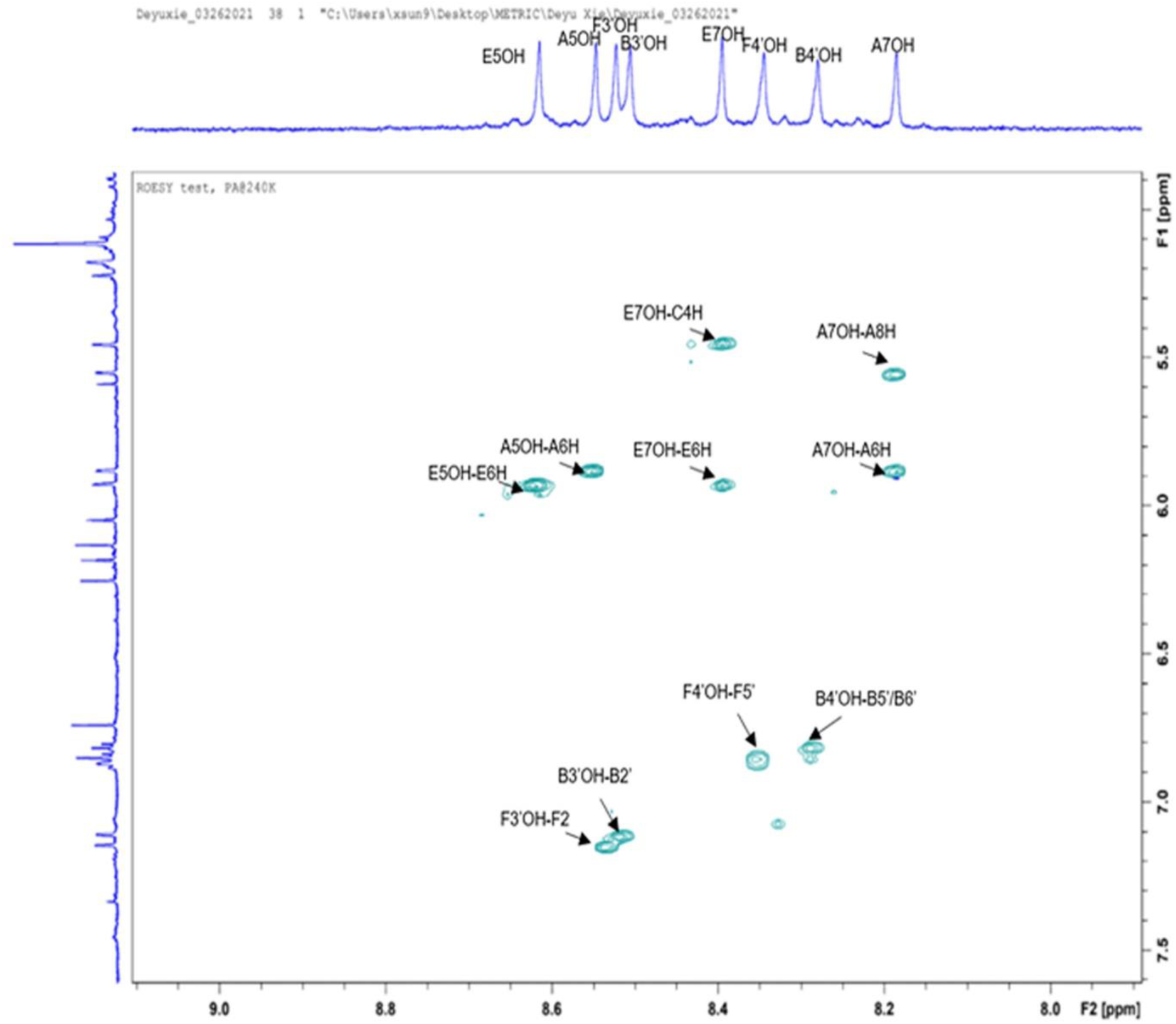
2D ROESY NMR analysis of EC-P1. This 2D NMR plot shows 10 cross peaks formed by the protons of E5OH, A5OH, F3’OH, B3’OH, E7OH, F4’OH, B4’OH, and A7OH with PPM values from 8.0 to 8.8 (F2 axis) and their nearby protons with PPM values from 5.0 to 7.5 (F1 axis). These cross peaks establish the correlations between these protons.

**Fig. S22.**
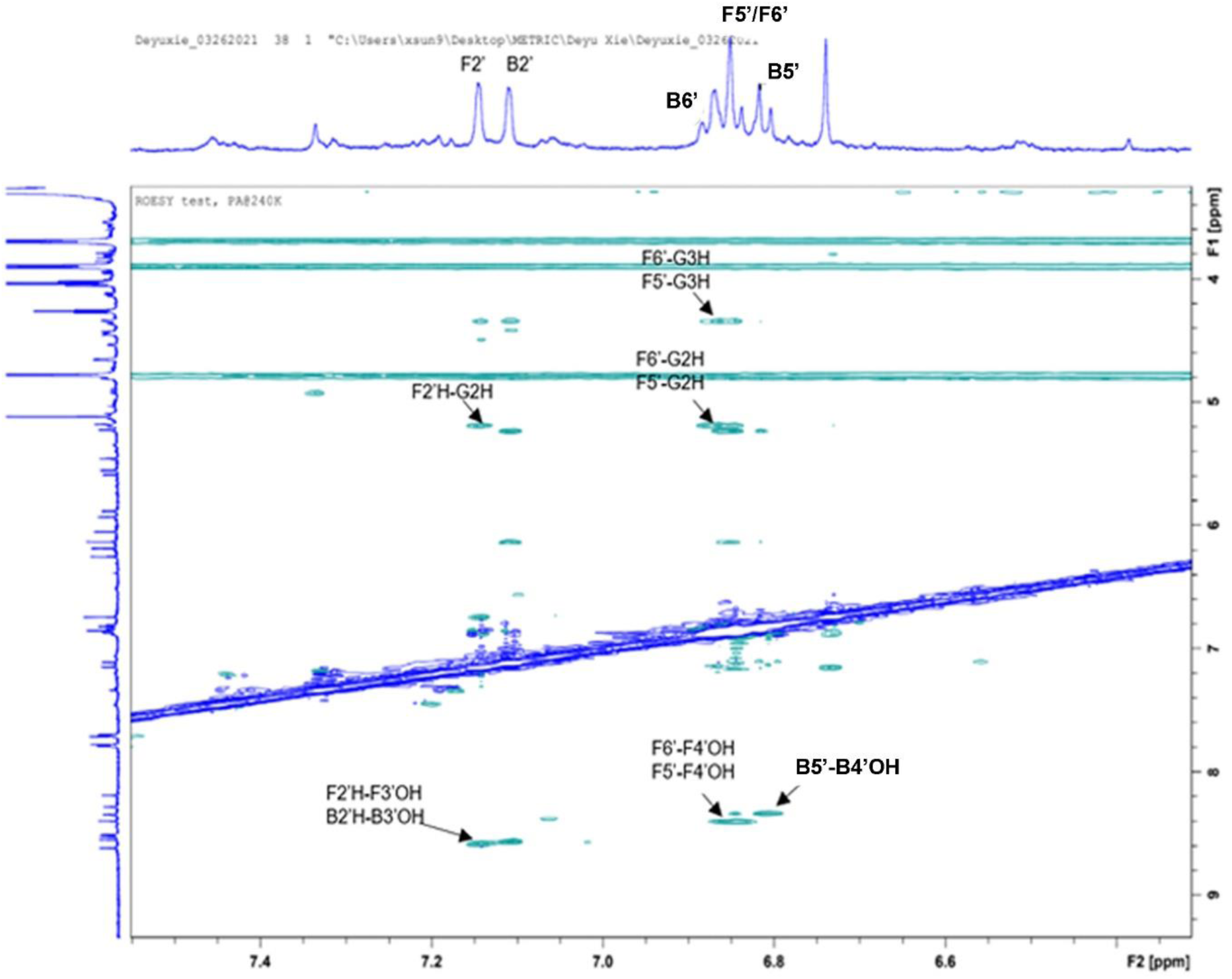
2D ROESY NMR analysis of EC-P1. This 2D NMR plot shows 10 cross peaks formed by the protons of F2’H, B2’H, B6’H, F5’H, F6’H, and B5’H with PPM values from 6.6-7.2 (F2 axis) and their nearby protons with PPM values from 4.0-8.8 (F1 axis). These cross peaks establish the correlations between these protons. However, no nearby protons crossed with the proton of B6’H were identified.

**Fig. S23.**
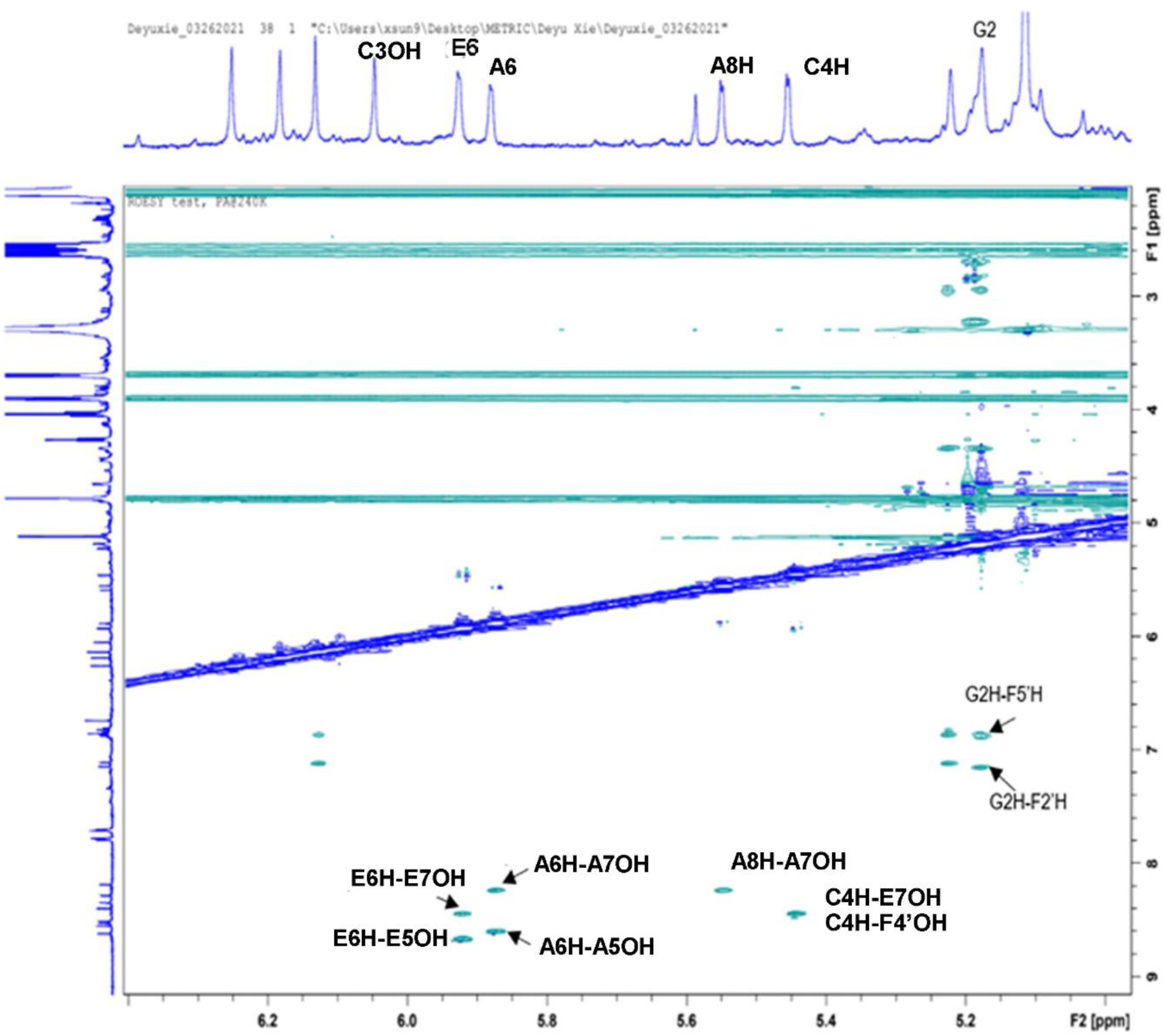
2D ROESY NMR analysis of EC-P1. This 2D NMR plot shows 9 cross peaks formed by the protons of E6H, A6H, A8H, C4H, and G2H with PPM values from 5.0 to 6.0 and their nearby protons with PPM values from 6.0 to 8.8. These cross peaks establish the correlations between these protons. However, no nearby protons crossed with the proton of C3OH were identified.

**Fig. S24.**
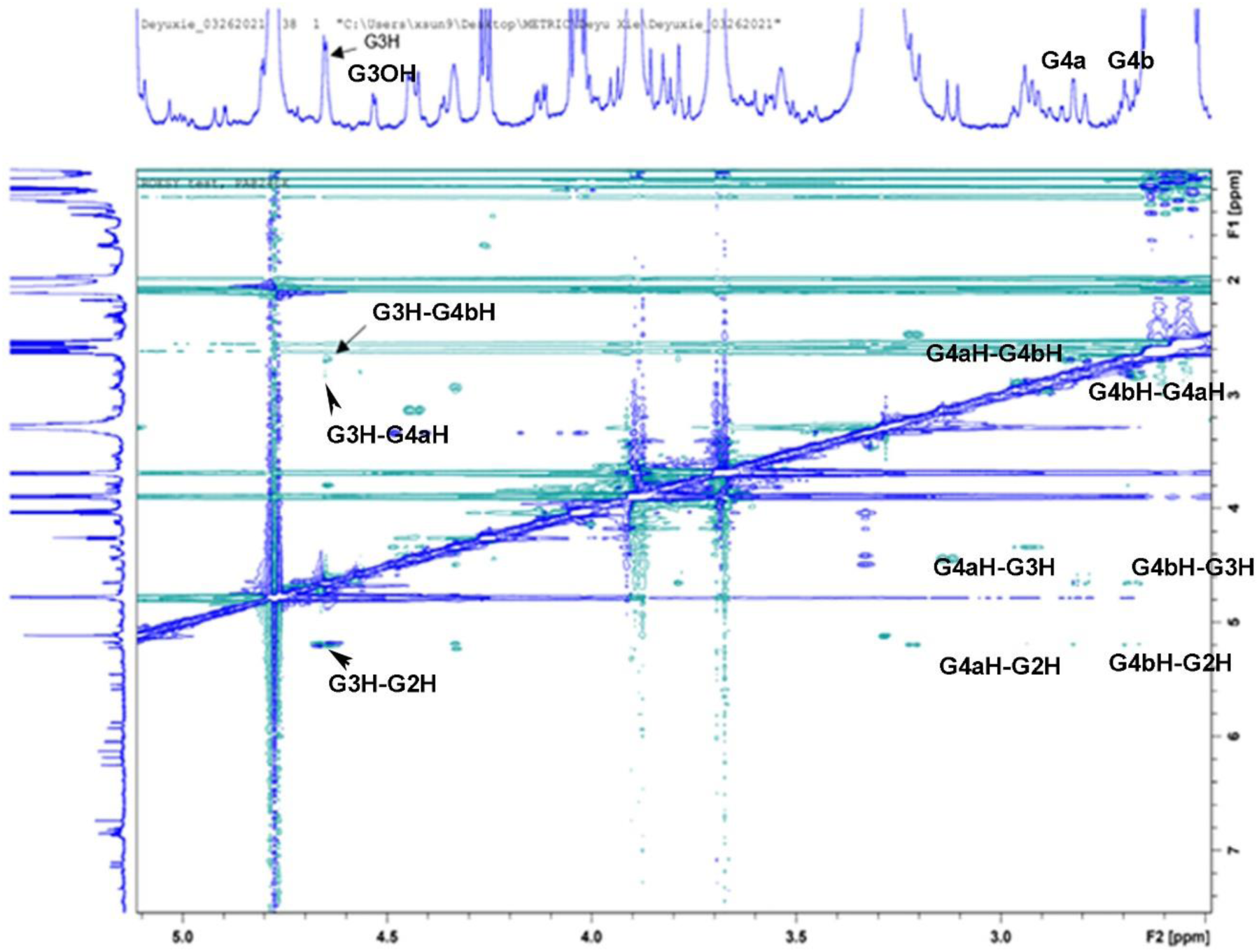
2D ROESY NMR analysis of EC-P1. This 2D NMR plot shows 9 cross peaks formed by the protons of G3H, G3OH, G4aH, and G4bH with PPM values from 2.5 to 5.0 (F2 axis) and their nearby protons with PPM values from 2.4 to 5.6 (F1 axis). These cross peaks establish the correlations between these protons. However, no nearby protons crossed with the protons of G3OH and G4aH were identified.

**Fig. S25.**
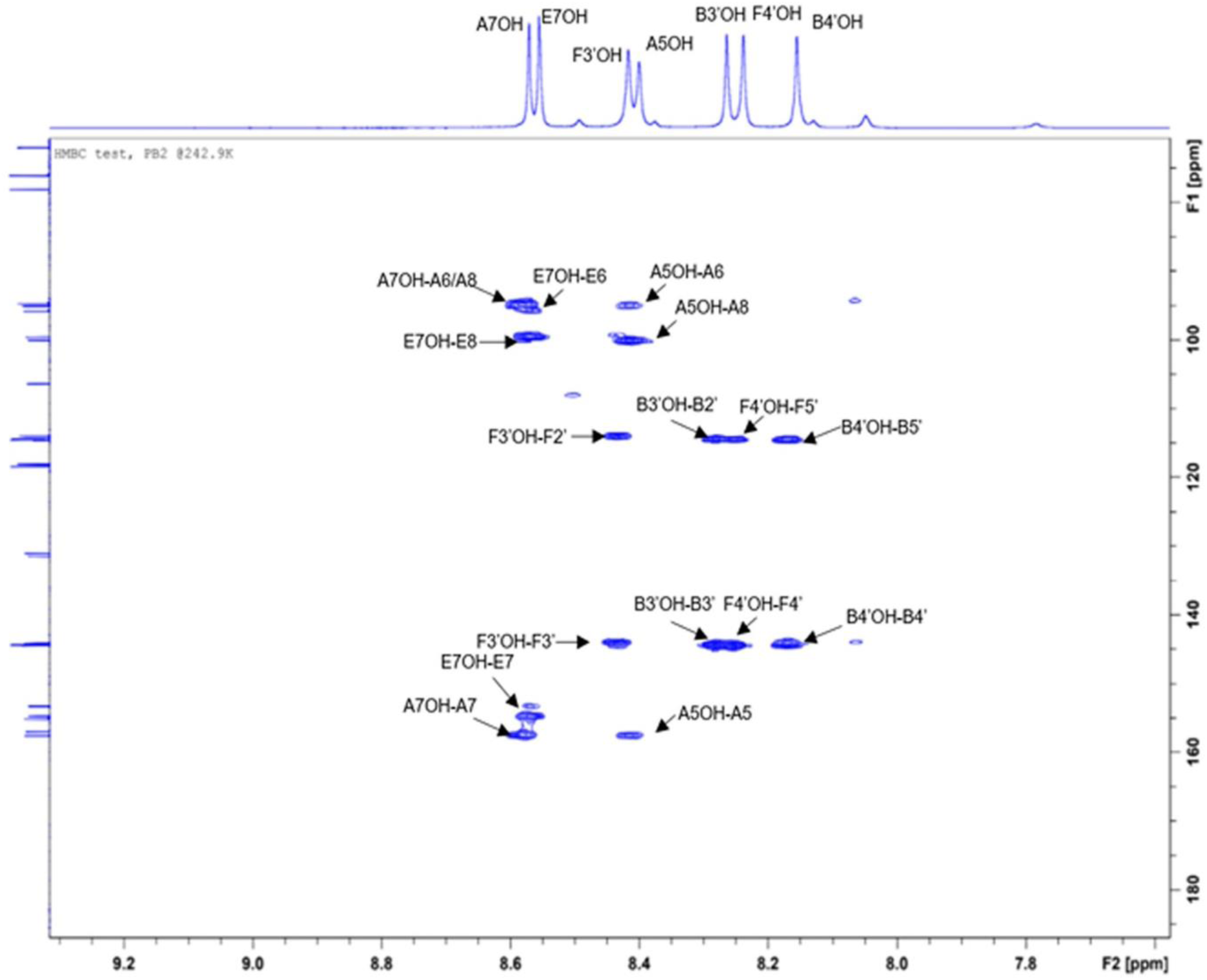
2D HMBC NMR analysis of PB2. This 2D NMR plot shows 16 cross peaks formed by the protons of A7OH, E7OH, F3’OH, A5OH, B3’OH, F4’OH, and B4’OH with PPM values from 8.0-8.8 (F2 axis) and their nearby carbons with PPM values from 80 to 165 (F1 axis). These cross peaks establish the correlations between these protons and carbons.

**Fig. S26.**
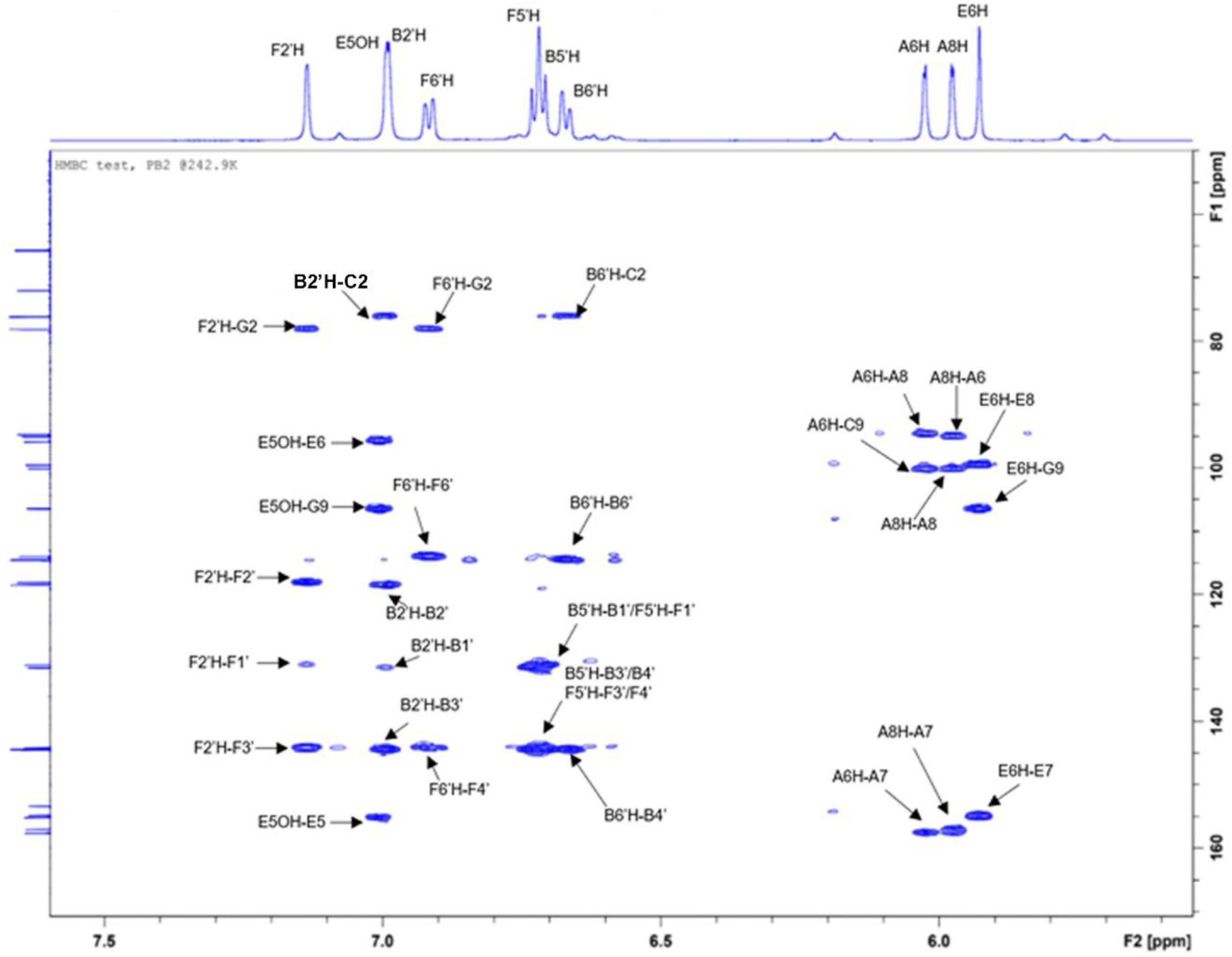
The 2D HMBC NMR analysis of PB2. This 2D NMR plot shows 32 cross peaks formed by the protons of F2’H, E5OH, B2’H, F6’H, F5’H, B5’H, B6’H, A6H, A8H, and E6H with PPM values from 5.4 to 7.5 (F2 axis) and their nearby carbons with PPM values from 90 to 160 (F1 axis). These cross peaks establish the correlations between these protons and carbons.

**Fig. S27.**
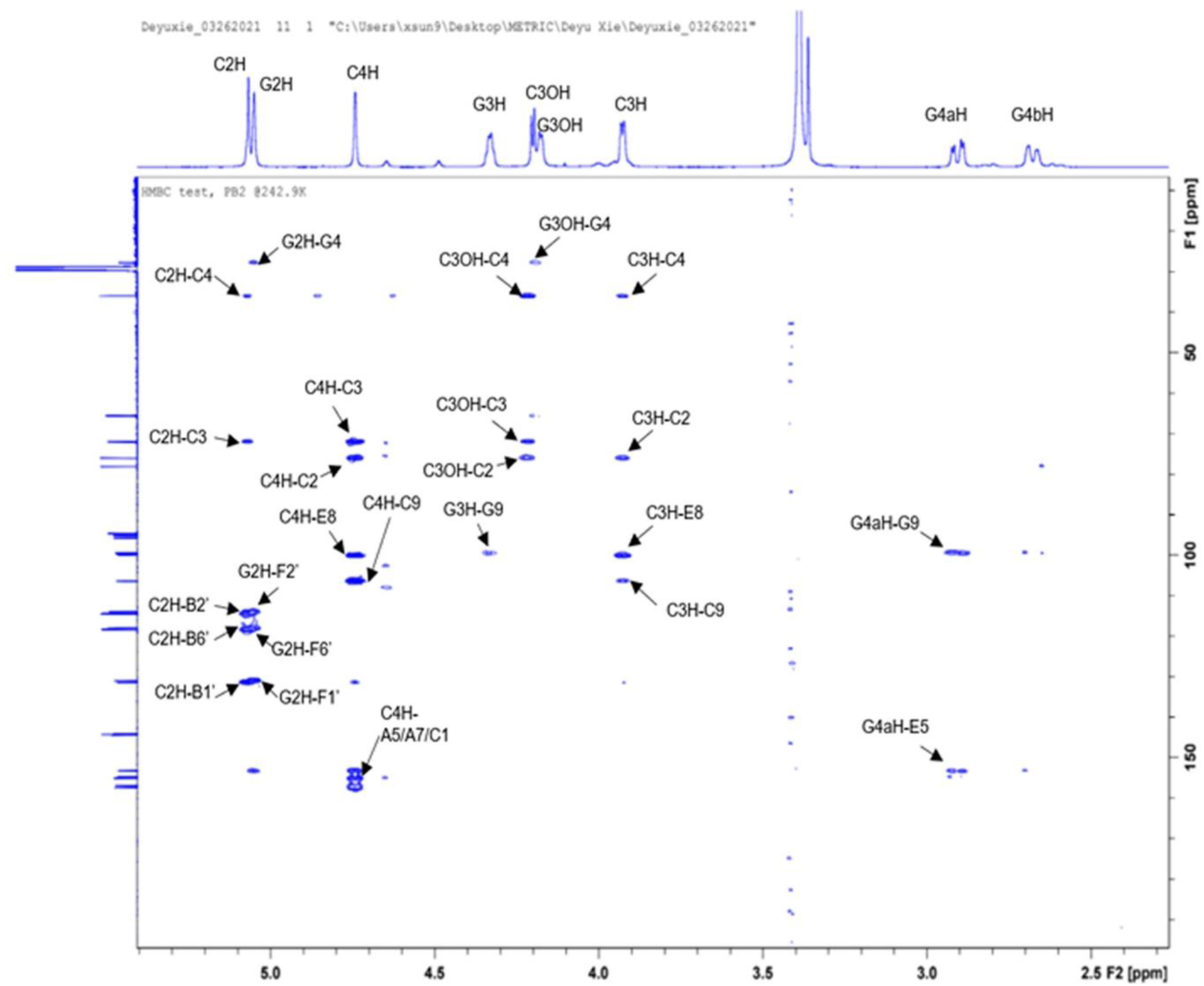
2D HMBC NMR analysis of PB2. This 2D NMR plot shows 27 cross peaks formed by the protons of C2H, G2H, C4H, G3H, C3OH, G3OH, C3H, G4aH, and G4bH with PPM values from 2.5 to 5.4 (F2 axis) and their nearby carbons with PPM values from 20 to 160 (F1 axis). These cross peaks establish the correlations between these protons and carbons. However, no nearby carbons crossed with G4bH were identified.

**Fig. S28.**
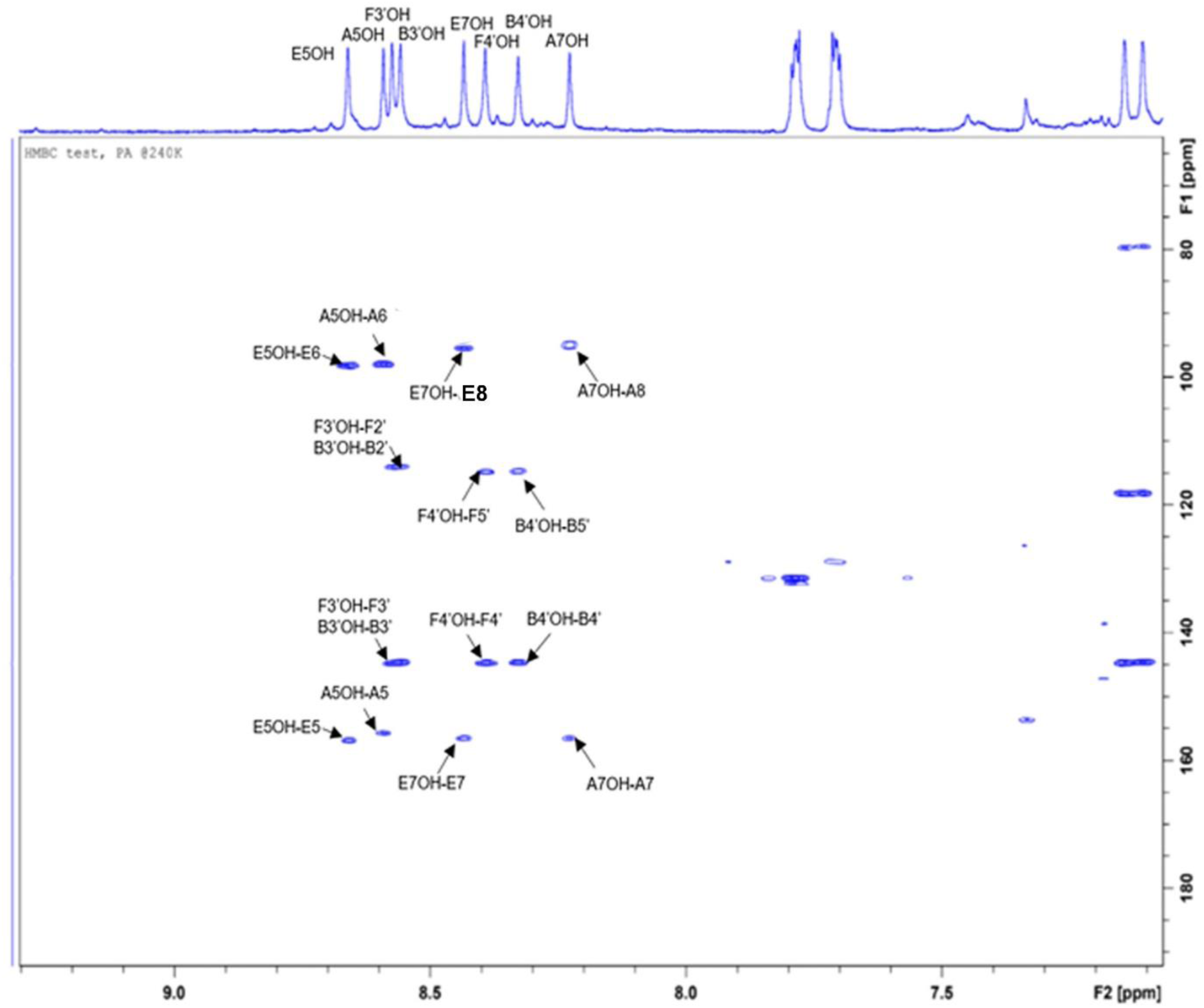
2D HMBC NMR analysis of EC-P1. This 2D NMR plot shows 16 cross peaks formed by the protons of E5OH, A5OH, F3’OH, B3’OH, E7OH, F4’OH, B4’OH, and A7OH with PPM values from 8.0 to 8.8 (F2 axis) and their nearby carbons with PPM values from 80 to 160 (F1 axis). These cross peaks establish the correlations between these protons and carbons.

**Fig. S29.**
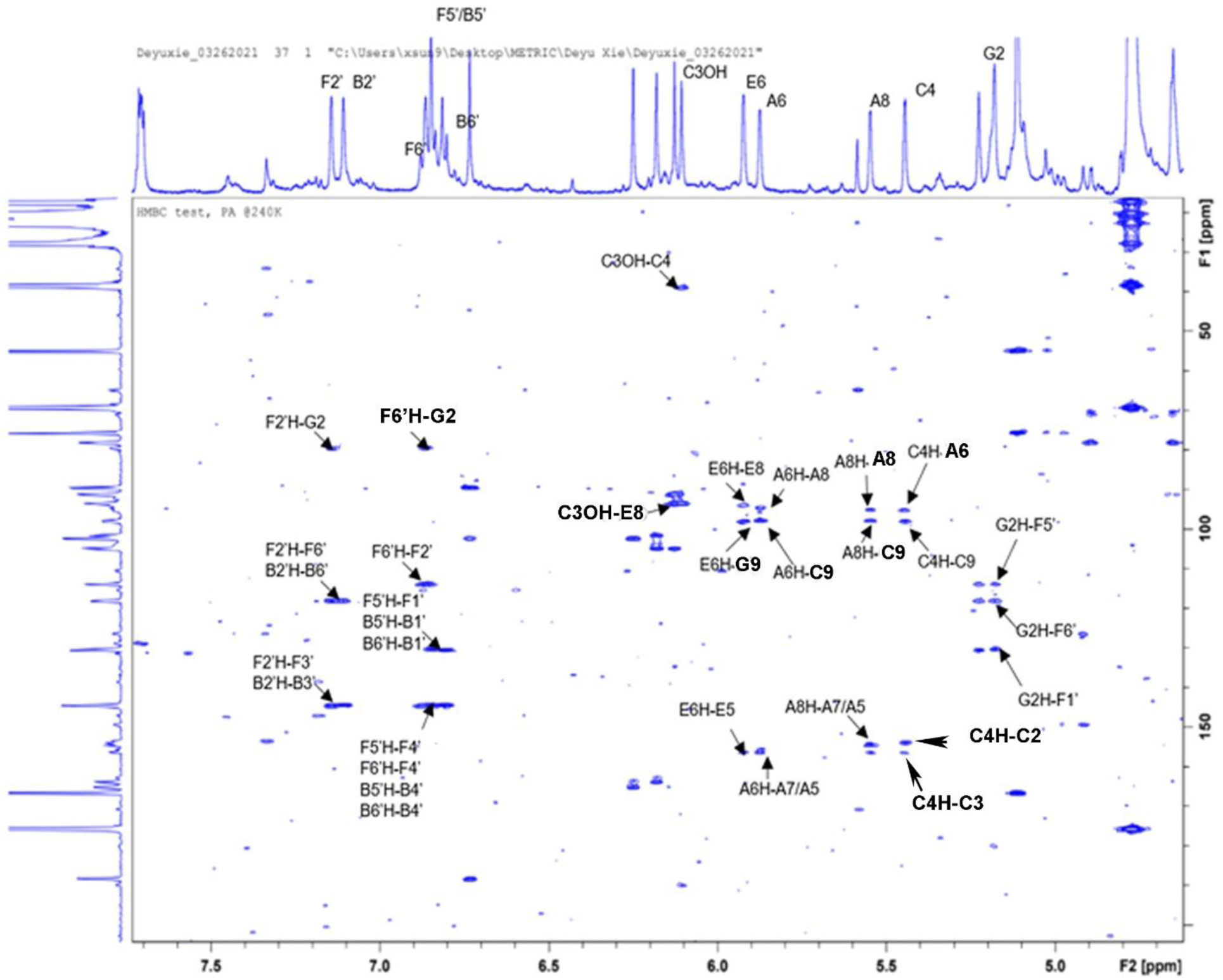
2D HMBC NMR analysis of EC-P1. This 2D NMR plot shows 34 cross peaks formed by the protons of F2’H, B2’H, F6’H, F5’H, B5’H, B6’H, C3OH, E6H, A6H, A8H, C4H, and G2H with PPM values from 5.0 to 7.5 (F2 axis) and their nearby carbons with PPM values from 30 to 160 (F1 axis). These cross peaks establish the correlations between these protons and carbons.

**Fig. S30.**
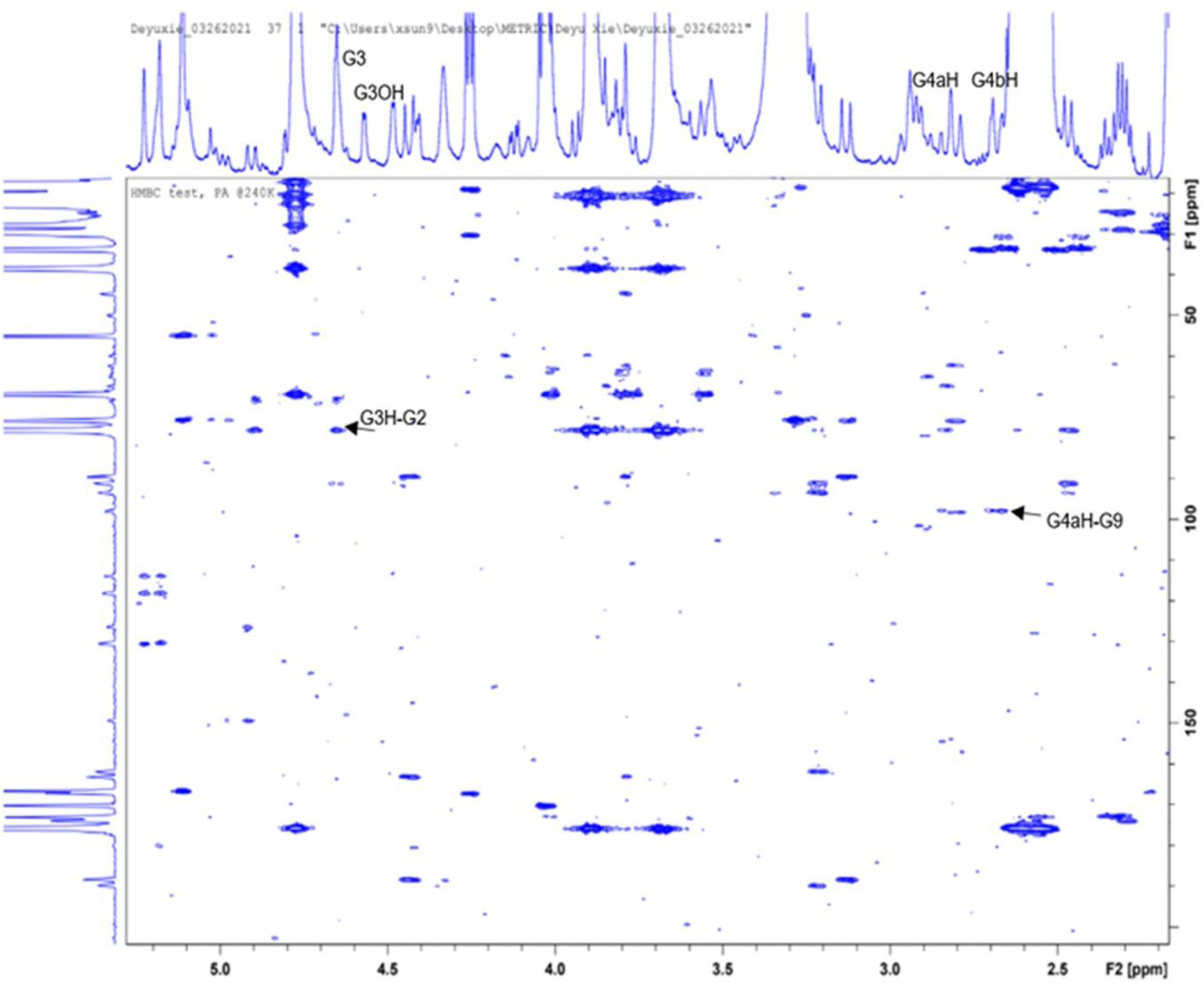
2D HMBC NMR analysis of EC-P1. This 2D NMR plot shows two cross peaks formed by the protons of G3H, G3OH, G4aH, and G4bH with PPM values from 2.5 to 5.0 (F2 axis) and their nearby carbons with PPM values from 20 to 160. These cross peaks establish the correlations between these protons and carbons. However, no nearby carbons crossed with G3OH and G4bH were identified.

**Figs S31—S33 provide evidence of carbocation and oligomer formation (Figure 5)**

**Fig. S31.**
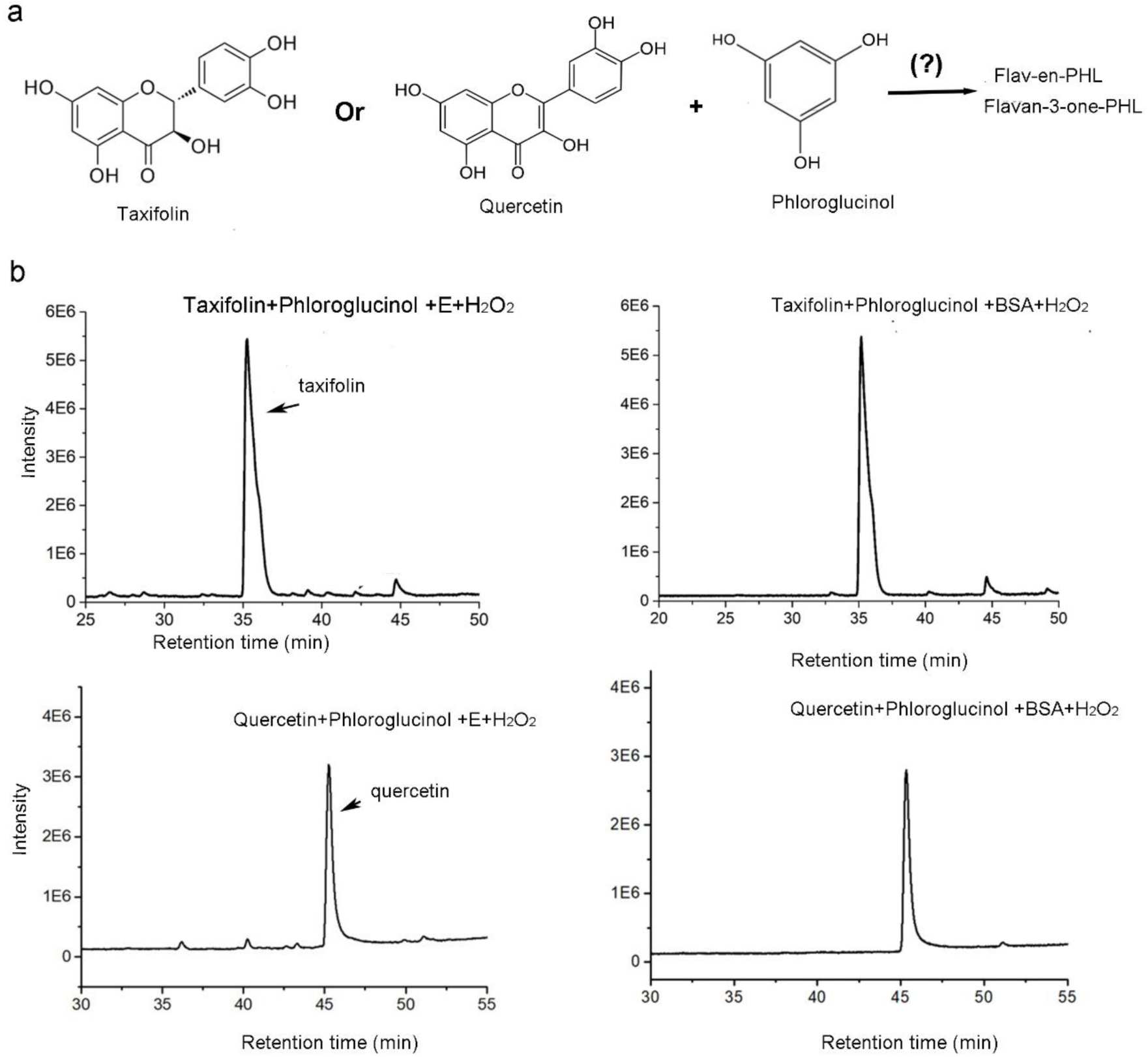
No 412 Dalton compound formed from incubations of quercetin or taxifolin, phloroglucinol, and recombinant FP. **a,** a scheme presents a hypothesis whether the incubation with enzyme, quercetin or taxifolin, phloroglucinol, and hydrogen peroxide produces conjugates in pH6 buffer. **b,** TIC profiles showed that no compound with a 411 [m/z]^-^ (molecular weight 412 Dalton) was detected from the incubations of FP (E: enzyme) or BSA (control), taxifolin or quercetin, phloroglucinol, and hydrogen peroxide in pH6 buffer.

**Fig. S32.**
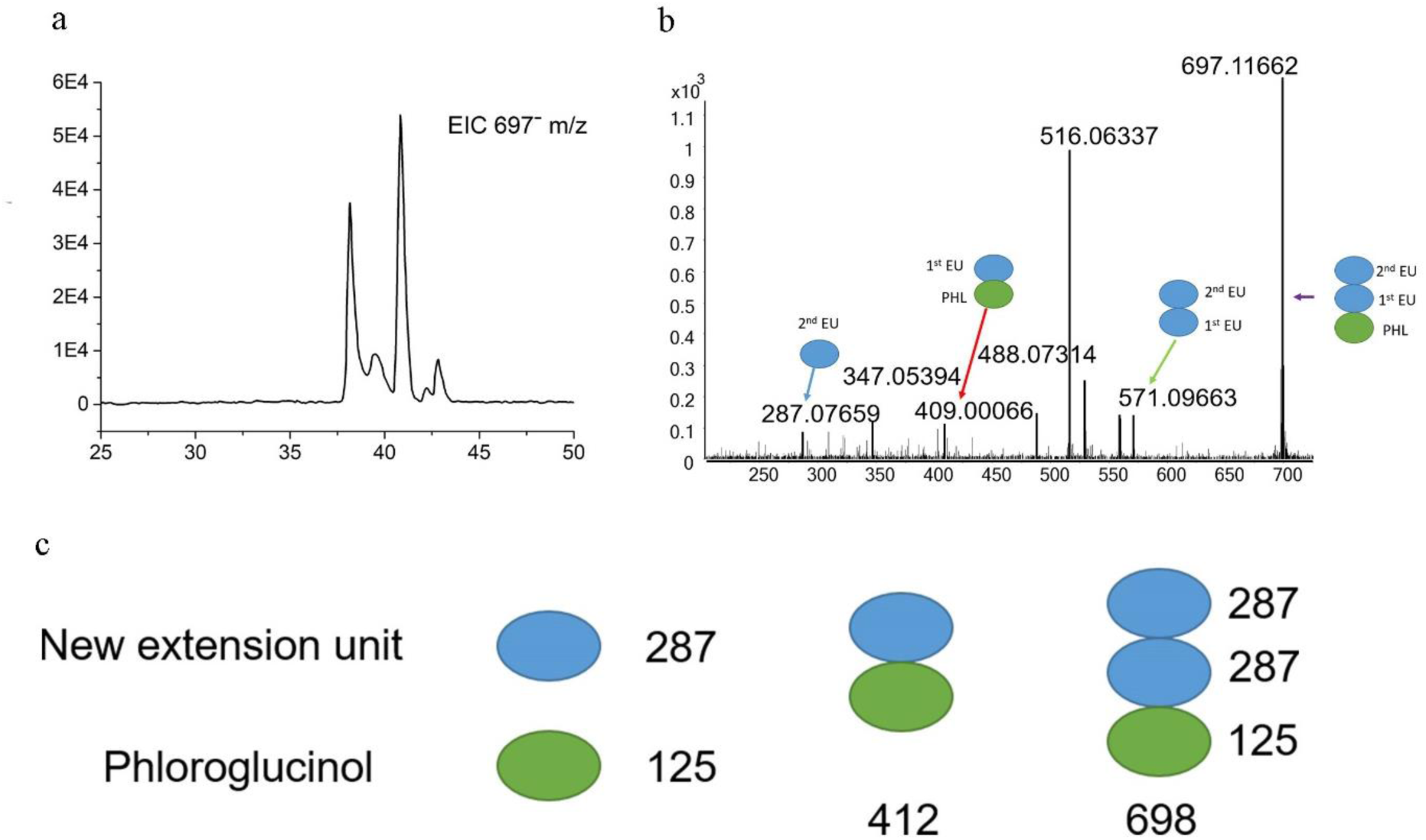
Trimeric compounds formed from phloroglucinol added in the incubation of recombinant FP and epicatechin. Extracted-ion chromatogram (EIC) was performed to characterize MS of compounds formed in the incubations consisting of FP, epicatechin, hydrogen peroxide, and phloroglucinol. **a,** an EIC profile showed five peaks characterized by a 697 [m/z]^-^ produced from the incubations. **b,** MS/MS fragmentation showed main fragments from all 697 [m/z]^-^ compounds, including 571, 516, 409, and 287. **c,** a carton diagram presents a mechanism by which the incubation of FP, epicatechin, hydrogen peroxide, and phloroglucinol produces trimeric compounds. Phloroglucinol forms the starter unit.

**Fig. S33.**
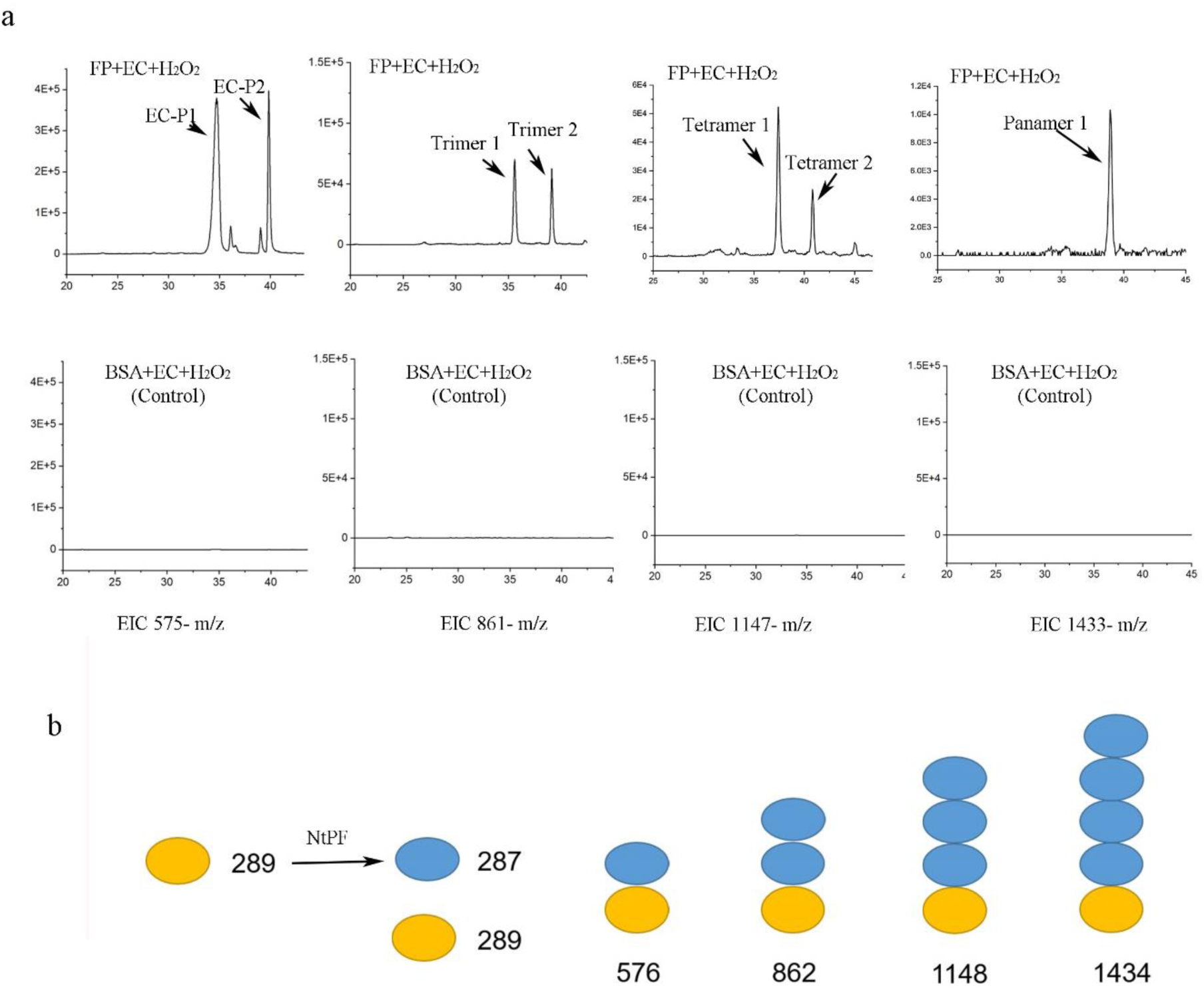
A series of new oligomeric compounds detected from the incubations of FP and epicatechin. Extracted ion chromatograms (EIC) were recorded to show compounds formed in the incubations consisting of FP, epicatechin, and hydrogen peroxide (FP+EC+H_2_O_2_). **a,** EIC showed 575, 861, 1147, and 1433 [m/z]^-^ corresponding to dimers (EC-P1 and EC-P2) timers, tetramers, and pentemers formed from enzymatic reactions (FP+EC+H_2_O_2_) but not from control incubations (BSA+EC+H_2_O_2_). **b,** a carton diagram shows a model by which the incubations form oligomers from the enzymatic reactions of epicatechin and FP.

**Figs. S34-S36 support the prevalence of compounds in the plant kingdom (Figure 6)**

**Fig. S34.**
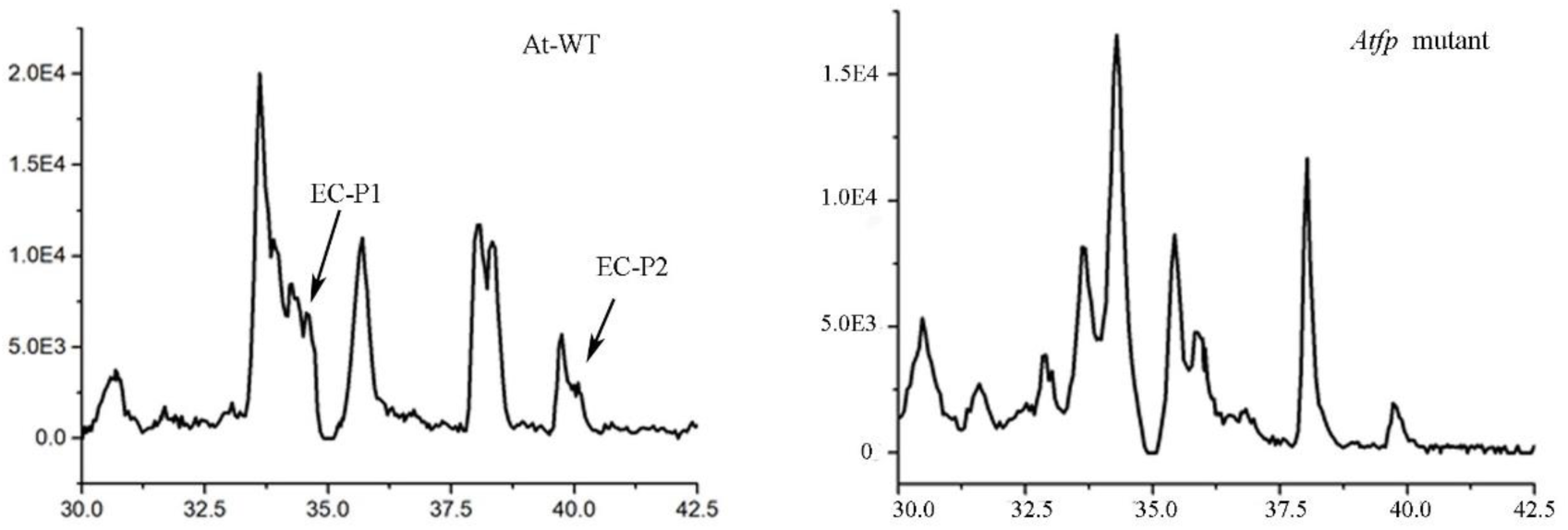
Snapshots showing EC-P1 and EC-P2 in seeds of wild-type plants but not in seeds of *atfp* mutants (Supporting Fig. 6a)

**Fig. S35.**
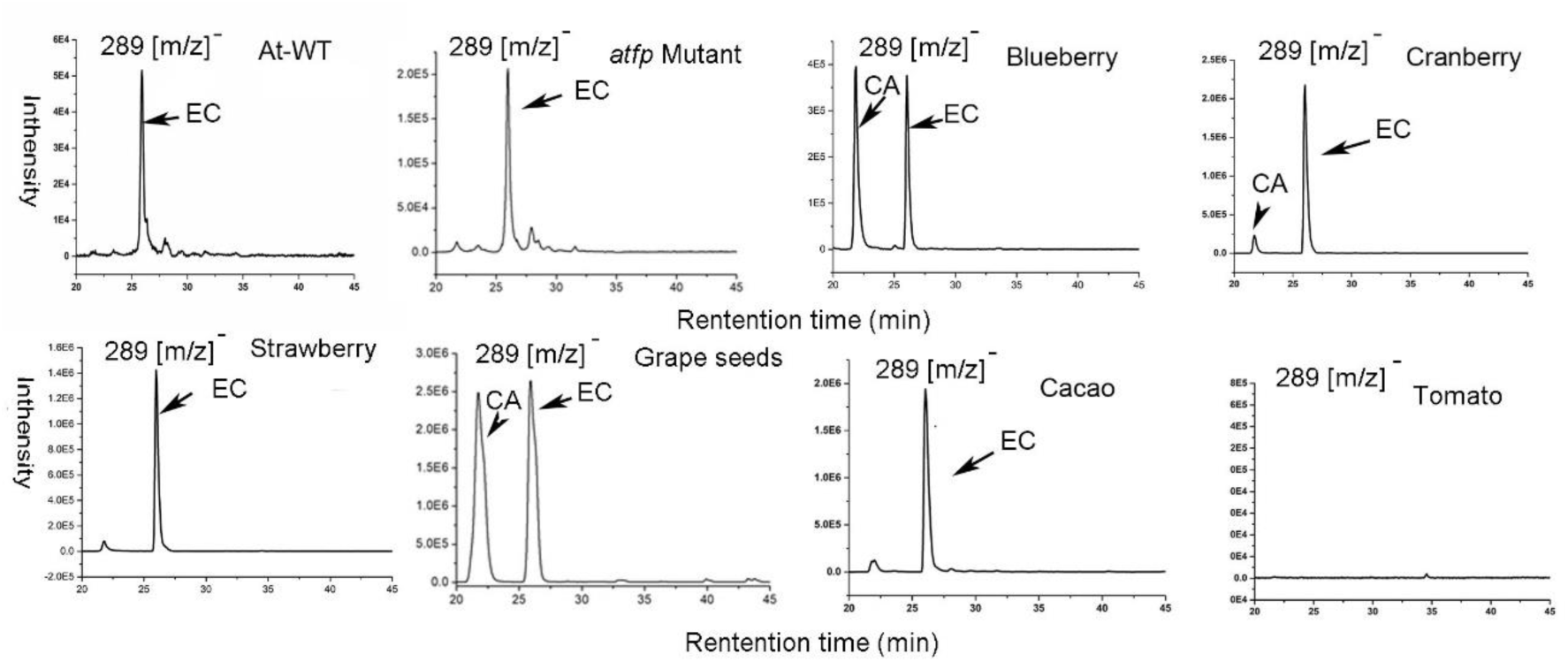
Analysis of epicatechin and catechin in tissues of eight plants. EIC profiling detected epicatechin (EC) in seeds of *A. thaliana* and *atfp* mutants, berry samples of blueberry, cranberry, and strawberry, seed simples of grape and cacao. Catechin (CA) was detected from blueberry, cranberry, and grape seeds. EC and CA were not detected from tomato.

**Fig. S36.**
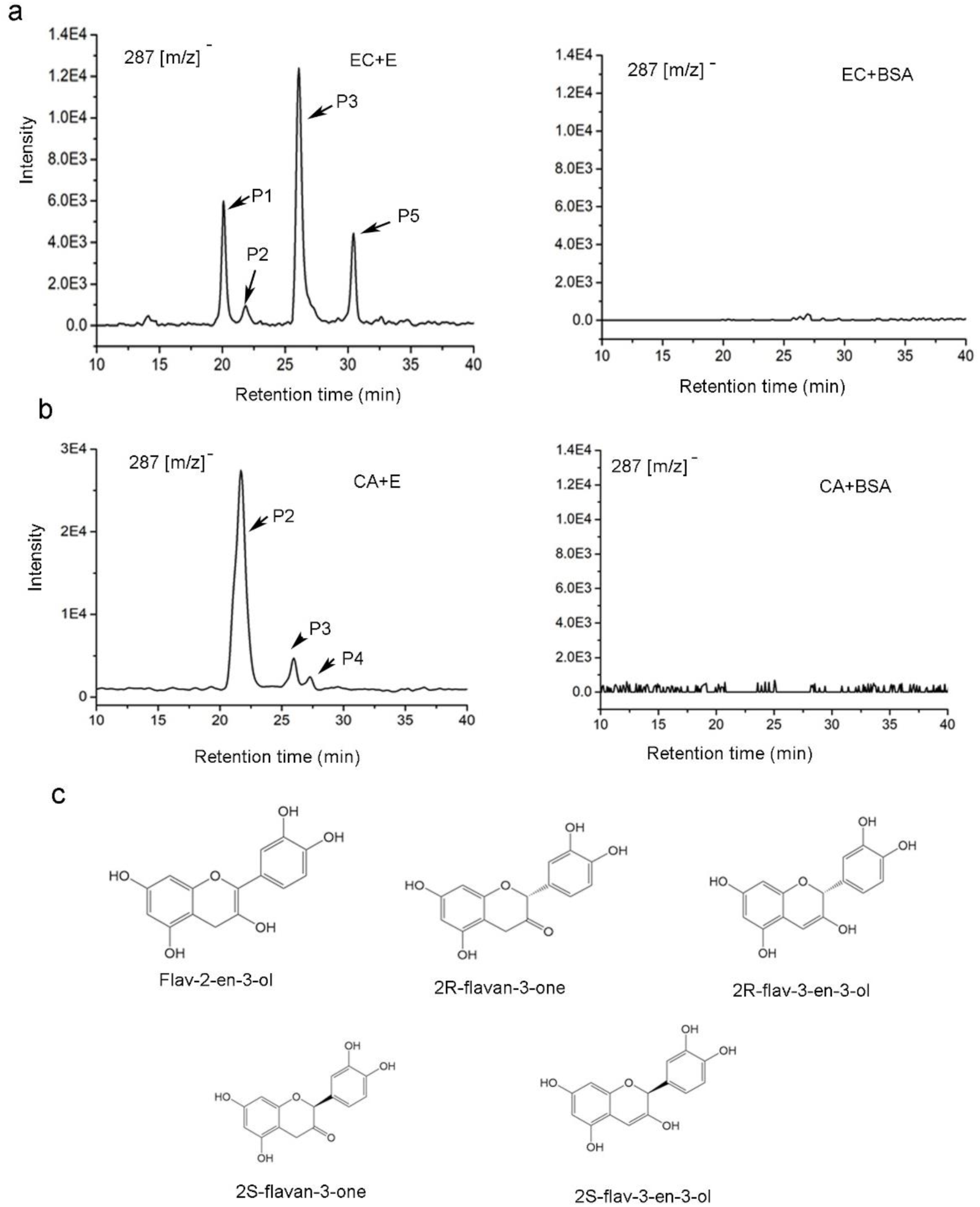
Detection of five 287 [m/z]^-^ compounds from the incubations of FP and epicatechin and catechin. **a,** EIC profiles show that four 287 [m/z]^-^ compounds are formed in the incubations consisting of FP, epicatechin (EC), and hydrogen peroxide, but are not formed in the control incubations consisting of BSA, EC, and hydrogen peroxide. **b,** EIC profiles show that three 287 [m/z]^-^ compounds are formed in the incubations consisting of FP, catechin (CA), and hydrogen peroxide, but are not formed in the control incubations consisting of BSA, CA, and hydrogen peroxide. **c,** five structures are predicted to be flav-2-en-3-ol, 2R-flavan-3-one, 2S-flavan-3-one, 2R-flav-3-en-3-ol, and 2S-flav-3-en-3-ol, which have a molecular weight 288 Dalton.

**Table S1.**
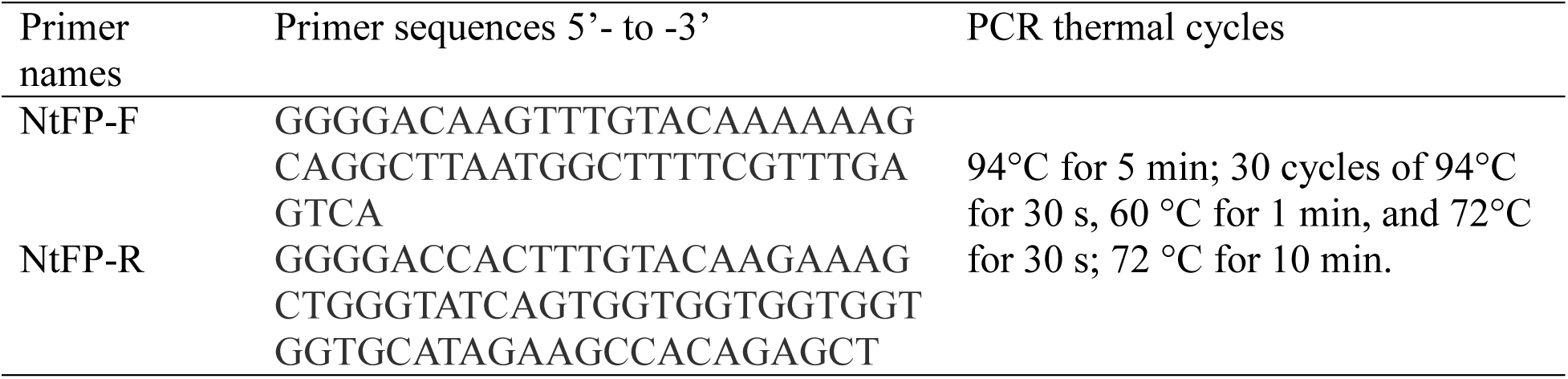
PCR primers and PCR thermal cycles

**Table S2.** The correlations of 26 PB2 protons and of 24 EC-P1 protons are established from 2D ROESY NMR tests

| Position<br>(Proton) | Correlations (Proton -Proton) |  |  |  |
| --- | --- | --- | --- | --- |
|  | PB2 |  | EC-P1 |  |
|  | PPM | Correlations | PPM | Correlations |
| A7OH | 8.57 | A7OH-A6H, A7OH-A8H | 8.18 | A7OH-A6H, A7OH-A8H |
| E7OH | 8.55 | E7OH-E6H | 8.39 | E7OH-C4H, E7OH-E6H |
| F3'OH | 8.26 | F3'OH-F2'H | 8.53 | F3'OH-F2'H |
| A5OH | 8.4 | A5OH-C4H, A5OH-A6H | 8.55 | A5OH-A6H |
| B3'OH | 8.26 | B3'OH-B2'H | 8.51 | B3'OH-B2'H |
| F4'OH | 8.15 | F4'OH-F3'OH | 8.36 | F4'OH-F5'H |
| B4'OH | 8.15 | B4'OH-B5'H | 8.28 | B4'OH-B5'/B6' |
| E5OH | 8.45 or<br>6.99 | E5OH-E6H | 8.61 | E5OH-E6H |
| F2'H | 6.98 | F2'H-G3OH, F2'H-G3H, F2'H-G2H, F2'H-F4'OH, F2'H-F3'OH | 7.15 | F2'H-G2H, F2'H-F3'OH |
| B2'H | 6.98 | B2'H-C2H, B2'H-B3'OH, B2'H-C3H, B2'H-C3OH | 7.1 | B2'H-B3'OH |
| F6'H | 6.67 | F6'H-G3H, F6'H-C3H, F6'H-G2H | 6.84 | F6'H-G3H, F6'H-G2H, F6'H-F4'OH |
| F5'H | 6.71 | F5'H-F4'OH | 6.84 | F5'H-G3H, F5'-G2H, F5'H-F4'OH |
| B5'H | 6.71 | B5'H-C2H, B5'H-B4'OH | 6.84 | B5'H-B4'OH |
| B6'H | 6.67 | B6'H-C2H | 6.8 | n/a |
| A6H | 6.03 | A6H-A5OH, A6H-A7OH | 5.81 | A6H-A5OH, A6H-A7OH |
| A8H | 5.98 | A8H-A7OH | 5.55 | A8H-A7OH |
| E6H | 5.92 | E6H-E5OH, E6H-E7OH | 5.92 | E6H-E5OH, E6H-E7OH |
| C2H | 5.07 | C2H-C3H, C2H-B6'H, C2H-B2'H | n/a | n/a |
| G2H | 5.04 | G2H-G3H, G2H, F6'H, G2H-F2'H | 5.18 | G2H-F5'H, G2H-F2'H |
| C4H | 4.71 | C4H-C3H, C4H-C3OH, C4H-F6'H, C4H-E5OH, C4H-F2'H, C4H-A5OH | 5.41 | C4H-E7OH, C4H-F4'OH |
| G3H | 3.93 | G3H-G4aH, G3H-G4bH, G3H-G2H, G3H-F6'H, G3H-F2'H | **4.65** | **G3H-G4aH, G3H-G4bH, G3H-G2H** |
| C3OH | 4.2 | C3OH-C4H | 6.05 | n/a |
| G3OH | 4.17 | n/a | 4.53 | n/a |
| C3H | 3.93 | C3H-C4H, C3H-C2H, C3H-B6'H, C3H-B2'H | n/a | n/a |
| G4aH, G4bH | 2.92, 2.69 | G4aH-G4bH, G4bH-G4aH, G4aH-G3H, G4aH-G2H | 2.67, 2.78 | G4aH-G4bH, G4bH-G4aH, G4aH-G3H, G4bH-G3H, G4aH-G2H, G4bH-G2H |

**Table S3.**
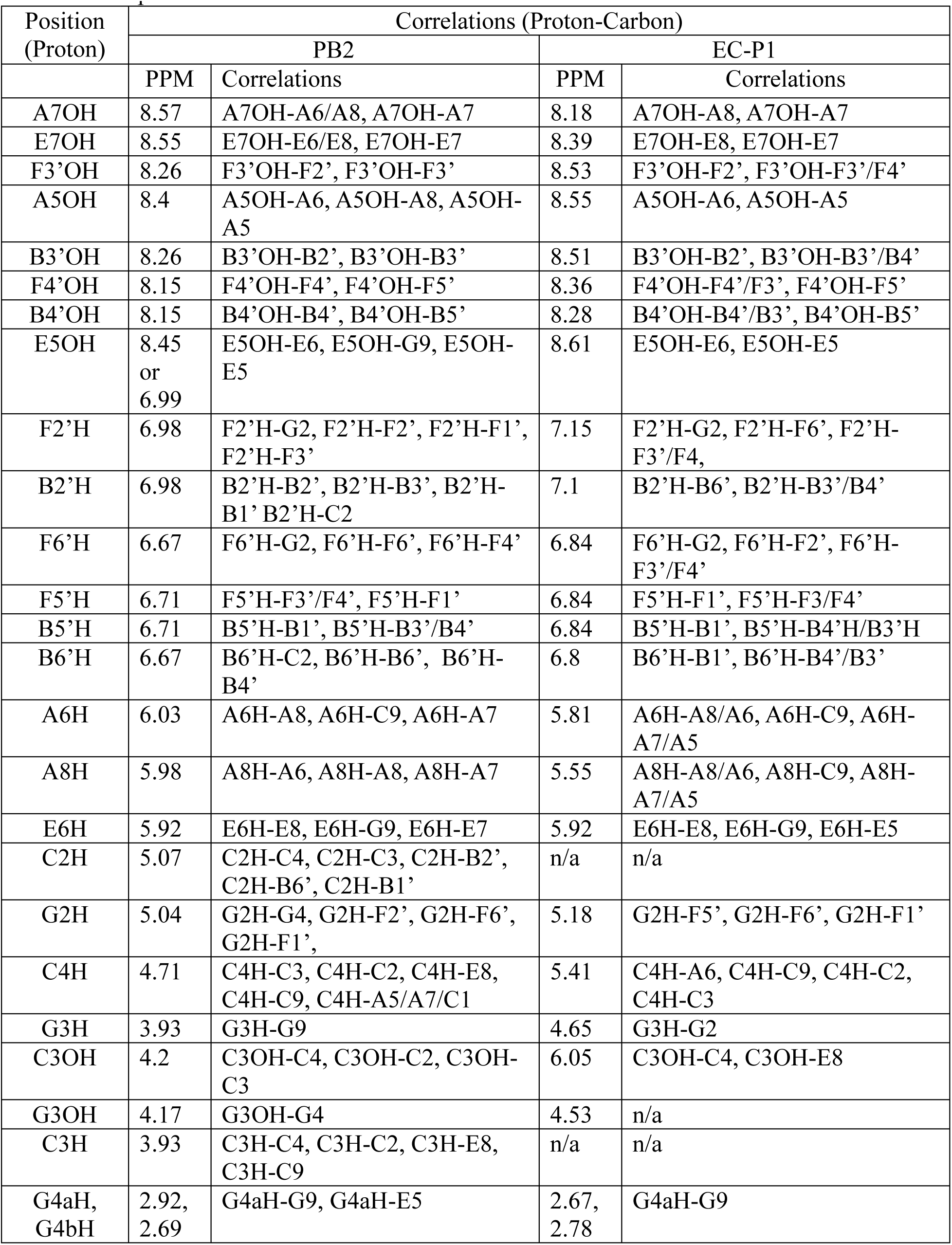
The proton-carbon correlations of PB2 and EC-P1 are established from HMBC test.

